# How to Approach Machine Learning-based Prediction of Drug/Compound-Target Interactions

**DOI:** 10.1101/2022.05.01.490207

**Authors:** Heval Atas, Tunca Doğan

## Abstract

The identification of drug/compound-target interactions (DTIs) constitutes the basis of drug discovery, for which computational predictive approaches have been applied. As a relatively new data-driven paradigm, proteochemometric (PCM) modeling utilizes both protein and compound properties as a pair at the input level and processes them via statistical/machine learning. The representation of input samples (i.e., proteins and their ligands) in the form of quantitative feature vectors is crucial for the extraction of interaction-related properties during the artificial learning and subsequent prediction of DTIs. Lately, the representation learning approach, in which input samples are automatically featurized via training and applying a machine/deep learning model, has been utilized in biomedical sciences. In this study, we performed a comprehensive investigation of different computational approaches/techniques for protein featurization (including both conventional approaches and the novel learned embeddings), data preparation and exploration, machine learning-based modeling, and performance evaluation with the aim of achieving better data representations and more successful learning in DTI prediction. For this, we first constructed realistic and challenging benchmark datasets on small, medium, and large scales to be used as reliable gold standards for specific DTI modeling tasks. We developed and applied a network analysis-based splitting strategy to divide datasets into structurally different training and test folds. Using these datasets together with various featurization methods, we trained and tested DTI prediction models and evaluated their performance from different angles. Our main findings can be summarized under 3 items: (i) random splitting of datasets into train and test folds lead to near-complete data memorization and produce highly over-optimistic results, as a result, it should be avoided, (ii) learned protein sequence embeddings work well in DTI prediction and offer high potential, even though no information related to protein structures, interactions or biochemical properties is utilized during their generation, and (iii) PCM models tend to learn from compound features and leave out protein features, mostly due to the natural bias in DTI data, indicating the requirement for new and unbiased datasets. We hope this study will aid researchers in designing robust and high-performing data-driven DTI prediction systems that have real-world translational value in drug discovery.

## 1. Introduction

Drug discovery is a long-term and costly process that involves the identification of bioactive compounds as drug candidates via screening experiments. Although the advancements in high-throughput screening technology allow the scanning of thousands of compounds simultaneously, it is still not possible to fully analyze a certain portion of the target and compound spaces due to the excessive number of possible protein-compound combinations. This situation led to the emergence of computational approaches, such as the virtual screening (VS), for the *in silico* prediction of unknown drug-target interactions (DTIs) to aid screening experiments ^1^. Conventional ligand-based (e.g., QSAR modelling) and structure-based (e.g., molecular docking) VS approaches aim to predict interactions between a set of compounds and a predefined target protein. Ligand-based approaches mainly achieve this by utilizing molecular property-based compound similarities ^2^, while structure-based approaches employ 3-D structures of targets and compounds ^3^.

Until recently, most machine learning-based computational drug discovery studies approached the subject of predicting physical interactions between drug candidate compounds and target proteins from a (ligand-based) chemo-centric point of view, only utilizing compound attributes/properties. These studies ignore protein features by treating target proteins just as labels for input compounds. As a rather novel approach in this area, proteochemometric (PCM) modelling aims to predict bioactivities by incorporating both compound and target features, usually via readily available molecular notation (e.g., SMILES) and amino acid sequence data, without requiring hard to obtain 3-D structures and dynamic information ^4^. PCM can predict bioactivity relationships between large sets of compounds and targets under a single system using statistical/data-driven modeling techniques such as machine learning. This characteristic of PCM also allows the identification of off-target affects -a significant limitation of conventional VS approaches ^4, 5^-, which is especially important for drug repurposing and side-effect identification. To construct a machine learning-based PCM model, first, input compounds and/or target proteins are converted into quantitative feature vectors, so called “representations”, based on their molecular properties (i.e., descriptors). These vectors are then processed together with the magnitude of the bioactivity/interaction between these compounds and targets (i.e., labels), via machine learning algorithms, during the process of supervised model training ^6^. For the automated artificial learning of DTIs to be successful, input feature vectors should comprise information about the interaction-related properties of compounds and targets. The better the input data is represented, the better the model can learn and generalize the shared properties among the dataset. Therefore, featurization of the input samples is crucial to construct models with high predictive performance.

Various types of featurization approaches have been used for representing compounds and proteins. Due to the abundance of ligand-based DTI prediction methods, compound representations are extensively studied in the literature ^7–9^. Therefore, this study focuses on protein representation techniques, which is a rapidly developing area lately. Sequence-based protein representations, which utilize amino acid sequences as input, are widely preferred in protein associated predictive tasks since 3-D structural information is not available for many proteins and/or proteoforms. Additionally, computational intensity of protein structured-based models is usually high. Considering algorithmic approaches, sequence-based protein representations can be grouped as conventional/classical descriptors (or descriptor sets) and learned embeddings. Conventional descriptors are mostly model-driven, meaning that they are generated by applying predefined rules and/or statistical calculations on sequences considering various molecular properties that include physicochemical ^10–12^, geometrical ^13, 14^ and topological ^12^ characteristics of amino acids, as well as sequence composition ^11, 15^, semantic similarities ^16^, functional characteristics/properties ^17–20^, and evolutionary relationships ^13, 21^ of proteins. Learned protein embeddings (a.k.a. representations) are constructed via data-driven approaches that project protein sequences into high-dimensional vector spaces in the form of continuous feature representations using machine/deep learning algorithms. These protein representation learning (PRL) methods usually borrow their data modeling concepts from the field of natural language processing (NLP), where amino acids in a sequence are treated like words in a sentence/document. Due to this reason, many PRL methods are also called “protein language models”. These models usually process raw protein sequences within unsupervised learning, without any prior knowledge about their physical, chemical, or biological attributes ^22^. Even though they are trained solely on the information about the arrangement of amino acids in the sequence, these models are still found to be successful in automatically extracting physicochemical properties ^23^ and functional characteristics of proteins ^24^. PRL methods have a wide range of applications including the prediction of secondary structure ^24–27^, ligand-target protein interaction ^28–30^, splice junction prediction ^31^, family classification ^23^, protein function ^32^, remote homology detection ^27, 33^, and protein engineering/design ^24, 27^.

For evaluating the effectiveness of different types of protein featurization in different areas of protein informatics, carefully designed benchmark studies are required. In contrary to studies that investigate compound featurization, only a few works are available for benchmarking protein representations. These studies mostly focus on tasks such as protein family prediction ^11^, bioactivity modelling ^34, 35^, and predicting biological properties for protein (re)design ^36^. Also, these studies mainly evaluate conventional descriptors rather than novel featurization approaches. As a result, there is an immediate requirement to evaluate cutting-edge protein language models, and comparing them against well-known conventional descriptors in the context of drug-target interactions for drug discovery/repurposing.

PCM modelling has shown promising results when compared to conventional approaches of DTI prediction ^34, 35^; however, it is still far away from conquering this problem. One of the reasons behind this (apart from the topic of featurization) is that the mechanism of learning is not well-understood in PCM, unlike ligand-based modeling. In ligand-based methods, the model predicts new interactors for a target protein based on molecular similarities to its known ligands. In PCM, there are two factors, i.e., the compound features and the protein features, and it is not clear to what degree similarities in-between protein samples and in-between compound samples contribute to the artificial learning of their interactions, and whether there is bias in this process. Another problem associated with data-driven DTI prediction is the reporting of overoptimistic performance results due to; *(i)* low coverage on compound and/or target spaces in training datasets, in terms of molecular and biological properties (i.e., limited variance), which prevents models from gaining the ability to generalize, and *(ii)* poorly planned and applied train/test dataset preparation (e.g., splitting data randomly) and model evaluation strategies. Most of the self-proclaimed high performing DTI prediction models in the literature are not translated well into real-world cases due to these non-realistic assessments. Recently, there have been efforts in terms of applying different dataset splitting strategies including temporal splitting ^37^, non-overlapped sampling ^9, 38^, cluster-cross-validation ^39^, and scaffold-based splitting ^40^ to build robust models. Temporal splitting strategy only considers a time-dependent data point separation. In non-overlapped sampling strategy, three different settings are applied: warm start (common drugs and targets are present in both the training and test sets), cold start for drugs (drugs in the training set are unseen in the test set while common proteins are shared in these sets), cold start for proteins (proteins in the training set are not involved in the test set, but common drugs are allowed to be present in both sets) ^41^. This strategy only differentiates samples in terms of identity, and does not take similarities between compound and/or proteins into account. Although cluster-cross-validation and scaffold-based splitting methods prevent the involvement of similar compounds in train and test sets, they do not take target protein similarities into consideration. These strategies are not sufficient for evaluating PCM-based DTI prediction models, in which there are three types of relationships to account for; *(i)* compound-target protein interactions, *(ii)* compound-compound similarities, and *(iii)* protein-protein similarities.

New computational approaches, evaluation strategies and datasets are required in order to address the aforementioned issues in data-centric evaluation and prediction of DTIs. With the aim of contributing to the field of data-driven bioactivity modelling for drug discovery and repurposing, here, we performed a rigorous benchmarking study. One of the goals in this study is to identify feature types with better representation capabilities to be used in the automated prediction of DTIs. To achieve this, we built prediction models for various sequence-based protein representations. We employed widely used conventional protein descriptors by selecting those that reflect different molecular aspects of proteins. We also utilized the state-of-the-art protein representation learning methods (i.e., protein language models). Another goal of this study is the preparation of new challenging benchmark datasets with high coverage on both compound and protein spaces, which can also be utilized in future studies. We carefully prepared small-, medium- and large-scale datasets by applying extensive filtering operations and a network-based splitting strategy to acquire realistic and well-balanced datasets. To our knowledge, this data splitting strategy which considers 3 types of relationships (i.e., drug-target interactions, protein-protein similarities, and compound-compound similarities), is proposed here for the first time. We used these datasets in our protein representation benchmarks. In this study, we also evaluated different forms of; (i) DTI modeling techniques, (ii) preliminary and explanatory data exploration approaches, and (iii) model performance evaluation and comparison strategies.

The study is summarized in a schematic workflow in Figure 1. Firstly, we prepared benchmark DTI prediction datasets by applying filters specific to each data scale and explored them via different data visualization techniques. We then split these datasets into train and test folds using different strategies to reflect the real-world data-centric challenges in drug discovery. For the construction of machine learning models, we implemented target feature-based and PCM modelling approaches, and trained/tested our models under various conditions. All details regarding the construction of datasets, representations and DTI prediction models are provided in the Methods section. In the Results and Discussion section, we evaluated the effectiveness of each protein featurization technique on different benchmarks and modeling approaches, and discussed their strengths and weaknesses in comparison to each other. We shared our datasets, results and source code in a re-usable form under the “ProtBENCH” platform at https://github.com/HUBioDataLab/ProtBENCH.

**Figure 1.**
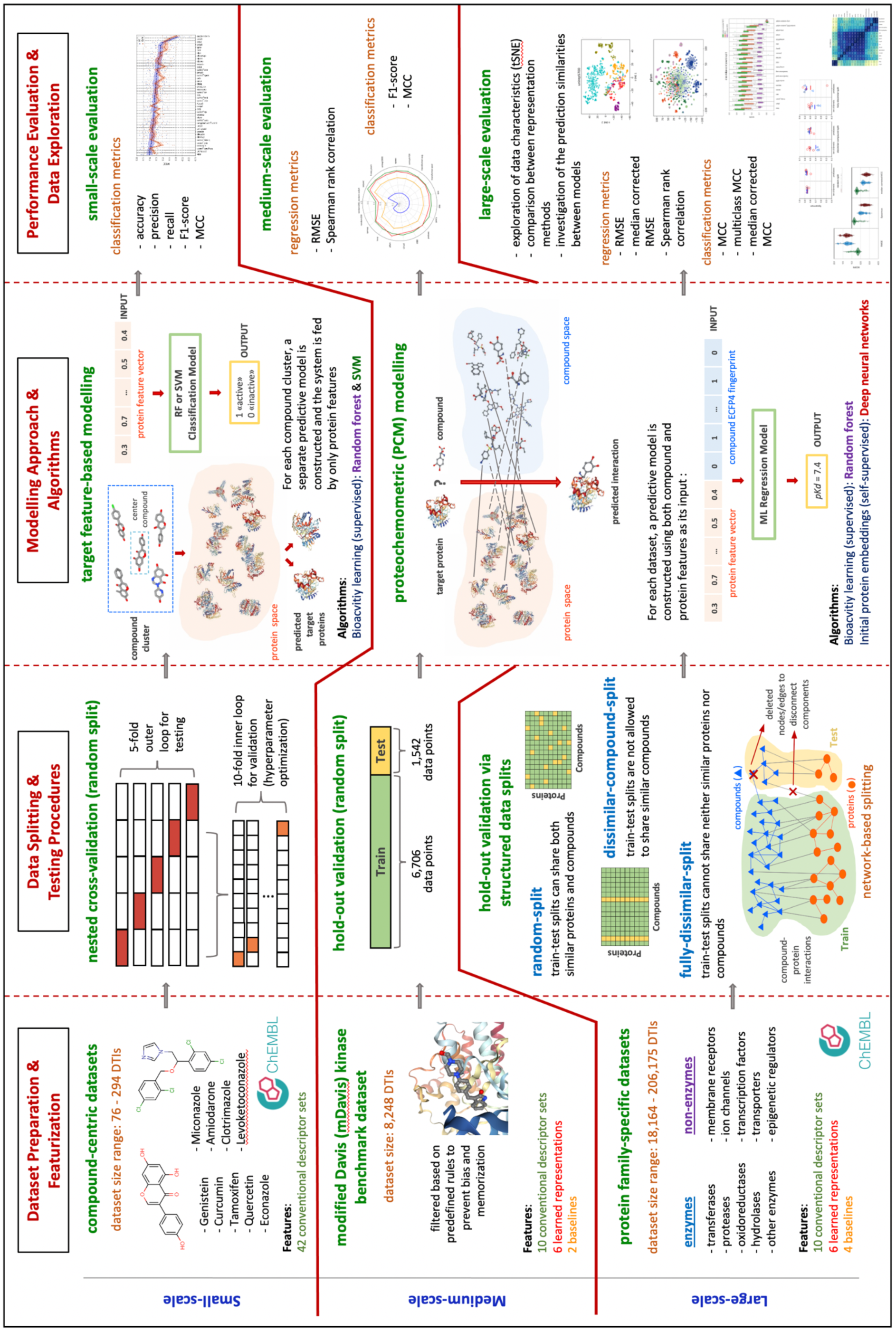
The schematic overview of the study.

As the first comprehensive benchmark study including both conventional and novel protein representation methods in the context of drug discovery and repurposing, we hope this work will aid researchers in choosing suitable approaches and techniques according to their specific modeling tasks. Furthermore, our newly constructed challenging benchmark datasets can be used as reliable, reference/gold-standard datasets in further studies to design robust DTI prediction models with real-world translational value.

## 2. Methods

In this section, we first explain the construction of benchmark datasets, with emphasis on the train/test data splitting strategies. Next, we explain featurization techniques used for representing of proteins and compounds. Then, we summarize modelling approaches and algorithms employed for DTI prediction, along with additional explorative analysis such as the t-SNE projections. Finally, we mention performance evaluation metrics and the tools/libraries we employed.

### 2.1. Dataset Construction and Splitting

In machine learning applications, two significant factors that affect the generalization capability of models are the dataset content/size and the approach used in splitting data points to train/validation/test folds. We constructed and used three groups of datasets at different scales (i.e., small, medium, and large), each of which have distinctive characteristics.

#### 2.1.1. Small-scale: Compound-centric datasets

Here, the aim is to construct datasets of target proteins to be used in DTI prediction models, in which the only input is target feature vectors, and the task is to classify them to their correct ligands. Each dataset is composed of targets of a specific drug/compound as reported in the ChEMBL (v24) database ^42^ considering experimentally measured bioactivities. Bioactivity data points with pChEMBL values, i.e., -log(IC50/EC50/Ki/Kd/Potency, …), greater than 5 (equivalent to IC50/EC50/Ki/Kd/Potency <10 uM) are placed in the positives (actives) dataset, and instances with pChEMBL <= 5 are placed in the negatives (inactives) set. In most cases, sizes of these compound centric training datasets were too small to construct robust prediction models. In order to overcome this problem, we first selected compounds with the highest number of active and inactive bioactivity data points, which we called “center compounds’’. Afterward, we constructed compound clusters around these center compounds by calculating pairwise molecular similarities between each center compound and all other compounds in the ChEMBL database using ECFP4 fingerprints and the Tanimoto coefficient. Compounds that are similar to a center compound with Tanimoto similarity >= 0.3 (as also used in previous studies such as ^43^) are placed in the cluster of the corresponding center compound and their bioactivity data (i.e., active and inactive targets) are incorporated into the cluster’s bioactivity dataset. Therefore, nine independent compound centric, single task classification datasets (with center compounds of Curcumin, Tamoxifen, Quercetin, Genistein, Econazole, Levoketoconazole, Amiodarone, Miconazole, Clotrimazole) were constructed, and their dataset sizes (i.e., the number of targets) range from 76 to 294. Statistics of these datasets, including cluster sizes, active and inactive number of targets, are summarized in electronic supplementary information (ESI) Table S1. In order to balance the number of active and inactive targets in each dataset (initially, the number of inactive targets were considerably low), new proteins which are less than 50% similar to positive targets and less than 80% similar to negative targets already existing in the dataset were selected from ChEMBL and added to the negatives dataset. Due to the small size of datasets, separating a hold-out test fold was not feasible. Therefore, a nested cross-validation approach (with 10-fold inner loop in validation and 5-fold outer loop in testing) was applied during model evaluation. These datasets are used in the small-scale target feature-based analysis described in section 3.2.

#### 2.1.2. Medium-scale: mDavis kinase dataset

We employed the previously proposed Davis kinase dataset ^44^ for performing benchmark analysis on medium-scale, which is a commonly used benchmark for regression-based DTI prediction. The original train-test instances in the Davis dataset are taken from the study by Ozturk et al. ^45^. This dataset includes ∼30,000 DTI data points (real-valued bioactivities); however, the activity values of ∼20,000 of them are recorded as 10 uM (i.e., pKd = 5). These are the data points correspond to cases in which an activity was not observed when the maximum dose of 10 uM is applied (so the highest dose is incorrectly recorded as the bioactivity value). In order to prevent bias, we removed these instances from both train and test portions of the dataset. For the train portion, three additional filters were applied to avoid data memorization. All bioactivities of a compound or target are discarded if the compound or target:

1. only contains active or inactive data points based on the threshold pKd = 6.2, which is the median bioactivity value of the dataset,
2. has an active-to-inactive ratio > 4 or < 1/4 considering its bioactivity data points,
3. has a bioactivity distribution with standard deviation < 0.3, which means bioactivity values vary within a narrow range.

A successful machine learning model is expected to learn general principles from data rather than memorizing it. The instances fulfilling the conditions above may not contribute to the learning process, as they can be easily predictable regardless of the algorithm or feature set, since they have very similar outcomes. We removed these instances from the dataset; otherwise, the model would perform well just by memorizing the outcome of these cases. After these filtering operations, the finalized set, which we call the modified Davis (mDavis) dataset, contains 6,706 train and 1,542 test data points. This dataset is used in the medium-scale PCM-based analysis described in section 3.3.

#### 2.1.3. Large-scale: Protein family-specific datasets

With the aim of constructing large-scale gold standard datasets, we applied rigorous filtering operations on the recorded bioactivities of target proteins from different protein families including membrane receptors, ion channels, transporters, transcription factors, epigenetic regulators, and enzymes with five subgroups (i.e., transferases, proteases, hydrolases, oxidoreductases, and other enzymes). Protein family information is taken from th ChEMBL ^42^ target protein classification. We excluded classes such as secreted proteins, other categories, and unclassified proteins which have inadequate number of bioactivity data points. Here, we actually mean protein super families; however, these terms are used in different (but related) contexts in various resources, as a result, we use the term “family” throughout the article for convenience.

For enzymes, subclasses belonging to the same main class were merged based on their EC number annotations. The merged enzyme classes and their corresponding EC numbers are given in ESI Table S2. Bioactivity data of these families are retrieved from the ChEMBL (v24) database. Bioactivity data points that satisfy the following criteria, target type: “single protein”, standard relation: “=”, pChEMBL value: “not null”, and assay type: “B” (binding assay) are included in the dataset and the rest are discarded. Dataset statistics of each protein family are provided in ESI Table S3.

For each protein family-based dataset, three types of train-test folds were extracted based on different dataset splitting strategies based on molecular similarities in-between compounds and proteins. For this, we binarized pairwise similarity measurements as “similar” or “non-similar”. UniRef50 clusters ^46^ were used for generating protein similarity matrices, where proteins in the same cluster were accepted as similar to each other (equivalent to a threshold of 50% sequence similarity). Otherwise, proteins were considered dissimilar to each other. For compounds, Tanimoto coefficient-based pairwise similarities were calculated using compound ECFP4 fingerprints and the RDKit library ^47^. Compound pairs with a Tanimoto score >= 0.5 were accepted as similar to each other. Otherwise, compounds were considered dissimilar to each other.

##### Random-split dataset

This dataset is constructed by applying a complete random splitting strategy, so that similar compounds and proteins are presented in both train and test sets. Random splitting is one of the most widely used dataset split strategies in machine learning applications; however, it eases the prediction task due to the sharing of highly similar instances between train and test sets. Thus, models usually display overoptimistic performance results. In our random-split protein family-specific datasets, at least 95% of proteins and 60% of compounds in test sets are found to be similar to the ones in their respective train sets.

##### Dissimilar-compound-split dataset

This dataset is constructed by applying a strategy that only considers compound similarities while distributing bioactivity data points into train-test splits. Compounds in train and test splits are dissimilar to each other (Tanimoto score < 0.5). Therefore, similar compounds are not allowed to take part in both train and test splits. This strategy makes the prediction task more difficult and realistic compared to random splitting and partly prevents the model from memorizing bioactivities over identical or highly similar compound fingerprints shared between train and test folds.

##### Fully-dissimilar-split dataset

The aim here is to create train test folds in a way that neither compounds nor proteins are similar to each other between train and test. This dataset is constructed using a network-based splitting strategy to separate bioactivity data points (i.e., compound-target pairs) into disconnected components. Later, each component is either used in training or test splits. Actually, this dataset is extremely challenging for any DTI prediction method. However, this approach is crucial to evaluate a DTI prediction model’s ability to accurately predict new targets and/or new ligands that are truly novel (i.e., there is no bioactivity information for these compounds and target proteins in source databases, moreover, there are no compounds and target proteins significantly similar to these compounds and targets in source bioactivity databases), as this is one of the most crucial expectations from the PCM modeling approach. The steps of the network-based splitting process are provided below:

*1)* Protein-protein and compound-compound pairwise similarity matrices were constructed independently for each protein family, based on protein family membership information and interacting compounds for those proteins (obtained from ChEMBL bioactivity data points). Similarity values were binarized according to the procedure explained above (i.e., 50% sequence/molecular similarity threshold for both protein and compounds).
*2)* A heterogeneous network was constructed for each protein family by merging similarity matrices and bioactivity data using the NetworkX Python library ^48^, where nodes represent proteins and compounds, and edges represent protein-protein or compound-compound similarities, and compound-protein interaction (bioactivity) relationships. It is ensured that any two components that are disconnected from each other in the network do not share any similarity at all (either directly or indirectly). As a result, all bioactivity data points in a particular component can be placed in the training fold, while the ones in another component can be placed in the test fold. As a result, bioactivity data points (i.e., compound-target pairs) in training and test folds are always guaranteed to be fully dissimilar from each other. In practice, the problem was that nearly all nodes in the network formed a giant connected component, which means that it was not possible to distribute data points to training and test folds over disconnected components.
*3)* In order to overcome this issue, we preferred to discard some of the nodes (e.g., compounds) and edges (e.g., bioactivity data points) from the dataset to subdivide the giant connected components into smaller pieces. Instead of removing nodes and edges randomly, which may cause the loss of a high number of data points, we employed the Louvain heuristic algorithm ^49^ to detect communities in the giant component. This algorithm computes the partition of graph nodes by maximizing the network modularity. By discarding bioactivity edges (or in some cases, discarding nodes if the edge of interest is a similarity-based edge between two compounds) between different communities, the number of disconnected components was increased. Finally, bioactivity data points in each component were assigned either to training or test sets in a way that the ratio of the number of training fold data points to the test fold could be held within reasonable values, which still varied considering different protein families (i.e., from the minimum of 8.70% to a maximum of 23.97%).

Discarded data points of the fully-dissimilar-split dataset were also excluded from training-test folds of random-split and dissimilar-compound-split datasets for keeping instances of three sets exactly the same, to yield fully comparable results. The sizes of these datasets (after discarding data points) range from 18,164 to 206,175 depending on the protein family. Detailed split-based statistics of are provided in ESI Table S4. These datasets are used in the large-scale PCM-based analysis described in section 3.4.

### 2.2. Types of Featurization for Proteins and Compounds

We converted proteins and compounds into fixed-length numerical feature vectors to be used in our DTI prediction models as input samples. The following sub-sections describes different featurization approaches used in this study.

#### 2.2.1. Protein representations

On the basis of sequence-based modeling approaches utilized, we divided this subsection into two categories as conventional protein descriptors and learned protein embeddings. These methods are explained below in terms of their molecular and technical aspects. Names, descriptions, and feature vector dimensions of these descriptors are given in Table 1.

**Table 1.** Properties of the selected protein descriptor sets and representations used in our benchmarks.

| Name | Approach | Description | Dimension |
| --- | --- | --- | --- |
| apaac <sup>10</sup> | Model-driven (physico-chemistry) | Amino acid composition regarding the sequence order correlated factors computed from hydrophobicity and hydrophilicity indices of a.a* | 80 |
| ctdd <sup>50</sup> | Model-driven (physico-chemistry) | Chain length-based distribution of a.a for selected physicochemical properties | 195 |
| ctriad <sup>51</sup> | Model-driven (physico-chemistry) | Triad frequency of residues classified on dipoles and volumes of aa side chains | 343 |
| dde <sup>15</sup> | Model-driven (sequence comp.**) | Dipeptide composition deviation | 400 |
| geary <sup>53</sup> | Model-driven (physico-chemistry) | Autocorrelation regarding the distribution of physicochemical properties of a.a | 240 |
| k-sep pssm <sup>21</sup> | Model-driven (sequence homology) | Column transformation-based position specific scoring matrix (pssm) profiles | 400 |
| pfam <sup>18</sup> | Model-driven (functional properties) | Protein domain profiles | 38-294 *** |
| qso <sup>58</sup> | Model-driven (physico-chemistry) | Sequence order effect based on physicochemical distances between coupled residues | 100 |
| spmap <sup>59</sup> | Model-driven (sequence comp.*) | Subsequence-based feature map | 544 |
| taap <sup>63</sup> | Model-driven (physico-chemistry) | Summation of corresponding residue values for selected physicochemical properties | 10 |
| random 200 | - | Randomly generated continuous numbers between 0 and 1 with uniform distribution | 200 |
| protvec <sup>23</sup> | Data-driven (learned embedding) | Sequence embedding utilizing skip-gram modelling approach | 100 |
| seqvec <sup>25</sup> | Data-driven (learned embedding) | Sequence embedding based on bi-directional language model architecture “ELMo” | 1024 |
| transformer <sup>27</sup> | Data-driven (learned embedding) | Transformer-architecture based embedding method that utilizes attention mechanism | 768 |
| unirep <sup>24</sup> | Data-driven (learned embedding) | Sequence embedding based on mLSTM architecture as a variation of recurrent neural networks | 1900 & 5700 |

\* amino acids, \*\* composition, \*\*\* size varies depending on the dataset, since pfam vectors only include the domains presented in the given protein dataset.

##### Conventional descriptor sets

This category comprises methods that employ model-driven approaches. This is achieved by transforming various molecular properties of proteins, such as sequence composition, evolutionary relationships, functional characteristics, or physicochemical properties of amino acids, into fixed-length numerical feature vectors with the implementation of predefined rules or statistical calculations. Hence, they convert protein sequences into a quantitative and machine-processible format that stores the relevant molecular information. Ten conventional protein descriptor sets used in all 3 of the benchmark analyses of this study are briefly explained below.

**-** *apaac (amphiphilic pseudo amino acid composition)* represents the amino acid composition of protein sequences without losing the residue order effect by using sequence-order factors. These factors are computed from correlation functions of hydrophobicity and hydrophilicity indices of amino acids. Therefore, apaac keeps the distribution of amphiphilic amino acids along the protein chain. It was proposed by Chou in 2005 and used for the prediction of enzyme subfamily classes ^10^.
**-** *ctdd (distribution)* provides distribution patterns of amino acids in terms of the class they belong to considering a particular property. It utilizes 7 types of physicochemical properties including hydrophobicity with 7 different versions, normalized Van der Waals Volume, polarity, polarizability, charge, secondary structures, and solvent accessibility. Each property is divided into 3 classes and 20 amino acids are distributed into these classes based on their values for corresponding property (i.e., helix -EALMQKRH-, strand -VIYCWFT-, and coil - GNPSD-classes for secondary structure property). The distribution patterns are determined according to five different positions (residues) for the corresponding class, which are the first residue, and the residues exactly at the 25%, 50%, 75%, and 100% of the sequence. These positions are divided by the length of the whole protein sequence for the calculation of fractions of each class. This descriptor set was first proposed by Dubchak for protein fold recognition task ^50^.
**-** *ctriad (conjoint triad)* is based on the frequency of triple amino acid combinations in a protein sequence, where amino acids are first converted into a 7-letter reduced alphabet. These seven groups include {A,G,V}, {I,L,F,P}, {Y,M,T,S}, {H,N,Q,W}, {R,K}, {D,E}, and {C}. Amino acid groups are determined according to dipoles and volumes of the amino acid side chains. Ctriad was first used for the prediction of protein-protein interactions by Shen et al. ^51^.
**-** *dde (dipeptide deviation from expected mean)* is a type of sequence composition descriptor set that relies on the deviation of dipeptide compositions from the expected means. Three parameters, i.e., dipeptide composition (*Dc*), theoretical mean (*Tm*), and theoretical variance (*Tv*), are computed for the construction of the dde descriptor set. Saravanan and Gautham proposed it in 2015 for the use of B-cell epitope prediction ^15^.
**-** *geary* utilizes the distribution of structural and physicochemical properties of amino acids along the sequence. It was first developed by Geary in 1954 ^52^ as a measure of spatial autocorrelation that uses the square-difference of property values. Li et al. served it as a protein descriptor set via the PROFEAT web server^53^. Also, Ong et. al. implemented it for the prediction of protein functional families^11^.
**-** *k-sep_pssm (k-separated-bigrams-pssm)* is a column transformation-based descriptor set that computes the bigram transition probabilities between residues in terms of their positional distances from each other, which corresponds to the “k” value. The transition probabilities are calculated from transformations on position-specific scoring matrix (pssm) profiles of proteins. Pssm profiles represent evolutionary conservation of amino acids in a protein sequence, which are derived from multiple sequence alignments of a homolog set of protein sequences. This descriptor set was first proposed in the study of Saini et al. for improving protein fold recognition ^21^. Wang et al. developed the POSSUM tool to calculate a set of PSSM-based descriptors including k-sep_pssm, and they utilized these descriptors for the prediction of bacterial secretion effector proteins ^54^.
**-** *pfam* represents domain profiles of proteins, according to protein domain annotations in the Pfam database ^55^, in the form of binary feature vectors. For each protein, it encodes the presence (1) and absence (0) of a unique list of domains presented in proteins in the corresponding dataset. This descriptor set was employed in the studies of Yamanishi et al. ^18^ and Liu et al. ^56^ with the purpose of predicting drug-target interactions.
**-** *qso (quasi-sequence order)* reflects the indirect effect of the protein sequence order by calculating coupling factors in terms of distances between contiguous residues in the sequence. The distances are determined using different amino acid distance matrices such as the Schneider–Wrede distance matrix ^57^, which is derived from hydrophobicity, hydrophilicity, polarity, and side-chain volume properties of amino acids. It was first utilised by Chou for the prediction of protein subcellular locations ^58^.
**-** *spmap (subsequence profile map)* is a feature space mapping method that represents sequence composition by calculating the distribution of fixed-length protein subsequence (with a length of 5 residues in the default version) clusters in a protein sequence. Subsequence clusters are generated using BLOSUM62 matrix-based similarities of all possible subsequences in the given protein set, extracted by the sliding windows approach. It was proposed by Sarac et al. for functional classification of proteins ^59^. Later, spmap was used for GO term ^60^ and EC number ^61^ prediction. In this study, spmap-based feature vectors were generated using clusters of 5-residue subsequences of ChEMBL target proteins.
**-** *taap (total amino acid properties)* represents the total sum of corresponding residue values in a protein sequence for the selected properties from the AAIndex database ^62^. It was first employed by Gromiha and Suwa for better discrimination of outer membrane proteins ^63^. The properties used in our study are normalized average hydrophobicity scale, average flexibility indices, polarizability parameter, free energy of solution in water, residue accessible surface area in tripeptide, residue volume, steric parameter, relative mutability, hydrophilicity value and the side chain volume.

iFeature stand-alone tool ^64^ was employed for the calculation of apaac, ctdd, ctriad, dde, geary and qso feature vectors. Protein domain annotations were retrieved from the Pfam database ^55^ for the construction of pfam feature vectors. Spmap feature vectors were calculated using our in-house algorithm explained above ^59^. For the construction of k-sep_pssm and taap vectors, POSSUM ^54^ and PROFEAT ^65^ web servers were used, respectively.

##### Learned embeddings

This category comprises protein representations that utilize solely data-driven approaches for the extraction of molecular information from protein sequences. Learned representations are constructed via the process of artificial learning, in which a model is trained on specific unsupervised/self-supervised tasks such as the prediction of the next amino acid in the sequence. For generating such protein representation models, deep neural network-based architectures and design choices that are frequently used in natural language processing (NLP) field are preferred. During the training process, the model takes protein sequences as input, projects them into a high-dimensional vector space, and generate output in the form of fixed-length numerical feature vectors called “embeddings”. These numerical feature vectors can later be used for representing proteins in other predictive tasks (mostly supervised) such as DTI prediction.

Four protein representation learning methods (making 6 models in total, as 2 of these methods have 2 versions each) used in this study are briefly explained below. Names, descriptions, and feature vector dimensions of these embeddings are given in Table 1.

**-** *unirep* is one of the best-known learned protein representations, which was developed in 2019 by Alley et al. using a variation of recurrent neural networks (RNN) called the multiplicative long-/short-term-memory (mLSTM) architecture ^24^. Alley et al. trained the model on approximately 24 million protein amino acid sequences in the UniRef50 clusters of UniProt, with the objective of predicting the next amino acid in these sequences. They evaluated the representation capability of unirep on various tasks including the prediction of protein stability, semantic similarity, secondary structure, evolutionary and functional information. In our study, we constructed both 1900- and 5700-dimensional unirep protein embeddings (obtained by averaging and summing the output embedding of size 1900x3, respectively) for sequences in our datasets and evaluated them as independent representation methods. The unirep model is available at https://github.com/churchlab/UniRep.
**-** *transformer* is a deep architecture that utilizes the attention mechanism in a way to allow the extraction of context without depending on the sequential order information in the input samples ^66^, and it is the current state-of-the-art in the representation learning and generative modelling of natural languages. As part of the “Tasks Assessing Protein Embeddings (TAPE)” study, Rao et al. developed a transformer-based protein representation learning model using the Bidirectional Encoder Representations from Transformers (BERT) algorithm ^27^. This model was trained on approximately 32 million protein sequence fragments taken from the Pfam domain annotation database ^55^, via masked-token prediction. It was also tested on tasks such as secondary structure prediction, contact prediction, remote homology detection, fluorescence landscape prediction, and stability landscape prediction. For each sequence, the model returns two different versions of representation vectors: (i) averaged, and (ii) pooled, both in 768-dimensions. We used both of these embeddings in our study as independent representation methods. TAPE transformer model is accessible at https://github.com/songlab-cal/tape.
**-** *protvec* was developed by Asgari and Mofrad ^23^ as one of the first models used in the construction of learned protein embeddings. It was trained on 546,790 sequences in the UniProtKB/Swiss-Prot database using the skip-gram modelling approach, in which, given a target residue, the model predicts the surrounding amino acids in the sequence. In protvec, protein sequences were embedded into 100-D vectors of 3-gram subsequences (i.e., 3 consecutively located amino acids) as biological words. For characterizing biophysical and biochemical properties of sequences, these 3-grams were analyzed qualitatively and quantitatively in terms of mass, volume, van der Waals volume, polarity, hydrophobicity, and charge. Protein feature extraction capability of protvec was also evaluated in terms of protein family classification and disordered sequence characterization tasks by representing each protein sequence as the summation of 100-D vectors of its 3-grams. 100-D vector representation of protvec can be retrieved from http://dx.doi.org/10.7910/DVN/JMFHTN.
**-** *seqvec* utilizes bi-directional language model architecture of the “Embeddings from Language Models (ELMo)” method for extracting features relevant to per-residue and per-protein prediction tasks. Heinzinger et al. developed the seqvec model by training on approximately 33 million UniRef50 sequences with the goal of predicting the next amino acid in the sequence ^25^. The authors evaluated the success of seqvec on the prediction of secondary structures and regions with intrinsic disorder at the residue level, and subcellular localization prediction at the protein level. 1024-dimensional seqvec protein embeddings can be computed using the seqvec data repository at https://github.com/rostlab/SeqVec.

##### Random feature vectors

We constructed dummy feature vectors (to be used in baseline prediction models) for performance comparison against real representations, with the aim of observing to what degree proteins descriptors are utilized by DTI prediction models. The one for proteins, namely random200, is a descriptor that constructs a feature vector (with the size of 200x1) for each protein sequence containing randomly generated continuous values ranging from 0 to 1 in each dimension. A similar random descriptor has also been constructed for compounds (i.e., 1024x1 sized binary vectors), which is explained below, in section 2.2.2.

#### 2.2.2. Compound representations

We employed the (circular) fingerprinting approach, and used Extended-Connectivity Fingerprints (ECFPs) as feature vectors (representations) of compounds. ECFPs are circular topological fingerprints that are widely used for molecular characterization, similarity searching, and structure-activity relationship modeling. ECFPs represent the presence of particular substructures by considering circular atom neighborhoods within a diameter range ^67^. We constructed 1024-bit ECFP4 fingerprints (corresponding to a radius of 2) using RDKit ^47^, for which compound SMILES notations were used as input. As output, a fixed-length binary fingerprint vector was generated for each compound by applying a hash function on its substructures. For the medium- and large-scale PCM models, we also generated 1024-bit “random compound fingerprints” to be used in dummy (baseline) models for evaluating the effect of compound information on DTI prediction performances. To be able to simulate ECFP4 fingerprints more realistically, we adjusted the frequency of ones and zeros in the vectors to 0.1 and 0.9, respectively (similar to real ECFP4 feature vectors in our datasets) by introducing prior probabilities during vector construction.

### 2.3. Modelling Approaches

In order to evaluate different protein featurization methods on DTI prediction, we utilized 2 different modelling approaches: (i) target feature-based modelling, and (ii) proteochemometric (PCM) modelling. Below, we summarized each approach together with the implementation details.

#### 2.3.1. Target feature-based modelling

In this modelling approach (which is also known as "*in silico* target-fishing” or “reverse-screening based modelling" in the literature), we trained an independent DTI prediction model for each selected drug/compound cluster (please see section 2.1.1 for more information about the dataset). Feature vectors of proteins that are in the positives and negatives dataset of the compound of interest are given to the model as input for training and testing. Here, the model predicts whether a query protein could be the target of the corresponding compound, via binary classification. Hence, the system input is solely composed of protein features, where compounds are just used as labels.

We generated separate models for each protein descriptor set using support vector machine (SVM) and random forests (RF) classifiers, as these are widely used and well-performing machine learning algorithms. The models are implemented with scikit-learn python library ^68^. For SVM models, “rbf” kernel was applied with optimized C and gamma parameters within ranges of [1,10,100] and [0.001,0.01,0.1,1], respectively. RF models were constructed by adjusting the parameters as; number of trees: 200, and the maximum feature number: the square root of the total number of features. Nested cross-validation (with 10-fold inner loop in validation and 5-fold outer loop in testing) was applied for model evaluation. In the end, we trained and tested 1935 RF and 1935 SVM models (i.e., 43 protein descriptor sets -including random200-for 9 different drug/compound clusters, 5-fold outer loop in nested cross validation: 43*9*5).

#### 2.3.2. Proteochemometric (PCM) modelling

We constructed PCM models for both medium-scale and large-scale datasets (please see sections 2.1.2 and 2.1.3 for more information about these datasets). Here, we only used the RF regression algorithm, since we observed that RF models performed better than SVM models in the previous analysis of target feature-based modelling. For parameters, we adjusted the number of trees to 100 and maximum ratio of features to 0.33 (corresponding to one third of the total number of features).

Here, the task is predicting the actual binding affinity (bioactivity) values of the input samples (i.e., compound-target pairs) in terms of pKd/pChEMBL values. We constructed PCM models for 10 conventional protein descriptor sets, including apaac, ctdd, ctriad, dde, geary, k-sep_pssm, pfam, qso, spmap and taap, and 6 learned representations including protvec, seqvec, transformer-avg, transformer-pool, unirep1900, and unirep5700. Since PCM models are based on compound-target pairs, protein representations were concatenated with 1024 bits ECFP4 representations of compounds to construct the finalized input feature vectors.

We generated two baseline models (to be used in both medium-scale and large-scale analysis) by concatenating random200 protein feature vectors with (i) real ECFP4 fingerprints, and (ii) random compound fingerprints, which are named “random200” and “random200_random-ecfp4” models, respectively. Furthermore, we constructed two additional baseline models to be used in the large-scale analysis, in which the protein features are not utilized at all. In the first one, we used the real ECFP4 fingerprint of the compound in the corresponding compound-target pair to represent the pair (called “only-ecfp4”), and in the second one, we used random compound fingerprints to represent input pairs (called “only-random-ecfp4”). Thus, we trained and tested 18 PCM models for the medium-scale analysis using the mDavis kinase dataset (i.e., models built on 10 conventional descriptor sets, 6 learned representations, and 2 baseline models). For the large-scale analysis, we trained and tested models for 20 featurization types (10 conventional descriptor sets + 6 learned embeddings + 4 baseline models) on 10 protein family-specific datasets each having 3 versions of train-test split folds (explained in section 2.1.3 in detail).

Therefore, we constructed 600 PCM models (i.e., 3 splits * 10 families * 20 types of featurization).

### 2.4. t-SNE Projection of Protein Representations on Large-Scale Datasets

t-distributed stochastic neighbor embedding (t-SNE) is a non-linear dimensionality reduction technique that is frequently employed for the visualization of high dimensional datasets ^69^. For exploratory analysis of protein family-specific large-scale datasets, we applied the t-SNE algorithm on the feature vectors of each protein featurization method and colored nodes according to protein (sub)families and train-test fold data points in two different analyses. For the application of t-SNE, we employed the scikit-learn ^68^ manifold module with default parameters (i.e., 2-D embedding space, perplexity=30, and Euclidean distance metric). We investigated different perplexity values in the range of 40 to 100, and decided that the default value performed sufficiently well for all projections.

### 2.5. Performance Evaluation

The performance of target feature-based classification models (in small-scale analysis) was evaluated via accuracy, precision, recall, F1-score, and Matthews Correlation Coefficient (MCC) metrics via nested cross-validation. F1-score is the harmonic mean of precision and recall; thus, it takes both false positive and false negative predictions into account. MCC incorporates all true and false predictions into the equation, and it is preferred over both accuracy and F1-score due to its robustness and reliability, especially in the cases of dataset imbalance ^70^.

The performance of PCM-based regression models (in both medium-scale and large-scale analysis) was evaluated using Root Mean Square Error (RMSE) and Spearman rank correlation (r_s_) metrics over the hold-out test sets. RMSE computes the deviation of predictions from the actual values, and lower RMSE scores indicate better model performance. Spearman correlation evaluates the relationship between the ranks of the predicted and actual values. One of the problems related to regression-based prediction models is that, the distribution of predicted values can have a shifted average (i.e., the rank of predictions is in correlation with the true labels; however, the mean/median prediction value is either higher or lower than the true mean). Value-based performance metrics suffer from this problem and report underestimated scores. In order to handle this problem in the large-scale analysis (where the problem is evident), we calculated an additional version of RMSE via median correction, so that the median value of predictions becomes equal to the median of the true value distribution (i.e., the median corrected RMSE score).

We also evaluated the results of PCM-based regression models on the basis of classification, using F1-score and MCC metrics. To achieve this in the medium-scale analysis (on the mDavis dataset), samples were classified as active (1) or inactive (0) based on an activity cut-off value of pKd = 7 (i.e., 100 nM in terms of Kd) using the RF classification algorithm. For the large-scale analysis over protein family-specific datasets, regression-based prediction results were converted into binary class and multiclass formats, as it was not possible to retrain 600 models due to high computational requirements. For the binary class, median pChEMBL values of the data points in the training datasets were used as threshold values to separate actives and inactives from each other (i.e., compound-target pairs with bioactivity values higher than the median value of the dataset are accepted as actives, and the ones equal to or lower than the median are accepted as inactives). We also calculated corrected version of MCC using the procedure explained above for “median corrected RMSE” score, and similarly called this metric the “median corrected MCC”. For the calculation of multi-class scores, samples were placed into six different classes based on their true pChEMBL values (class1: <5.0, class2: 5.0 - 5.5, class3: 5.5 - 6.0, class4: 6.0 - 6.5, class5: 6.5 - 7.0, and class5: >=7.0) and calculated average MCC scores over all 6 classes. The reason behind using such a variety of performance metrics was to evaluate models from as many different aspects as possible.

The equations for the basic versions of these metrics are given below:

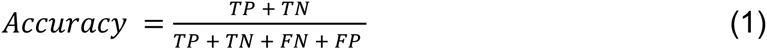

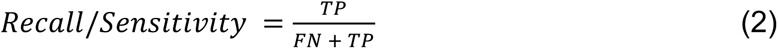

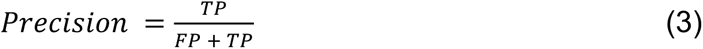

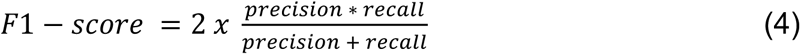

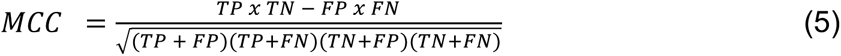

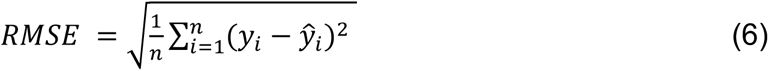

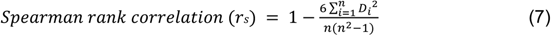

where 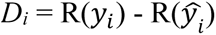; *D_i_* denotes the difference between ranks of true (*y_i_*) and predicted 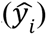 values of samples with the dataset size *n*. TP, TN, FP, and FN represent the total counts of true positive, true negative, false positive, and false negative predictions, respectively.

In this study, we used Python (v3) programming language, scikit-learn library ^68^ for the t-SNE projection and machine learning applications, NetworkX package ^48^ for splitting protein family-specific datasets, RDKit toolkit ^47^ for compound featurization and clustering, POSSUM ^54^ and PROFEAT ^65^ web tools as well as iFeature stand-alone tool ^64^ for protein featurization, and seaborn ^71^ and matplotlib ^72^ libraries for the heatmap analysis and data visualization.

## 3. Results and Discussion

In this section, we evaluate and discuss the results of our benchmark experiments. For this, we first carried out a data exploration analysis via visualizing our data (section 3.1). Next, we trained DTI prediction models under different settings and measured the performance of different protein representations in the small-scale (3.2), medium-scale (3.3), and the large-scale (3.4) analyses. At each section, we discussed model performances from various aspects to address shortcomings in bioactivity modelling studies and contribute to the development of better data-driven models in the field of drug discovery and repurposing.

In this study, we employed random forest (RF) as our main machine learning algorithm (along with SVM in some of the cases) for predicting DTIs. The reasons behind using a classical machine learning algorithm in this benchmark study rather than more complex deep learning-based architectures is that: (i) RF has been used in this field for a long while and shown to work well on numerous occasions, and (ii) deep learning-based complex architectures have already been used in the training stage of learned representations (i.e., protein embeddings); thus, the use of an additional complex architecture in the supervised DTI prediction stage could have prevent the observation of the ability of learned representations in extracting ligand interaction-related properties of proteins, and also, hinder the evaluation of model-driven (i.e., conventional descriptor sets) and data-driven (i.e., learned representations) approaches on a common ground.

### 3.1. Exploration of Data Characteristics

In this subsection, we first visualized members of protein family-specific datasets on 2-D via t-SNE projection. Then, we analyzed split-based characteristics of our datasets by plotting pairwise similarity distributions of proteins and compounds, bioactivity distributions of train-test folds, together with their respective t-SNE embeddings.

#### 3.1.1. t-SNE projection of protein families

For each protein representation, two independent t-SNE projections (one for the enzyme, and another one for the non-enzyme protein families) were carried out (Figure 2a and 2b). Projections for 8 protein featurization methods are shown in Figure 2, and the remaining ones (9 of them) are available in ESI Figure S1. As displayed in these t-SNE plots, generally, protein families are well clustered in both enzyme and non-enzyme projections, with slightly less apparent clusters in enzymes, probably due to the sharing of enzyme-specific properties between proteins. Also, members of the other-enzymes class are scattered among other clusters as its members do not have distinctive characteristics. Although the majority of protein representations are successful at separating at least some of the families, projections of learned embeddings have clearer clusters in general, which indicates their ability of extracting family-specific features. Considering conventional descriptor sets, homology (i.e., k-sep_pssm) and domain profiles (i.e., pfam) are observed to have more distinctive abilities for the classification of protein families, compared to physicochemistry (e.g., apaac, ctdd, ctriad, geary, qso) and sequence composition (i.e., dde). The t-SNE projection of spmap, being a sequence composition-based descriptor set based on protein subsequence (5-mer) clusters, is similar to the projection of random200 descriptor set. This result indicates that 5-residue subsequences of proteins cannot capture family-specific patterns. Highly distinct from other representations, taap has a projection in the form of an S-shaped curve. Feature vectors of proteins with similar residue content and sequence length are similar to each other (independent from the actual order of amino acids on the sequence) according to the taap descriptor set, since taap uses the total sum of the amino acid-based property values to represent a protein. Due to the fact that t-SNE aims to preserve local neighborhoods, proteins form a continuous curve similar to time-series data when represented by taap.

**Figure 2.**
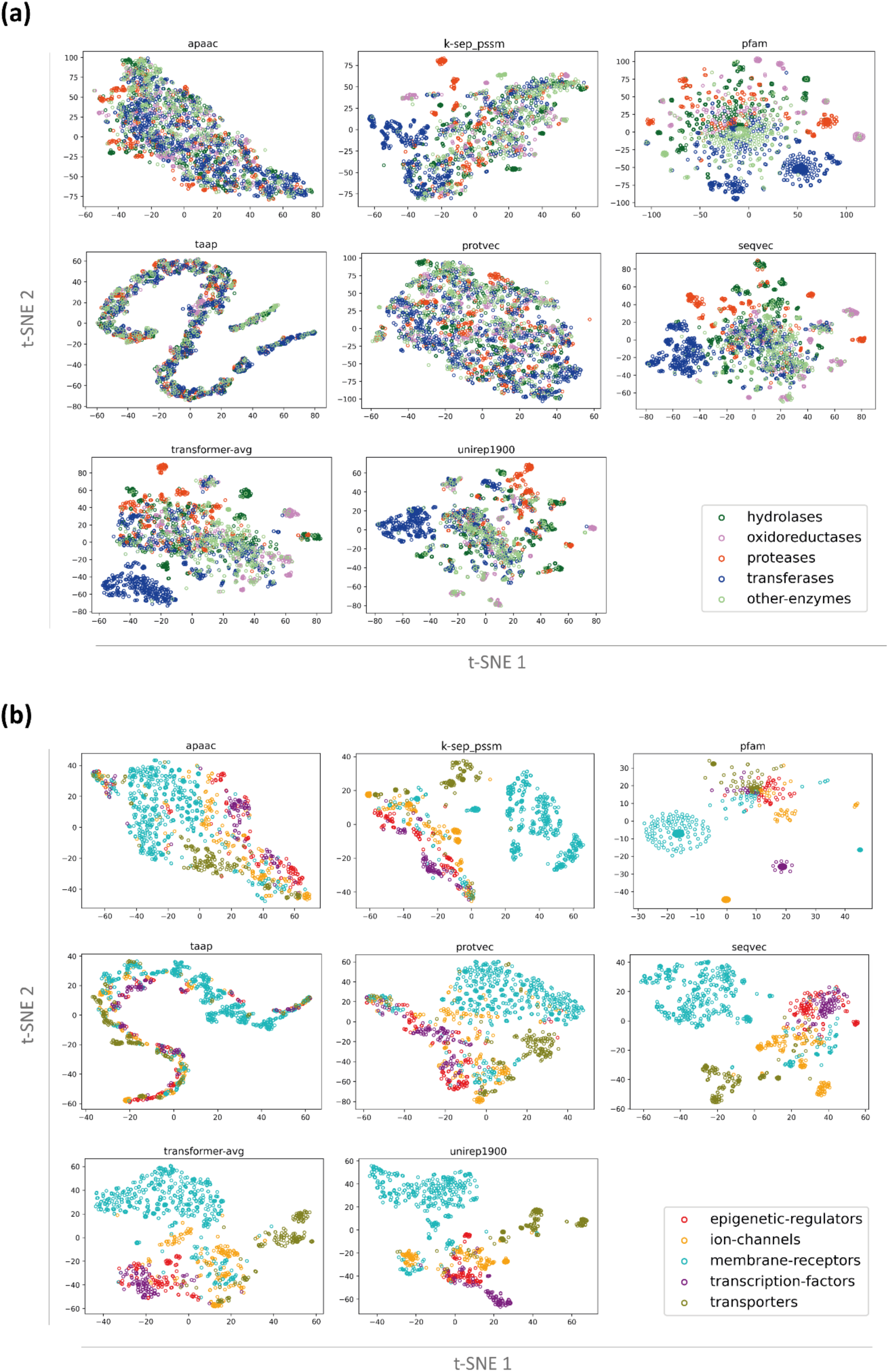
t-SNE based visualization of conventional (apaac, k-sep_pssm, pfam, taap) and learned (protvec, seqvec, transformer-avg, unirep1900) protein representations on; **(a)** enzymes including hydrolases, oxidoreductases, proteases, transferases, and other-enzymes groups, and **(b)** non-enzyme protein families including epigenetic regulators, ion channels, membrane receptors, transcription factors, and transporters, in different colors.

#### 3.1.2. Split-based characteristics of protein family-specific datasets

##### Pairwise similarity distributions

To explore protein and compound diversity in our datasets, we evaluated protein-protein and compound-compound pairwise similarities of the members of a selected representative protein family (i.e., transferases), in terms of “train vs. train”, “test vs. test”, and “train vs. test” dataset comparisons for each split strategy (i.e., random-split, dissimilar-compound-split, and fully-dissimilar-split). For this, we aligned protein sequences using EMBOSS Needle global pairwise sequence alignment tool ^73^ and plotted histograms based on identity values of unique protein pairs in the corresponding datasets. We extracted pairwise compound similarities by calculating Tanimoto coefficient between fingerprint representations using the *simsearch* function of the Chemfp python package ^74^. Since it was highly infeasible to calculate pairwise similarities for billions of compound pairs, we randomly sampled 10% of all compounds in the transferases dataset and set the minimum similarity detection threshold as 0.1. Again, we only considered a unique list of compound pairs.

Figure 3 displays similarity distributions of pairs of proteins and compounds involved in the transferases dataset, in which the values may be greater than one since the plot is normalized to equalize the total area to one (i.e., the density plot). Having a similarity value in the range of 0 - 0.5 for the majority of protein and compound pairs in all plots demonstrates the high diversity of samples which is a desirable characteristic for computational bioactivity modelling. As displayed in Figure 3, similarity distributions only slightly change between different split methods, considering “train vs. train” and “test vs. test” sample similarities, whereas there are significant differences between the samples of “train vs. test”, for both compounds and proteins, in terms of different splits. The absence of similarity values greater than 0.5 for compound “train vs. test” pairs in the dissimilar-compound-split dataset, and both protein and compound “train vs. test” pairs in the fully-dissimilar-split dataset validates the similarity-centric characteristics of our datasets. Exceptional pairs of proteins with high similarity values in the fully-dissimilar-split dataset stem from the discrepancies between UniRef50 clusters and our pairwise alignment results, and their number is found to be insignificant (please note that the frequencies are given on logarithmic scale in Figure 3). These results validate the capability of our methodology in terms of producing challenging (and presumably realistic) train-test datasets, so that the bioactivity prediction models trained and tested on these datasets hopefully reflect the real-world performances while discovering novel drug candidates and/or new targets.

**Figure 3.**
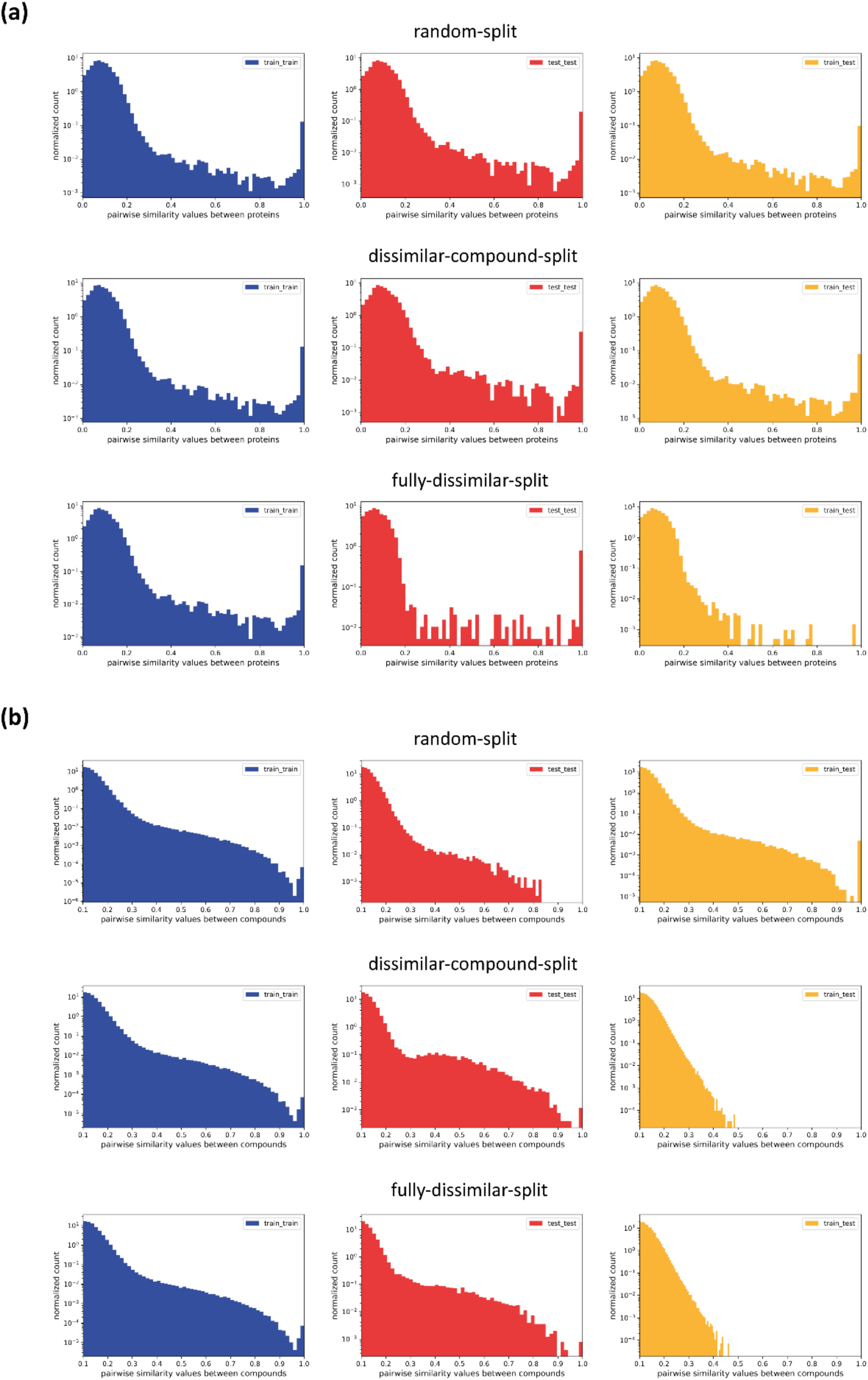
Pairwise similarity distributions of **(a)** proteins and **(b)** compounds for “train vs. train”, “test vs. test”, and “train vs. test” samples in random-split, dissimilar-compound-split, and fully-dissimilar-split of the transferases dataset (shown in the logarithmic scale).

##### Bioactivity distributions

We also plotted bioactivity distributions of protein family-specific datasets based on train-test samples of each split. Figure 4 displays pChEMBL value-based histograms for transferases, ion channels, and membrane receptors (plots for the remaining families are available in ESI Figure S2). Median bioactivities vary between 5.7 and 7.1 for different protein families. When comparing bioactivities of train and test sets of each family, it is observed that distributions have similar shapes, regardless of the dataset split strategy. In addition, they generally have very similar mean and median values, although the difference is slightly higher in the fully-dissimilar-split datasets of some families. Having bioactivity distributions that are consistent with each other in training and test folds implies good coverage of bioactivity data and supports the suitability of our large-scale datasets for bioactivity modelling. These results also indicate that a stratified-split strategy is not required for our datasets.

**Figure 4.**
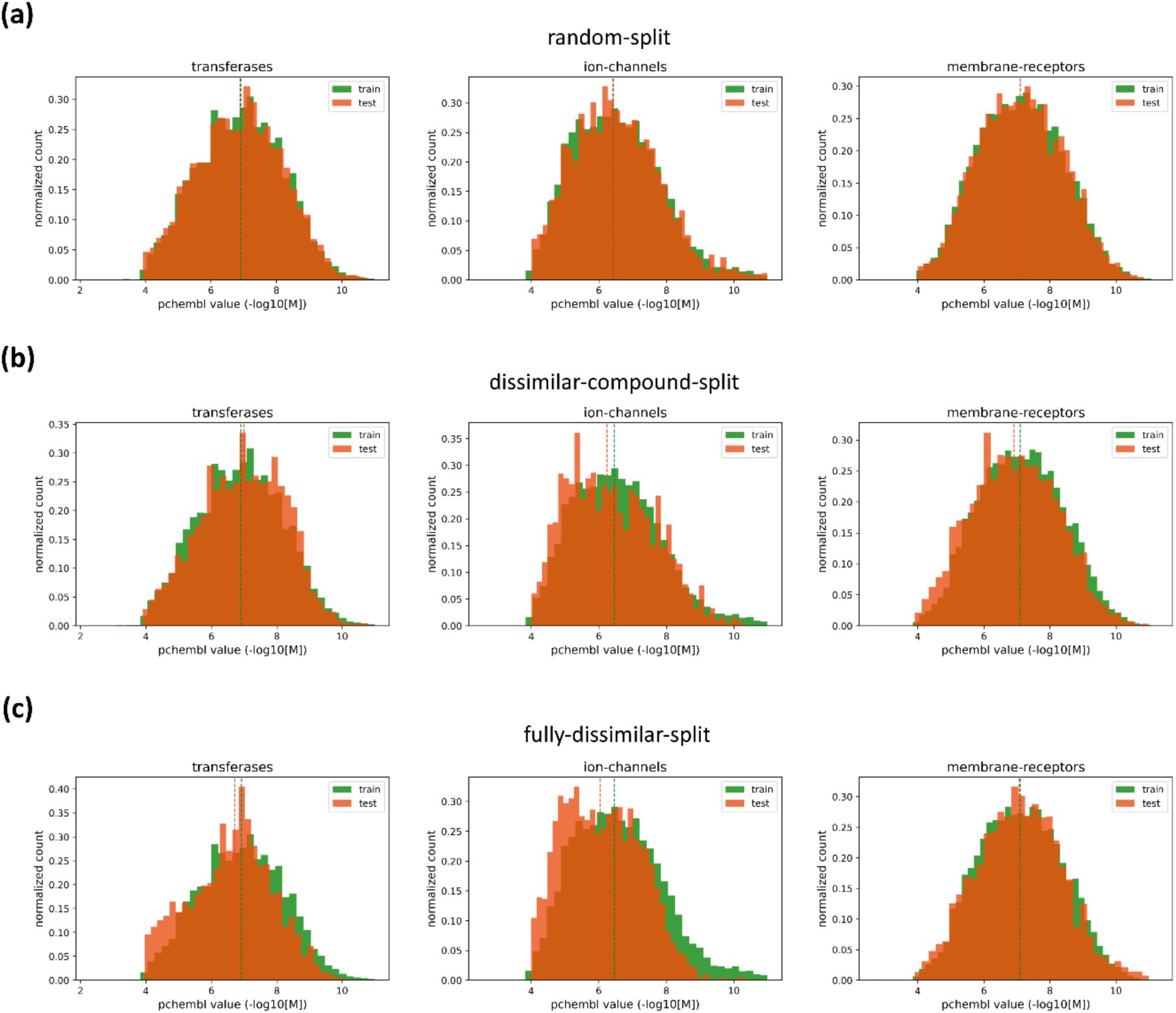
Histogram plots displaying bioactivity distributions of transferase, ion channel, and membrane receptor families based on train (green bars) and test (orange bars) samples of; **(a)** random-split, **(b)** dissimilar-compound-split, and **(c)** fully-dissimilar-split datasets, along with their median values shown as vertical dashed lines.

##### t-SNE projection of train-test datasets for three splits

In this analysis, we visualized the distribution of bioactivity data points (i.e., compound-protein pairs) on 2-D via t-SNE projections to observe how train and test fold samples are separated from each other under different splitting settings. For each protein family-based dataset, 1,500 data points were randomly selected (from both train and test folds), since the number of training samples dominates test samples in the original datasets. Each bioactivity data point was represented by the concatenation of its protein and compound feature vectors, and used as input to the t-SNE algorithm.

In Figure 5, t-SNE plots of transferases and ion channels (i.e., the representative families, as these are two widely utilized target families in drug discovery) are given for k-sep_pssm and unirep1900 representations. Panel a, b, and c correspond to the random-split, dissimilar-compound-split, and the fully-dissimilar-split datasets, respectively. For the random-split dataset, 2-D embeddings of the train and test samples largely overlap, since they share similar proteins and compounds. These overlaps significantly decrease in dissimilar-compound-split dataset and almost disappear in the fully-dissimilar-split dataset, as expected. This analysis can be considered as a visual validation of the implemented splitting strategies, and it provides clues about the difficulty levels of our prediction tasks.

**Figure 5.**
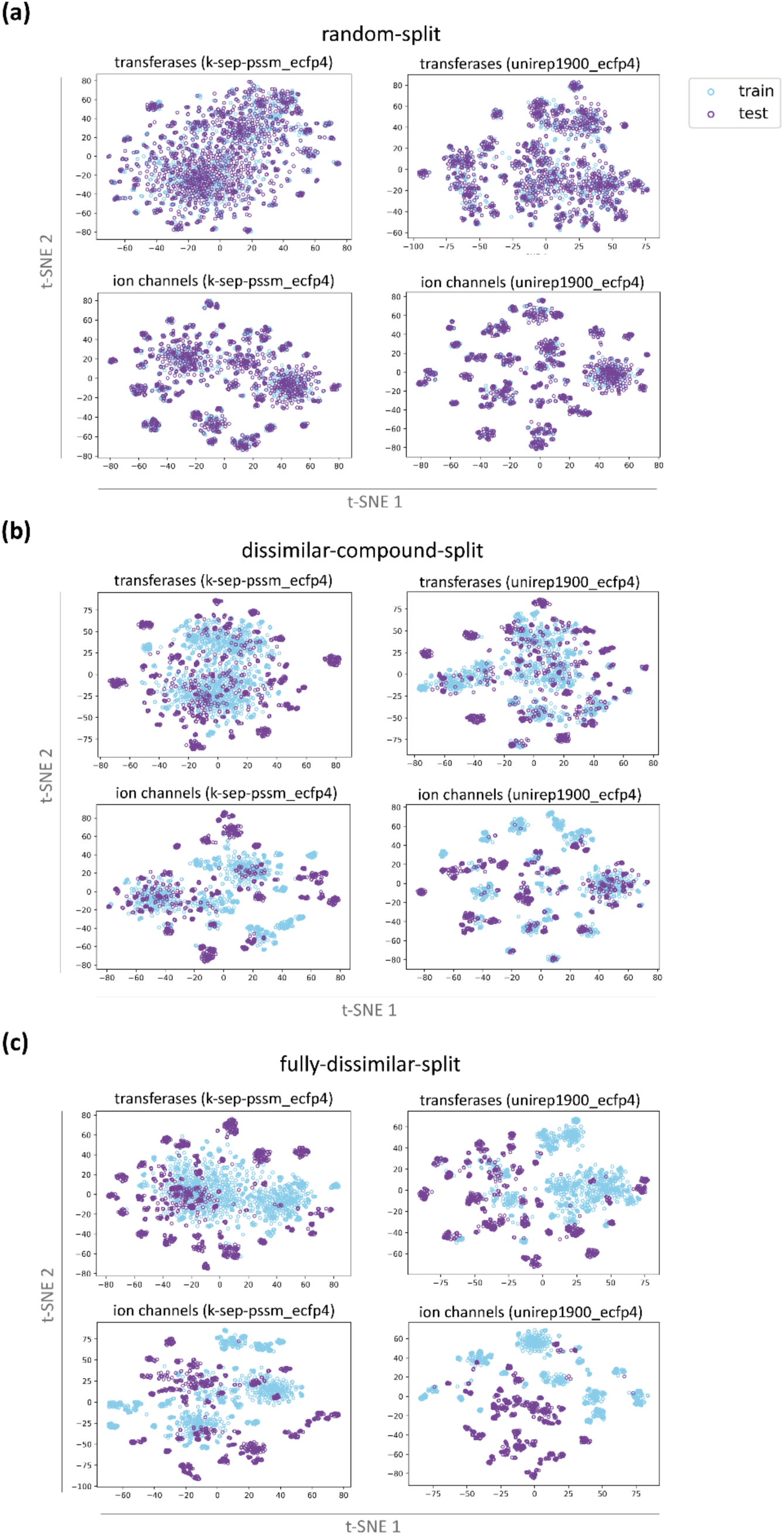
t-SNE projections of train-test samples (i.e., compound-protein pairs) of transferase and ion channel families for k-sep_pssm and unirep1900 representations on; **(a)** the random-split, **(b)** dissimilar-compound-split, and **(c)** the fully-dissimilar-split datasets.

### 3.2. Small-Scale Analysis (Target Feature-based Modelling)

There are numerous conventional descriptor sets for proteins in the literature, most of which can be utilized for DTI prediction. Evaluating all descriptor sets on our large-scale datasets would not be feasible considering the computational cost; as a result, we decided to carry out a small-scale analysis to pre-select the descriptor sets that are successful in DTI prediction, and use the selected descriptors in both the medium-scale and large-scale analysis later. Additionally, it was required to determine the supervised learning algorithm to be used for DTI prediction in this study, and due to, again, the computational complexity related issues, we decided to make a performance comparison (between SVM and RF) on these small-scale datasets.

In this analysis, we assessed the success of SVM- and RF-based DTI prediction models, each utilizing one of the 42 conventional protein descriptor sets (including the ones explained in section 2.2.1. and additional ones that fall into the same categories as these), and a baseline (i.e., the random200 descriptor). The full list of descriptors sets is provided in ESI Table S5, and details can be obtained from ^54^ and ^64^. The models are trained and tested on 9 independent compound-centric datasets (i.e., the clusters of Curcumin, Tamoxifen, Quercetin, Genistein, Econazole, Levoketoconazole, Amiodarone, Miconazole, and Clotrimazole) via nested cross-validation using the target feature-based modelling approach (please see section 2.3.1). In this approach, the system only employs protein features as input, so it eliminates the effect of compound representations on the model prediction performance, which is expected to provide a suitable setting for an initial comparison of protein representations. Here, the task of each model is the binary classification of input proteins, as active or inactive, against the corresponding compound cluster.

Figure 6 displays mean F1-score and MCC values of 9 datasets for each representation model, in which orange and blue colors correspond to SVM and RF models, respectively (all results including accuracy, precision, recall, F1-score, and MCC metrics are given in ESI Table S5). The ranking of protein descriptor sets on the horizontal axis was done according to decreasing RF model scores. Figure 6 clearly displays that RF models outperform SVM models with a few exceptions such as the pfam model in terms of the MCC score. When model performances are compared in terms of protein representations, pssm-based descriptors perform better than other descriptors in general. These results indicate that evolutionary relationships of proteins carry important knowledge regarding bioactivity/interaction mechanisms. Some of the sequence composition-based descriptors such as dde, tpc, and spmap, and physicochemistry-based descriptors such as apaac and paac, also performed well. Moreover, obtaining scores that are significantly higher than the baseline (i.e., random200), even for the models with the lowest performance, implies that protein representations carry signals/patterns relevant to bioactivity modelling. However, these results cannot be generalized as they cover only a small portion of the bioactivity space; thus, it is important to observe how these models behave when the data scale is changed.

**Figure 6.**
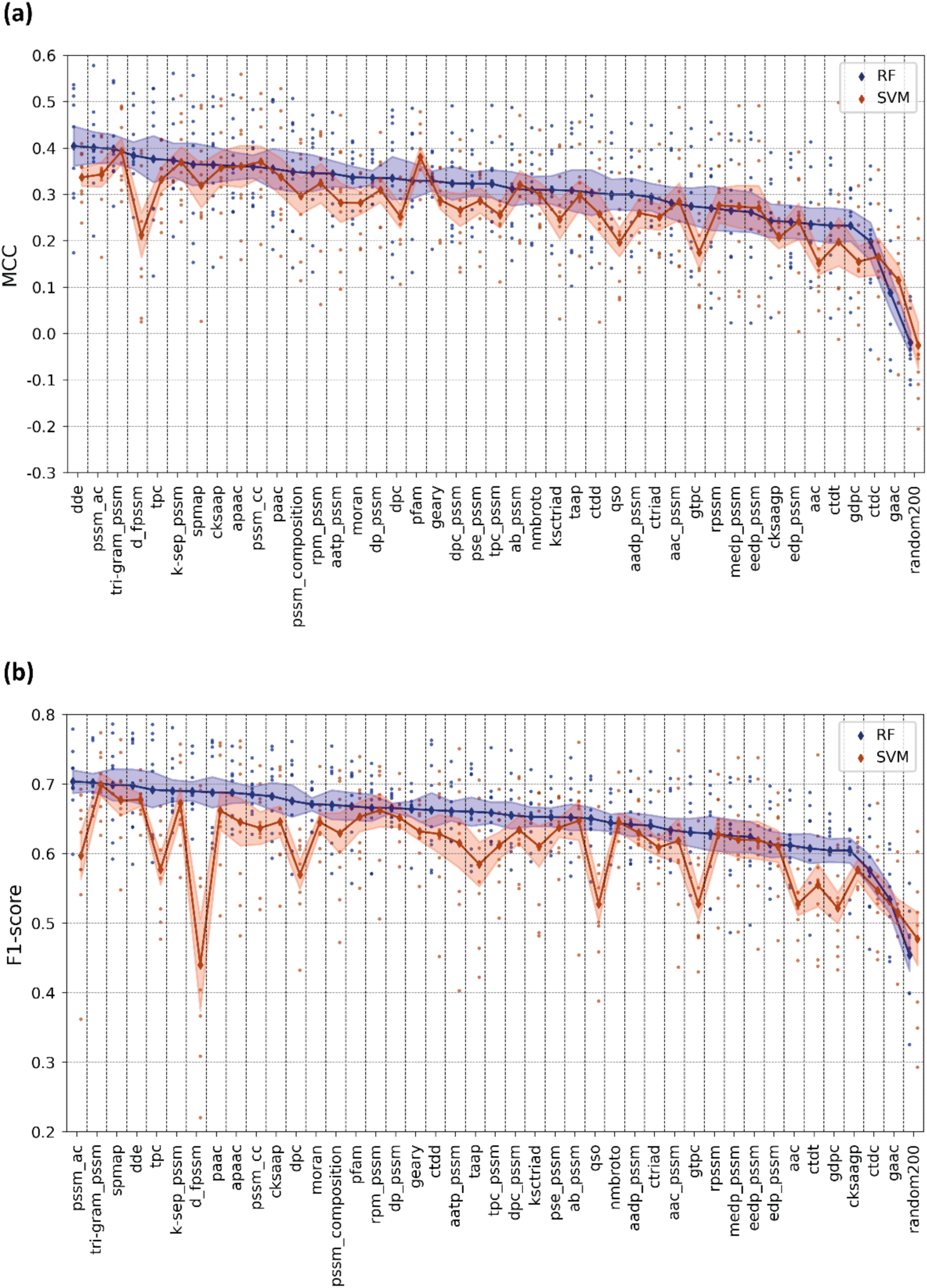
Mean **(a)** MCC and **(b)** F1-score test results of RF- and SVM-based DTI prediction models constructed via target feature-based modelling approach.

At the end of this analysis, we decided to continue with RF, to be used throughout the study. Also, we selected 10 conventional descriptor sets with both high and low performances, and distinct properties regarding the protein features they incorporated and used them in the following benchmarks (i.e., apaac, ctdd, ctriad, dde, geary, k-sep_pssm, pfam, qso, spmap and taap).

### 3.3. Medium-Scale Analysis (PCM Modelling)

PCM modelling approach can handle high numbers of training instances, belonging to different compounds and proteins, within a single predictive model, in contrast to ligand- and target feature-based modelling which requires the generation of separate models for each protein or compound (or compound cluster), respectively. Thus, PCM modeling brings the advantage of learning from larger datasets, which is a critical requirement in machine learning, in general. Another advantage of PCM modeling is the joint utilization of compound and protein features to better model their interaction-related properties, without the requirement of scarce and difficult to analyze 3-D structural information, unlike target-based structure modelling approaches. In the following benchmarks, we aimed to evaluate protein representations in terms of PCM modeling, over the problem of regression-based DTI prediction.

As mentioned above, we selected 10 out of the total 42 descriptor sets by removing the ones that have similar featurization approaches and similar performance scores in the previous small-scale analysis. As a result, we constructed PCM models for 10 selected conventional protein descriptor sets and 6 learned protein embeddings using RF regression algorithm on the mDavis kinase dataset. We also built 2 baseline models using randomly generated representations, within the same setting (please see section 2.3.2).

Model performance results based on RMSE, Spearman rank correlation, MCC and F1-score (all computed on the hold-out test set of the mDavis dataset) are given in Figure 7 (also available in ESI Table S6). The results indicate that the rankings of models are mostly consistent among both classification and regression metrics with slight differences, excluding pfam. As a domain profile-based descriptor set, pfam is the best performing model in terms of F1-score (0.538) and has a moderately high MCC score (0.41); however, it is also one of the worst performers in terms of RMSE (0.854) and Spearman (0.497) scores. It can be inferred from these results that domain profiles of proteins might not contain sufficient information to make precise bioactivity value predictions, but it can be useful if the aim is just to classify protein-compound pairs as active or inactive (i.e., binary prediction). The results also indicate that the seqvec model displays the best performance for almost all metrics (RMSE: 0.794, Spearman: 0.571, MCC: 0.445, F1-score: 0.53). Apart from seqvec, other learned embeddings also have higher performance scores compared to conventional descriptors in general. Mean Spearman rank correlation and MCC scores of learned representations are 0.530 and 0.417, respectively, whereas the same scores are 0.511 and 0.388 for conventional descriptor sets. Learned embeddings do not utilize any molecular or biological knowledge during their self-supervised training, but still, they are capable of representing proteins that yield high performance DTI prediction. Well performing descriptors in the previous small-scale analysis, k-sep_pssm (homology) and apaac (physicochemistry), also have competitive performance results here (Spearman: 0.545 and 0.532, respectively). On the other hand, dde (Spearman: 0.508) and spmap (Spearman: 0.491) could not yield their high ranks here in the medium-scale analysis (i.e., dde and spmap had the ranks of 1 and 8 on the small-scale, whereas, they ranked 9 and 16 on the medium-scale, respectively). It is possible to state that while homology- and physicochemistry-based descriptors gained from increased dataset size (i.e., for apaac and k-sep_pssm, small-scale analysis mean MCCs are 0.361 and 0.374, respectively, whereas their medium-scale analysis mean MCCs are 0.418 and 0.434), sequence composition could not improve its performance when trained on larger datasets.

**Figure 7.**
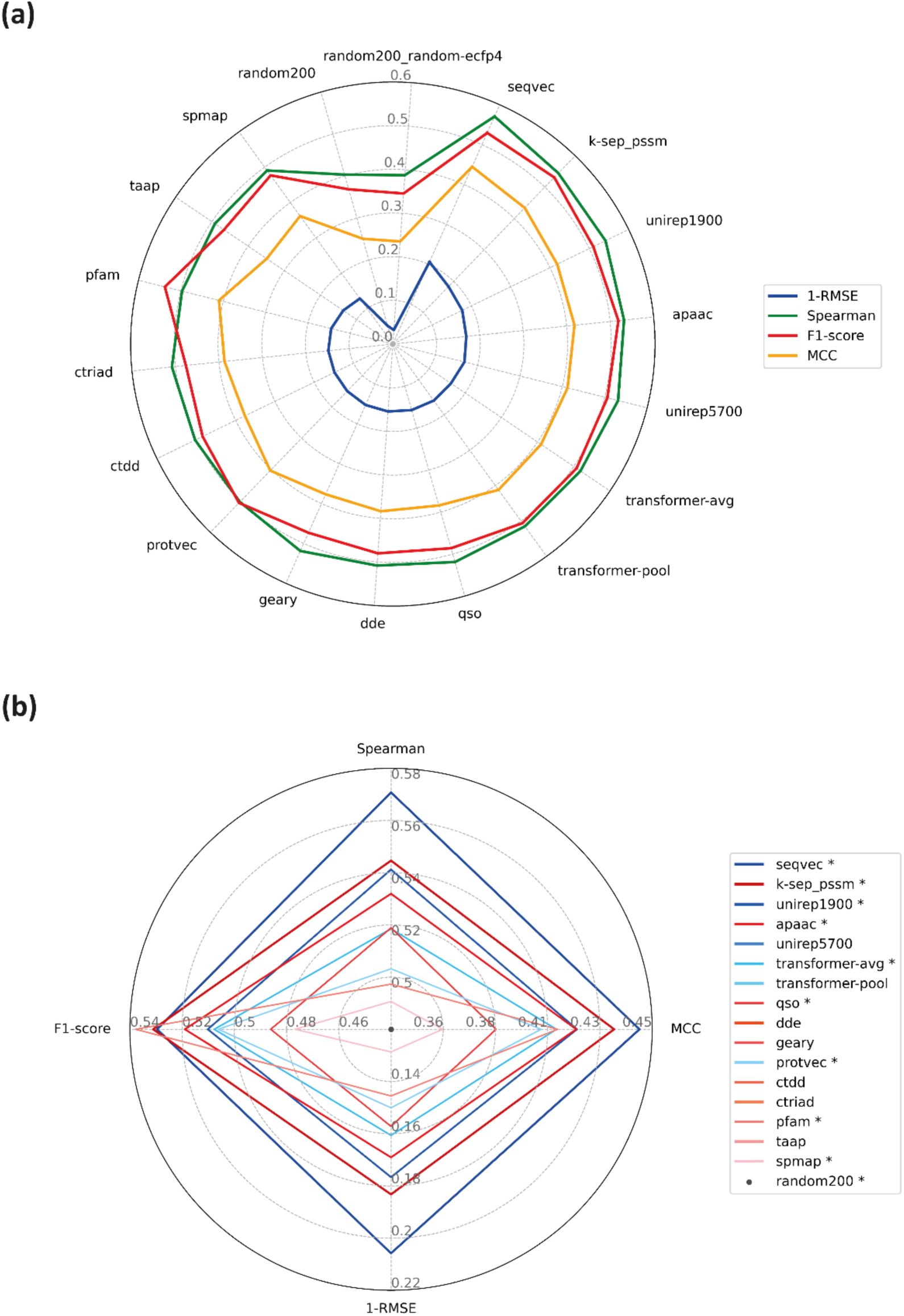
Test performance results of medium-scale PCM models (on the mDavis dataset) based on RMSE (the scores are reported as 1-RMSE, so higher values represent better performance), Spearman’ s rank correlation, MCC and F1-score; **(a)** each color corresponds to an evaluation metric, and **(b)** scores are displayed only for the selected representative models (marked with asterisk in the legend). The ranking in the legend is based on the models’ performance from best to worst according to their RMSE scores. Shades of red and blue represent conventional descriptors and learned representations, respectively.

Also, there is an overall increase in MCC scores of conventional descriptor sets (excluding dde and spmap) when we compare the results of small- and medium-scale analyses. In addition to the contribution of the increased sample size, this situation can be associated with the involvement of compound features in PCM-based models, which probably led to a better learning over the joint protein-compound interaction space. On the other hand, PCM models here had lower F1 scores than the target feature-based models in the small-scale analysis. In order to calculate MCC and F1-scores for PCM models, we converted real-valued predictions into binary format at the cut-off value pChEMBL = 7, which is also used in other studies as a bioactivity threshold for kinase inhibitors ^75^. However, only 27% of the test samples became active at this threshold, causing a class imbalance in the mDavis kinase dataset. Although F1-score is one of the most widely used metrics for classification tasks, it is not as robust against the class imbalance problem as the MCC metric, which makes MCC a more appropriate option in this scenario, Finally, the baseline models displayed the lowest performances, similar to the results of the target feature-based modelling analysis.

### 3.4. Large-Scale Analysis (PCM Modelling)

The main goal of this analysis is evaluating protein representations over a highly realistic scenario, especially in terms of discovering new drugs and/or targets, using our carefully prepared large-scale datasets, and to compare their overall performance in machine learning-based DTI prediction. Secondly, we aimed to display how model performances can change dramatically when the same samples are distributed to train and test sets differently, to point out the importance of train-test data split. Furthermore, we evaluated the suitability of various performance metrics under different modeling approaches.

For the large-scale benchmark analysis of protein representations, we constructed protein family-specific bioactivity datasets including enzyme (i.e., transferases, hydrolases, oxidoreductases, proteases, and other enzymes) and non-enzyme groups (i.e., membrane receptors, ion channels, transporters, transcription factors, and epigenetic regulators). For each family, three versions of train-test splits with differing difficulty levels were constructed by considering pairwise similarities of proteins and/or compounds (please see section 2.1.3 for details). PCM models were trained independently on each of these splits using the same protein representations employed in the previous (medium-scale) analysis. As a result, 600 DTI prediction models were built, trained, and tested in total (please see section 2.3.2 for details).

We evaluated model performances from several perspectives using multiple scoring metrics. Median corrected RMSE and Spearman correlation scores are displayed as line plots in Figure 8, in which the light colored (transparent) circles indicate individual model performances on each protein family, and the dark colored diamonds represent mean scores averaged over all families. The models are ranked according to descending performance on the fully-dissimilar-split dataset (for both metrics). In Figure 9, model performances are provided as box plots over three different forms of the MCC metric. The models are ranked according to descending mean values of median corrected MCC scores for the fully-dissimilar-split and dissimilar-compound-split datasets, and according to multiclass MCC scores for the random-split dataset. Protein family-specific performances are available at ESI Table S7.

**Figure 8.**
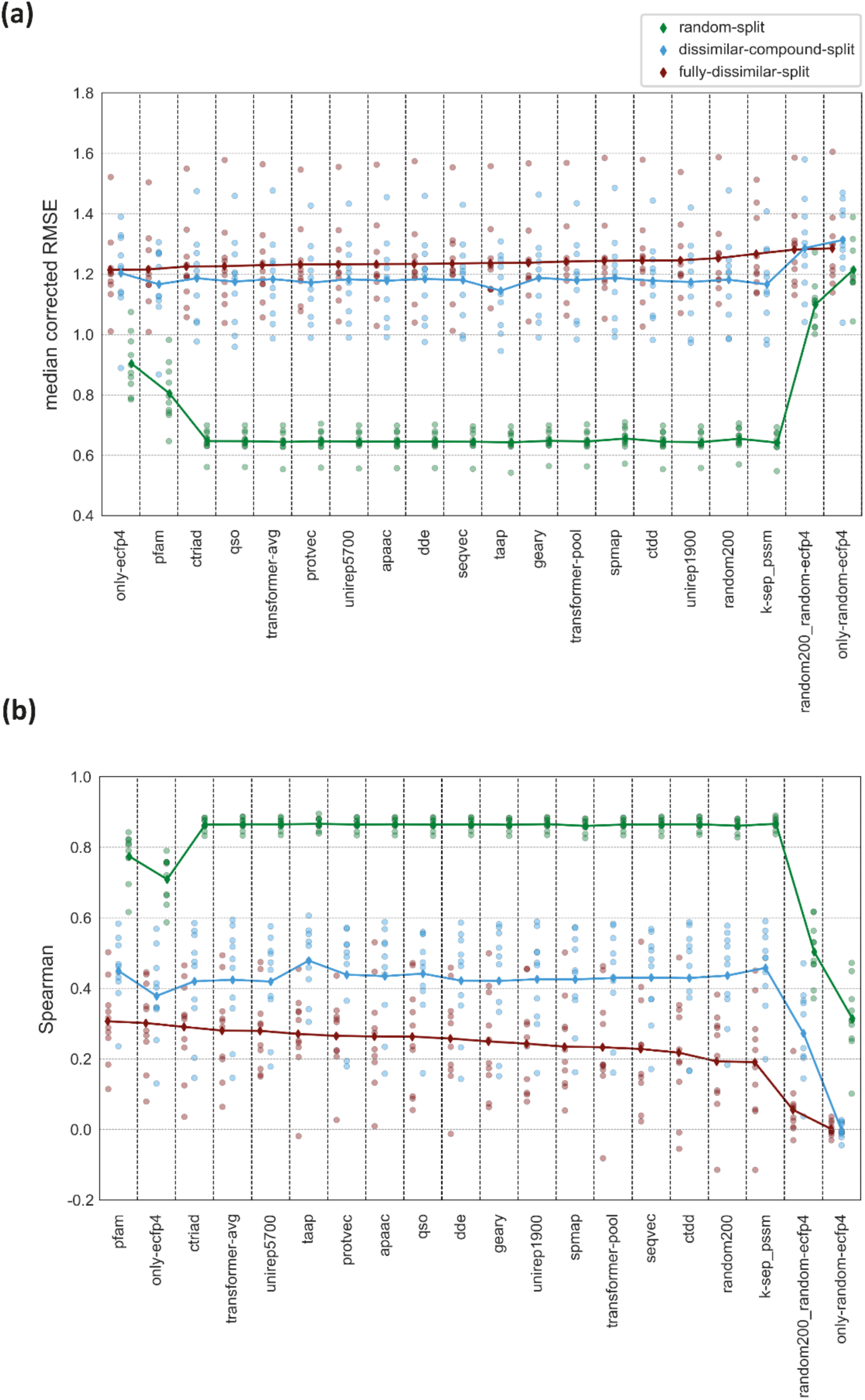
Regression-based test performance results of protein family-specific PCM models (each using a different representation type as input feature vectors) for random-split, dissimilar-compound-split, and fully-dissimilar-split datasets based on **(a)** median corrected RMSE, and **(b)** Spearman correlation scores. The models are ranked according to decreasing performance on the fully-dissimilar-split dataset.

**Figure 9.**
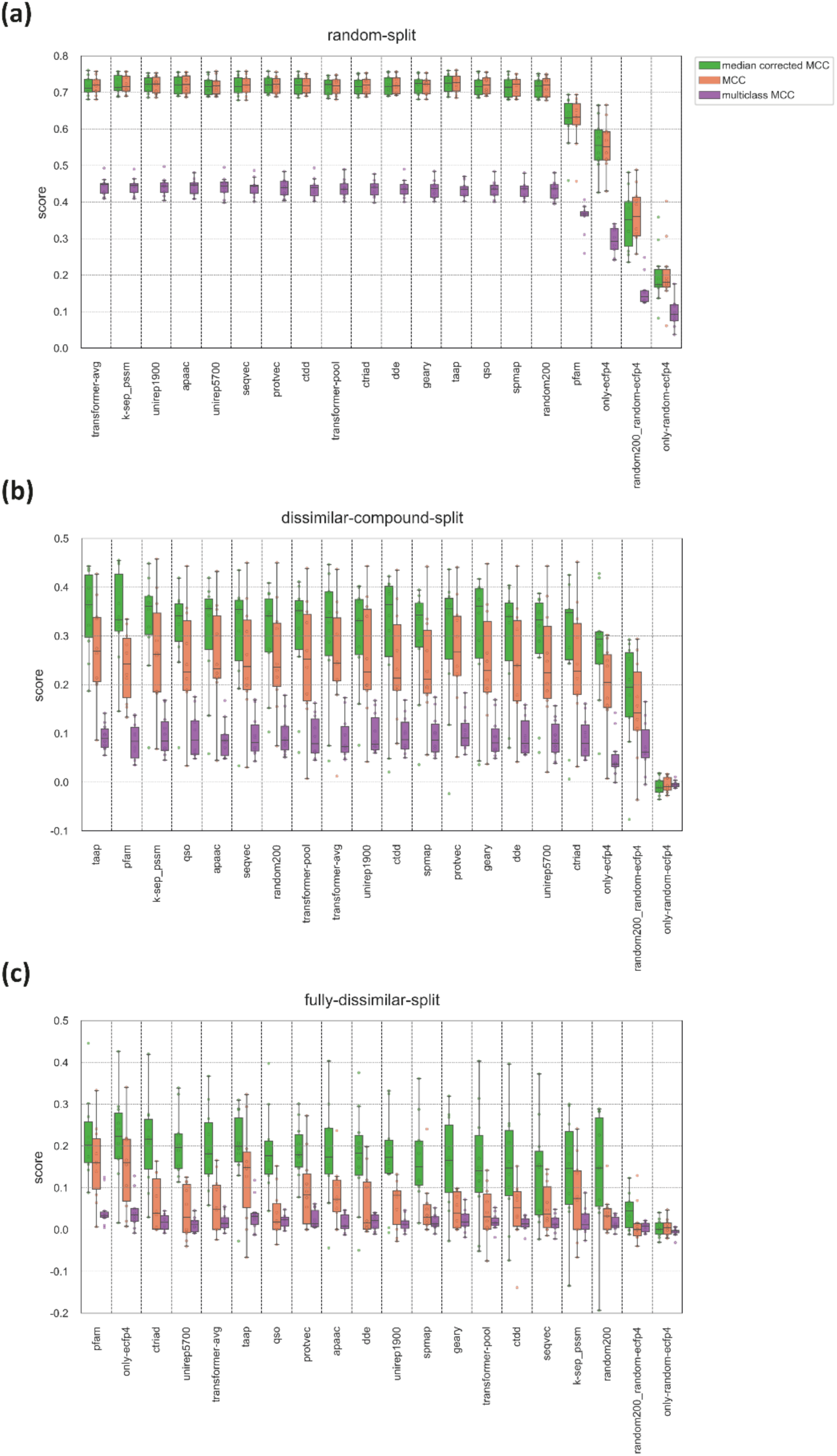
Classification-based test performance results of protein family-specific PCM models (each using a different representation type as input feature vectors) in terms of MCC scores for **(a)** random-split, **(b)** dissimilar-compound-split, and **(c)** fully-dissimilar-split datasets. The models are ranked according to decreasing performance on the fully-dissimilar-split dataset.

#### Investigation on performance metrics

The intra-family rankings of models are generally consistent with each other among five different metrics (Table S7). However, there are some discrepancies between the scores depending on the data split. Considering regression metrics, some of the models trained/tested on the fully-dissimilar-split and dissimilar-compound-split datasets show high performance in terms of RMSE (i.e., low RMSE values), whereas at the same time, they displayed low Spearman correlations, which indicates inconsistency. In challenging scenarios (e.g., on the fully-dissimilar-split and dissimilar-compound-split datasets), continuous value-based prediction of bioactivities (via regression) is unstable and unreliable due to the difficulty of the task. Thus, it would be a better choice to evaluate model performances on Spearman rank-based correlation scores on these challenging datasets, while RMSE could be a better option to differentiate models trained and tested over easy cases (i.e., the random-split dataset). In classification-based assessment, the single class MCC metric is not as restrictive as the regression or multiclass evaluation metrics since it is less sensitive to deviations in prediction values. However, it may suffer from the shifted mean problem when applied to regression-based PCM models by binarizing bioactivity values (see Methods 2.5). Obtaining MCC values close to 0 (Figure 9) despite moderate Spearman correlation scores (Figure 8) on challenging datasets is a sign of a systematic shift on model prediction outputs, which we handled by conducting median correction on the real-valued prediction results, as explained in Methods 2.5. In Figure 9, it can be observed that median correction provided a significant increase in single class MCC scores of the fully-dissimilar-split and dissimilar-compound-split datasets. Also, median corrected MCC scores are highly consistent with the Spearman correlation scores (Table S7). Considering the multiclass MCC metric, prediction scores are around zero for most of the models on challenging split sets. Since this metric expects prediction values to fit narrow intervals, it is more restrictive than the single class-based metrics. However, this seems to be an advantage for evaluating models on the random-split set. As seen in Figure 9a, on the random-split dataset, the variance of the mean multiclass MCC score distribution is greater than the single class MCC scores (i.e., models are better separated from each other). Furthermore, its ranking is highly consistent with the results of the medium-scale experiments, in which the top performers were learned representations, together with k-sep_pssm and apaac conventional descriptor sets.

Thus, it can be inferred that the multiclass MCC metric discerns models better than binary class MCC in the random data split setting, and it partly handles the overfitting problem which frequently occurs on randomly split large-scale datasets.

#### Evaluation of protein representations

Performance results in Figure 8 and 9 clearly indicate that the representation capability of different protein descriptor sets depend on the protein family and the difficulty level of the split used for training and testing. Also, there is no significant difference between the mean performances of different protein representations for a particular dataset split, with a few exceptions. Considering family-based performance averages, pfam is one of the best representations on the fully-dissimilar-split and dissimilar-compound-split datasets, while it is the lowest performer on the random-split dataset (Figure 8 and 9). Contrary to pfam, k-sep_pssm is one of the best performers on the random-split and dissimilar-compound-split datasets but the worst one on the fully-dissimilar-split dataset (Figure 8 and 9), though the performance results on the random-split dataset are very close to each other. As a homology-based descriptor set, k-sep_pssm is expected to capture hidden similarities between evolutionarily related sequences, especially by taking advantage of the presence of highly similar proteins between the train and test splits. On the other hand, the utilization of protein domain profiles seems to make pfam more suitable for acquiring bioactivity related information from evolutionarily distant sequences, probably due to highly sensitive HMM-based domain/family profile search procedures implemented in Pfam and similar databases. Interestingly, taap, which is a simple descriptor set, is involved in the top-performing PCM models for all dataset splits. However, taap was one of the lowest performers in the small-(among the selected 10 conventional descriptor sets) and medium-scale analyses. Its simplicity is observed to become an advantage with the increase in bioactivity dataset size and complexity. Apart from these, physicochemistry-based descriptors including apaac (in all splits), ctriad (on the fully-dissimilar-split dataset) and qso (on both the fully-dissimilar-split and dissimilar-compound-split datasets), and learned representations perform well in the large-scale analyses. In particular, the top performance results of unirep5700 and transformer-avg on the fully-dissimilar-split dataset demonstrate the potential of protein representation learning methods in the data-driven DTI prediction.

We also conducted protein family-specific evaluations to understand whether different protein representations display similar results across families. In Figure 10, we plotted the performance of the models of protease and the ion channel families, in the form of a conventional descriptor set vs. learned representation comparison, using the Spearman and median corrected MCC scores, for all three dataset splits. For a fair comparison, we selected four well-performing conventional descriptors instead of including all of them, since we have only four different types of learned representations. For this, we involved apaac, k-sep_pssm, pfam, and taap as conventional descriptor sets and protvec, seqvec, transformer-avg, and unirep5700 as learned representations. Figure 10 shows that learned representations outperform conventional descriptors in the challenging splits of proteases, considering both metrics. However, the results are the opposite for the ion channel family, on which the conventional descriptor sets performed better. On the random-split dataset, there is no observable difference between conventional descriptor sets and the learned representations, probably due to the non-discriminative characteristic of this data splitting strategy, which poses non-challenging cases for all models.

**Figure 10.**
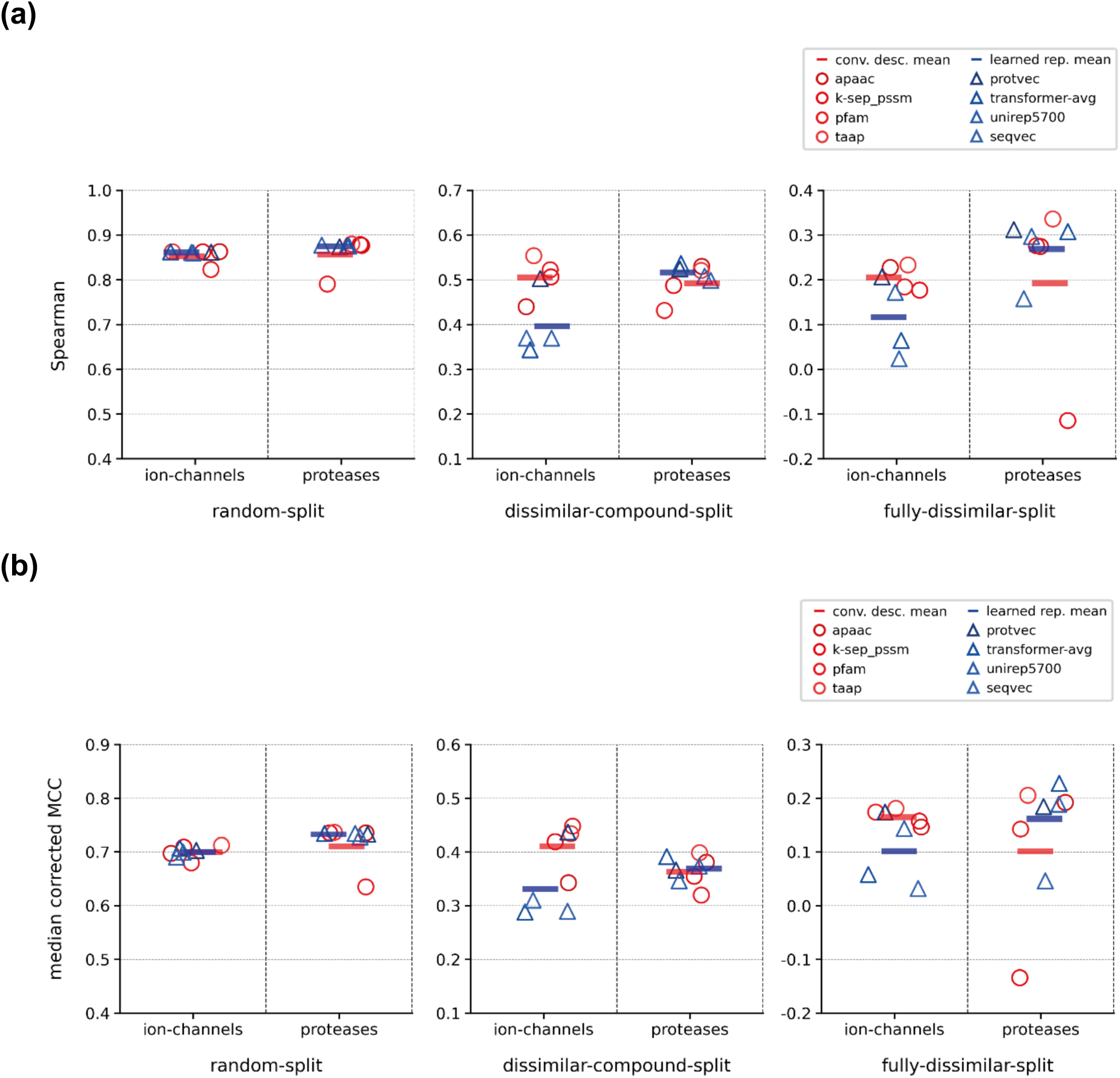
Performance comparison of well performing conventional descriptor sets and learned representations for three different splits of ion-channel and protease family datasets in terms of; **(a)** Spearman rank correlation, and **(b)** median corrected MCC scores.

Results presented in Figure 10 are also correlated with the scores on other protein families (Table S7), as a result, it is possible to state that conventional descriptor sets perform slightly better than learned representations on non-enzyme protein families, whereas learned representations perform slightly better on enzyme families. This may be due to the intrinsic properties of these protein families, such as, the information about the arrangement of amino acids on the sequence may carry critical signals regarding ligand interactions of enzymes, whereas, physical and chemical properties of amino acids may be more important to model the interactions of non-enzyme families like membrane receptors, ion channels, and etc. In fact, protein representation learning models comprise a critical advantage considering this matter. It is possible to limit the training dataset of these models to a specific protein family, so that, the resulting representations will contain information specific to that family. All of the learned representations we used in our study were trained on large datasets including all protein families, and this may have prevented learning family specific properties.

When taking all these findings into account, we can clearly state that the representation capabilities of different featurization approaches considerably vary among protein families and splitting strategies, even though some common inferences can be made. We believe that, while choosing a featurization approach in DTI prediction, protein family-specific findings should be taken into account, rather than considering the overall (i.e., average) results. Regarding learned representations, re-training (or fine tuning) the models using a distinct dataset with desired characteristics (e.g., members of a certain family) may be a good choice to better learn the features specifically associated with that group of proteins.

#### Comparison of data splitting strategies

To compare models across three dataset splits, we plotted performance scores by pooling 200 models of each split (including the baseline models) without any grouping by families or representation methods. The results are displayed in Figure 11 via violin plots. This figure shows that there is a significant decrease in overall performances with the increasing difficulty levels of splits, which is not a surprising outcome. Nevertheless, it highlights the importance of splitting datasets into train/test folds for performance evaluation, with the aim of preventing the reporting of over-optimistic results and yielding the fair assessment of model successes. Figure 11 also displays that the model performances are distributed more evenly over the whole range of scores in the fully-dissimilar-split and dissimilar-compound-split datasets, compared to the random-split dataset, in which most of the models produced very similar scores, creating a dense region on the plot. This observation indicates randomsplitting has less power in terms of distinguishing different models from each other.

**Figure 11.**
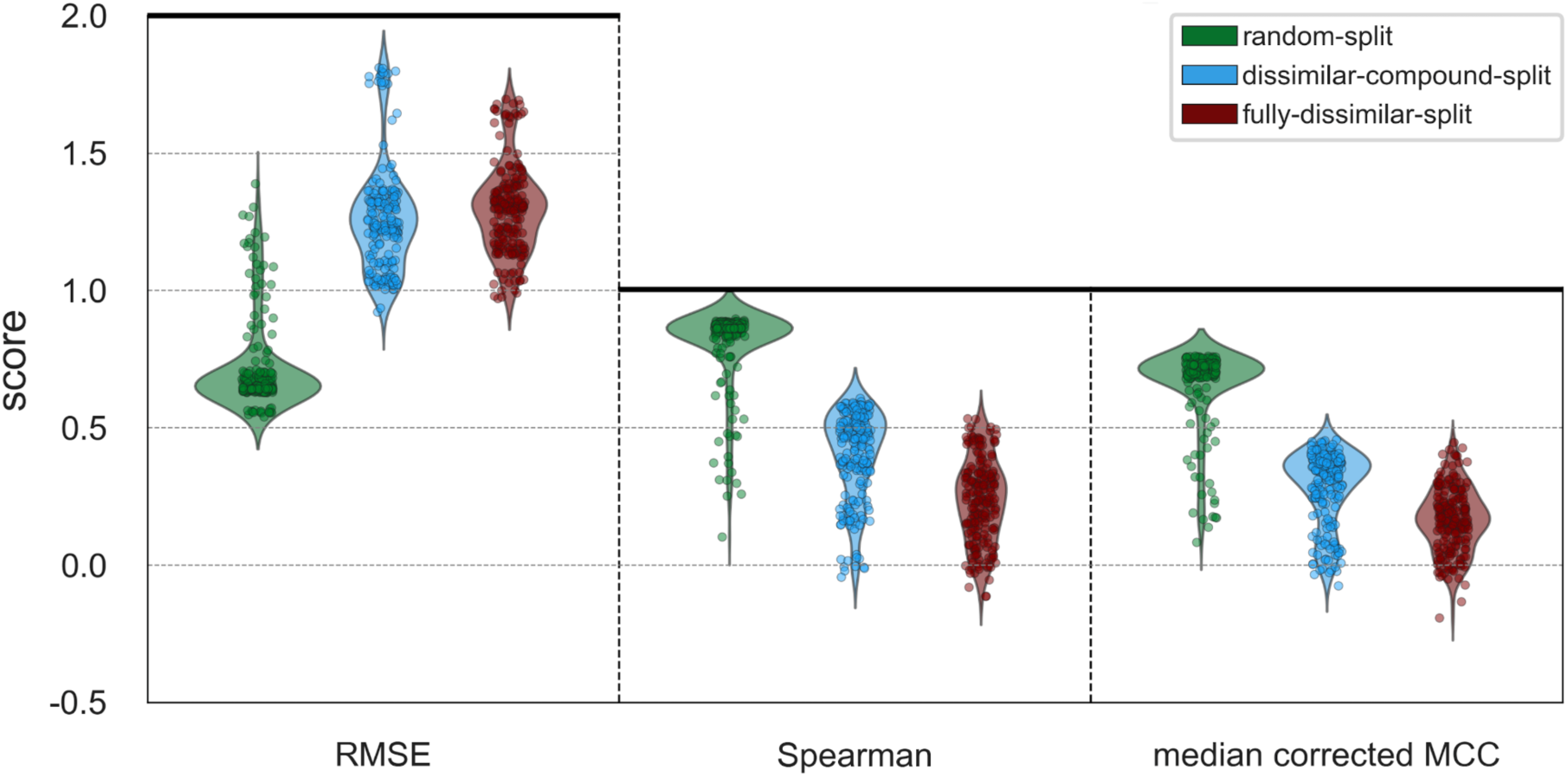
Split-based test performance scores of family-specific PCM models in terms of RMSE, Spearman rank correlation, and median corrected MCC metrics.

In the fully-dissimilar split, neither similar proteins nor similar compounds are shared between train and test folds. As a result, this dataset is suitable to evaluate the performance of DTI prediction models in terms of predicting novel ligands to understudied targets (or completely new target candidates). Whereas in the dissimilar-compound split, similar proteins are presented in between train and test sets. Nevertheless, it is useful for discovering novel ligands against well-studied target proteins, or proteins for which structurally highly similar and well-studied targets exist.

#### Examination of baseline models

Table 2 contains family-based average Spearman scores of the best performing models and the baseline models, for each dataset split. The models based on randomly generated protein and/or compound representations have lower performance scores on the fully-dissimilar-split dataset, which is mainly due to the absence of identical proteins and compounds (or ones with high similarity) in between train and test samples. One of the baseline models included in this analysis uses only compound representations (i.e., only-ecfp4 model). This model does not utilize a protein vector. As a result, the model learns activities over the compound features only, without any information regarding which protein this compound interacts with. This is different from a conventional ligand-based DTI prediction model, in which target proteins would be used as labels of the input compounds (i.e., as “a target of protein X” or “not a target of protein X”). Here, since the information about proteins is not utilized at all, the model tries to learn interactions blindly, and make predictions without knowing which target it is giving predictions for.

**Table 2.** Protein family-based average Spearman scores of the best models and baseline models in each dataset split.

| Name of the descriptor set/representation<br>(explanation) | Fully-dissimilar-split | Dissimilar-compound-split | Random-split |
| --- | --- | --- | --- |
| **Best performing protein representation**<br>(compound: ECFP4) | 0.363 | 0.518 | 0.868 |
| **random200** (protein: random continuous vectors,<br>compound: ECFP4) | 0.193 | 0.436 | 0.861 |
| **only-ecfp4** (no protein vector, compound side: ECFP4) | 0.302 | 0.379 | 0.709 |
| **random200-random-ecfp4** (protein: random continuous<br>vectors, compound: random binary vectors) | 0.056 | 0.272 | 0.504 |
| **only-random-ecfp4** (no protein vector, compound side:<br>random binary vectors) | 0.002 | -0.002 | 0.315 |

The average Spearman correlation score of the best performing model on the fully-dissimilar-split dataset is around 0.3, which is quite close to the only-ecfp4 model. This indicates that the success obtained by even the best model has mostly originated from the characteristics of compounds (i.e., a certain compound being active no matter which target it has been screened against, or another compound being inactive in most of the experiments). Thus, these results reveal the requirement for; (i) unbiased model training datasets, and (ii) novel/improved featurization techniques, to construct robust DTI prediction models that can be utilized in the pharmaceutical industry, especially under these challenging scenarios.

Model performances are higher on the dissimilar-compound-split dataset compared to the fully-dissimilar-split dataset, due to the inclusion of similar (and identical) proteins between training and test. Also, models based on completely random vectors (on both the compound and protein sides) have lower performances, expectedly. On both of the challenging datasets, the best model is well differentiated from the random vector-based baseline models. Although the overall mean difference between the best model and random200 model is considerably low on the dissimilar-compound-split, the differences are distinct when making protein family-specific comparisons rather than taking the average of all families (e.g., for ion channels; the average Spearman score of the top performing models including k-sep_pssm, pfam, taap, and protvec is 0.52, and the Spearman score of random200 model is 0.37). On the dissimilar-compound-split dataset, random200 model outperformed the only-ecfp4 model by learning the relationship between the bioactivity data points of the same proteins which are shared between training and test. As experimental bioactivity measurements are mainly obtained from target-based assays, the number of bioactivity data points per protein is considerably high, compared to the number of bioactivity data points per compound (Table S3 and S4). Also, in many assays, different derivatives of the same compound are tested, which result in similar bioactivity values. Due to this bias in experimental assays, memorization over protein identity yields falsely successful results, as reflected in the performance of the random200 model on the dissimilar-compound-split dataset (average Spearman score = 0.436).

On the random-split dataset, the best model displays a high success rate (Spearman score: 0.868). However, high performance scores of the baseline models, including those based on randomly generated vectors (e.g., random200), clearly indicates the overoptimistic evaluation, and emphasizes the importance of train-test data splitting, once again. These results also demonstrate the importance of baseline model-based investigation in the field of DTI prediction, for a fair and realistic performance evaluation. It is possible to state that, the results reported in previous DTI prediction studies in which (i) the models are only evaluated based on random splitting (including both hold-out testing and fold-based cross validation), and (ii) there is no proper baseline model comparisons, may be invalid.

#### Exploration of the prediction similarities between family-specific PCM models

In this experiment, we plotted heatmaps based on pairwise similarities between the protein family-specific PCM model predictions via calculating their intersections, using a categorization composed of six classes (i.e., pChEMBL value bins of <5, 5.0 to 5.5, 5.5 to 6.0, 6.0 to 6.5, 6.5 to 7.0, and 7.0>=). To calculate the similarity between a pair of models, for each bioactivity data point, we count a similar prediction if both models predict pChEMBL values in the same bin (no matter they are correct or not), otherwise we count a non-similar prediction. We then calculate percent similarity values based on all counts. To emphasize prediction similarity values between model pairs, color scales were arranged so that the darkest color corresponds to the maximum value, and the lightest color was set to 85%, 65%, and 20% similarity for the random-split, dissimilar-compound-split, and the fully-dissimilar-split datasets, respectively.

In Figure 12, heatmaps of transferase and ion channel families are given for all three dataset splits (heatmaps for the remaining families are available at ESI Figure S3). As observed from Figure 12, the overall consensus between models decreases with increasing difficulty levels (i.e., the average similarity is over 80% for most of the models in the random-split dataset, while this value drops to 30-60% in the fully-dissimilar-split dataset). Although clusters vary across different splits and protein families, generally the learned embeddings and physicochemistry-based conventional descriptors are clustered among themselves. Considering the fully-dissimilar-split dataset of transferases; the average prediction similarity between the models that utilize learned representations (except protvec) is 60.8%, and among the models that use physicochemistry-based conventional descriptor sets (i.e., qso, apaac, geary, ctriad) is 68.2%, whereas the average prediction similarity between the physicochemistry-based conventional vs. learned representations (considering the same models) is 46.5%. These findings are also parallel to the t-SNE projection results provided in Figure 2. Considering the type of utilized information, all learned representations exploit the arrangement of amino acids on the protein sequence. On the other hand, physicochemistry-based descriptors aggregate pre-calculated amino acid-based features to construct protein feature vectors. This difference is also reflected in their prediction similarities. Spmap and random200 representations are often clustered together and have similar t-SNE projections, as well. Finally, models that utilize pfam and taap descriptor sets are quite differentiated from the rest on the random-split and dissimilar-compound-split datasets, which is expected based on their distinct featurization strategies.

**Figure 12.**
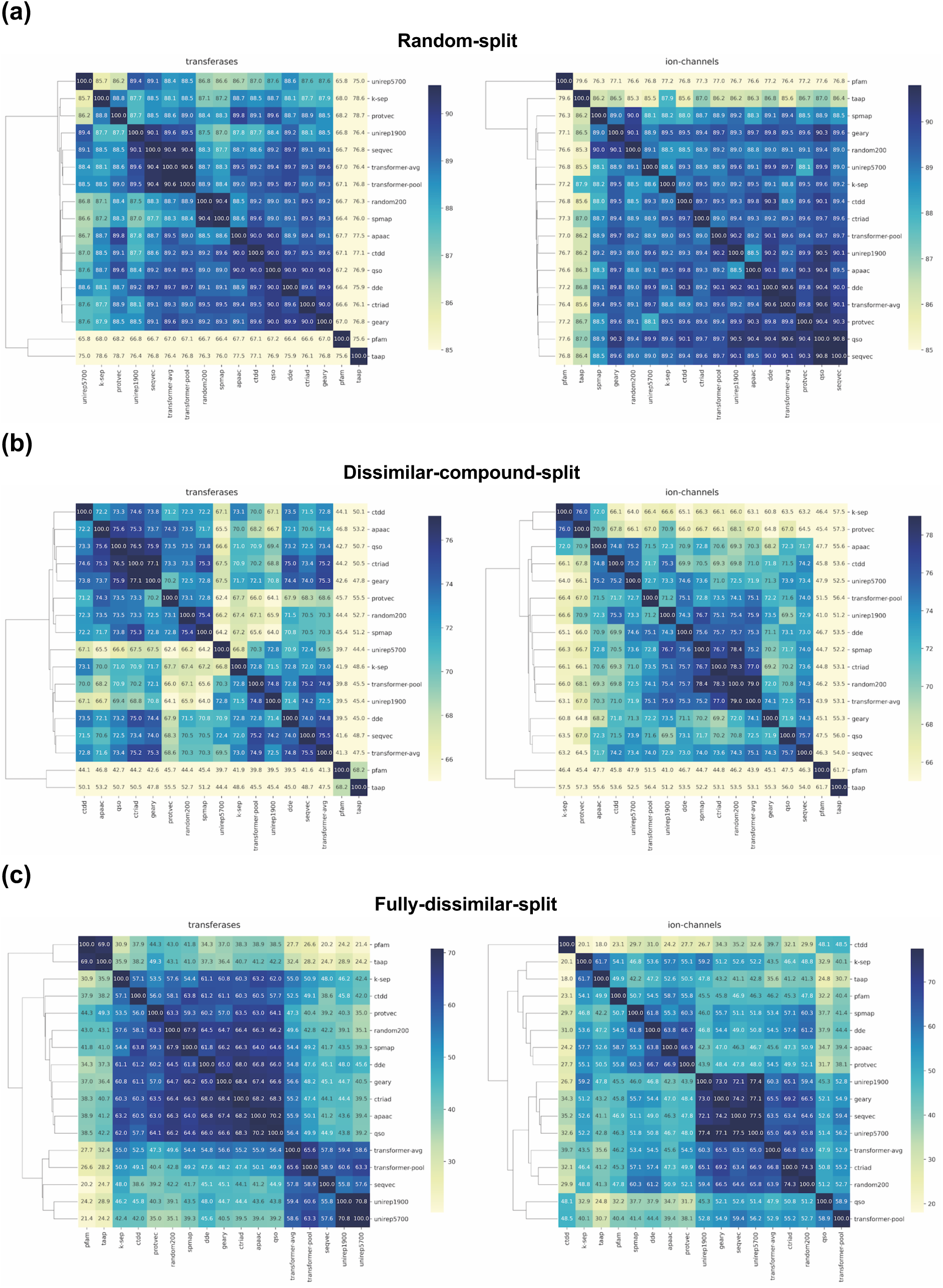
Clustered heatmaps of different protein featurization approaches for transferase and ion channel families on; **(a)** the random-split, **(b)** dissimilar-compound-split, and **(c)** the fully-dissimilar-split datasets.

The results of this analysis can be used to obtain rational combinations of featurization approaches to better represent proteins in DTI prediction models (e.g., concatenating feature vectors that have a low correct prediction overlap). This may yield a more successful learning of interaction-relevant properties of proteins, and significantly improve the overall model performances.

## 4. Conclusion

In this study, we performed a rigorous benchmark analysis to investigate; (i) bioactivity datasets at different scales and their splitting into train-test folds, (ii) preliminary and explanatory analysis of data, (iii) different modeling and algorithmic approaches, (iv) the representation capability of various protein featurization techniques, and (v) robust and fair performance evaluation strategies, for machine learning-based DTI prediction modelling. For this, we built target feature-based and PCM-based models, and trained/tested them on carefully constructed datasets with varying sizes and difficulty levels, using numerous protein representations, and evaluated them from different perspectives. Datasets, results and the source of the study is fully shared in our “ProtBENCH” platform at https://github.com/HUBioDataLab/ProtBENCH.

Below, we summarized the major contributions of our study to the literature:

*(i)* We proposed challenging benchmark datasets with high coverage on both compound and protein spaces that can be used as reliable, reference/gold-standard datasets for DTI modelling tasks. These datasets are protein family-specific, and each has three versions in terms of train/test splits for different prediction tasks (i.e., random split for predicting known inhibitors for known targets, dissimilar-compound split for predicting novel inhibitors for known targets, and fully-dissimilar split for predicting new inhibitors for new targets). Thus, they yield fair evaluation of models at multiple difficulty levels and facilitate the prevention of over-optimistic performance results. We evaluated these datasets in the framework of PCM modeling, which is a highly promising data-driven approach for high performance ML-based drug discovery. These datasets can be used in future studies to evaluate newly proposed modeling and/or algorithmic techniques for DTI prediction.
*(ii)* We employed a network-based strategy for splitting data into train-test folds, by considering both protein-protein and compound-compound pairwise similarities, which is proposed here for the first time, according to our knowledge. This strategy ensures that train and test folds are totally dissimilar from each other with a minimum loss of data points. One of the current limitations in drug development is the problems related to discovering novel molecules that are structurally different from existing drugs and drug candidates. The network-based splitting strategy we applied here forces prediction models to face this limitation by supplying more realistic, hard-to-predict test samples. Hence, it can aid researchers in designing more powerful and robust DTI prediction models that have a real translational value.
*(iii)* Protein representation learning have a wide range of applications with promising results in different sub-fields of protein science, despite being a relatively new approach. However, the studies regarding their usage in DTI prediction modelling are limited, and there is no comprehensive benchmark study to evaluate their performance against well-known and widely used featurization approaches. Due to this reason, we extended the scope of our study by involving state-of-the-art learned representations and discussed their potential in DTI prediction.

One of the critical observations of this study is the dramatic change in performance scores when the samples are distributed to train and test sets differently, (i.e., scores on datasets with challenging splits are significantly lower compared to the results on randomly split datasets), which highlights the importance of data splitting to conduct realistic evaluations for drug and/or target discovery. This study also emphasizes the importance of exploratory analysis of datasets and the usage of multiple scoring metrics as well as the inclusion of baseline models for a proper discussion of model successes.

Regarding the performance-based comparison of different protein featurization approaches, it is not possible to put forward an outstanding representation method, as their success largely depends on the dataset and the applied splitting strategy. Although both conventional descriptor sets and learned embeddings have their own strengths and weaknesses depending on the case, competitive results of learned embeddings display their potential wide-spread utilization in drug discovery and development in the near future. On the other hand, considerably low performance results on challenging datasets (e.g., fully-dissimilar-split) in the overall evaluation revealed the requirement of unbiased bioactivity datasets and further improved protein representation techniques to capture hidden and complex features shared between highly distant homologs.

As future work, we plan to develop a new computational DTI modeling approach that utilizes numerous types of biological and biomedical entities on top of compounds and target proteins. Drug discovery and development is composed of a series of complex problems, and there are multiple factors affecting the success of a drug candidate. This is mainly due to the extremely dynamic and complicated structure of biological systems. As a result, it is not possible to computationally handle drug discovery solely by simple virtual screening. Considering this fact, taking a systems-based approach with the integration and utilization of direct and indirect relationships in molecular and cellular processes including protein-protein interactions, drug/compound-target protein interactions, and signaling/metabolic pathways, together with high level concepts such as protein-disease relationships, drug-disease indications, pathway-disease modulations, and phenotypic implications could increase the success rate in drug discovery. Thus, we aim to construct a new type of systems-level DTI representation and subsequent prediction framework, using CROssBAR ^76^ which is an open-source system that integrates large-scale biological/biomedical data and represents them in the form of heterogeneous and computable knowledge graphs. The newly proposed framework will utilize graph representation learning algorithms to process these biomedical knowledge graphs, and will be trained, validated/optimized, and tested on our realistic and challenging datasets.

We hope that the results of this study, together with the data-driven approaches proposed, and the benchmark datasets prepared and shared, will aid the ongoing work in computational drug discovery and repurposing.

