## Supplementary Information Document for "How to Approach Machine Learning-based Prediction of Drug/Compound-Target Interactions"

### Supplementary Tables

**Table S1.** Statistics of the compound-centric datasets used in the small-scale analysis.

| Name of the center compound | ChEMBL id of the center compounds | cluster size (total # of compounds) | # of active targets | # of inactive targets |
|---|---|---|---|---|
| Curcumin | CHEMBL116438 | 1648 | 94 | 66 |
| Tamoxifen | CHEMBL83 | 754 | 147 | 35 |
| Quercetin | CHEMBL50 | 543 | 92 | 64 |
| Genistein | CHEMBL44 | 479 | 102 | 57 |
| Econazole | CHEMBL808 | 246 | 54 | 23 |
| Levoketoconazole | CHEMBL295698 | 243 | 79 | 27 |
| Amiodarone | CHEMBL633 | 194 | 73 | 28 |
| Miconazole | CHEMBL91 | 170 | 49 | 22 |
| Clotrimazole | CHEMBL104 | 52 | 38 | 37 |

**Table S2.** Enzyme sub-classes of the protein family specific datasets used in the large-scale analysis.

| Main class EC number(s) of the merged enzyme classes | Names of merged enzyme classes |
|---|---|
| 1 | Oxidoreductases, Cytochrome P450s |
| 2 | Transferases, Kinases |
| 3 (except 3.4) | Hydrolases, Phosphatases, Phosphodiesterases |
| 3.4 | Proteases (a.k.a. Peptidases) |
| All remaining | Other, Lyases |

**Table S3.** Statistics of the protein family specific datasets used in the large-scale analysis.

| | Datapoints | | Proteins | | Compounds | |
|---|---|---|---|---|---|---|
| **Protein family** | **initial** | **discarded** | **initial** | **discarded** | **initial** | **discarded** |
| epigenetic regulators | 19,219 | 1,055 | 125 | 8 | 10,637 | 276 |
| hydrolases | 71,996 | 9,051 | 623 | 12 | 56,313 | 7,282 |
| ion channels | 38,097 | 3,865 | 241 | 26 | 32,376 | 2,073 |
| membrane receptors | 220,810 | 14,635 | 640 | 3 | 133,440 | 8,357 |
| other enzymes | 36,857 | 4,007 | 456 | 9 | 21,428 | 3,372 |
| oxidoreductases | 59,023 | 4,545 | 404 | 2 | 42,338 | 2,895 |
| proteases | 99,100 | 11,839 | 409 | 20 | 64,688 | 8,369 |
| transcription factors | 25,292 | 216 | 119 | 0 | 17,244 | 80 |
| transferases | 191,864 | 31,900 | 1,020 | 39 | 124,266 | 18,963 |
| transporters | 29,165 | 804 | 190 | 2 | 18,493 | 523 |

**Table S4.** Split-based statistics of the protein family specific datasets for; **(a)** the random-split, **(b)** dissimilar-compound-split, and **(c)** the fully-dissimilar-split sets (ratio represents the ratio of train to test).

**(a)**

| Random-split | Bioactivity datapoints | | | Proteins | | | Compounds | | |
|---|---|---|---|---|---|---|---|---|---|
| Protein family | train | test | ratio | train | test | ratio | train | test | ratio |
| epigenetic-regulators | 16,675 | 1,489 | 11.2 | 117 | 83 | 1.4 | 9,797 | 1,388 | 7.1 |
| hydrolases | 59,021 | 3,924 | 15.0 | 610 | 304 | 2.0 | 46,575 | 3,851 | 12.1 |
| ion-channels | 31,287 | 2,945 | 10.6 | 213 | 142 | 1.5 | 27,961 | 2,898 | 9.6 |
| membrane-receptors | 197,919 | 8,256 | 24.0 | 636 | 428 | 1.5 | 121,954 | 8,030 | 15.2 |
| other-enzymes | 29,465 | 3,385 | 8.7 | 442 | 256 | 1.7 | 16,757 | 3,040 | 5.5 |
| oxidoreductases | 52,277 | 2,201 | 23.8 | 402 | 240 | 1.7 | 38,277 | 2,169 | 17.6 |
| proteases | 83,561 | 3,700 | 22.6 | 389 | 219 | 1.8 | 54,628 | 3,598 | 15.2 |
| transcription-factors | 23,498 | 1,578 | 14.9 | 118 | 80 | 1.5 | 16,465 | 1,529 | 10.8 |
| transferases | 150,247 | 9,717 | 15.5 | 980 | 634 | 1.5 | 100,462 | 9,070 | 11.1 |
| transporters | 26,313 | 2,048 | 12.8 | 188 | 114 | 1.6 | 17,141 | 1,958 | 8.8 |

**(b)**

| Dissimilar-compound-split | Bioactivity datapoints | | | Proteins | | | Compounds | | |
|---|---|---|---|---|---|---|---|---|---|
| Protein family | train | test | ratio | train | test | ratio | train | test | ratio |
| epigenetic-regulators | 16,763 | 1,401 | 12.0 | 117 | 52 | 2.3 | 9,536 | 825 | 11.6 |
| hydrolases | 58,953 | 3,992 | 14.8 | 597 | 178 | 3.4 | 45,913 | 3,118 | 14.7 |
| ion-channels | 31,329 | 2,903 | 10.8 | 208 | 69 | 3.0 | 27,656 | 2,647 | 10.4 |
| membrane-receptors | 197,652 | 8,523 | 23.2 | 627 | 252 | 2.5 | 118,914 | 6,169 | 19.3 |
| other-enzymes | 29,325 | 3,525 | 8.3 | 433 | 157 | 2.8 | 15,428 | 2,628 | 5.9 |
| oxidoreductases | 52,190 | 2,288 | 22.8 | 396 | 107 | 3.7 | 37,763 | 1,680 | 22.5 |
| proteases | 83,405 | 3,856 | 21.6 | 383 | 130 | 2.9 | 53,493 | 2,826 | 18.9 |
| transcription-factors | 23,444 | 1,632 | 14.4 | 116 | 38 | 3.1 | 16,015 | 1,149 | 13.9 |
| transferases | 150,216 | 9,748 | 15.4 | 956 | 393 | 2.4 | 98,103 | 7,200 | 13.6 |
| transporters | 26,273 | 2,088 | 12.6 | 182 | 63 | 2.9 | 16,545 | 1,425 | 11.6 |

**(c)**

| Fully-dissimilar-split | Bioactivity datapoints | | | Proteins | | | Compounds | | |
|---|---|---|---|---|---|---|---|---|---|
| Protein family | train | test | ratio | train | test | ratio | train | test | ratio |
| epigenetic-regulators | 16,675 | 1,489 | 11.2 | 117 | 52 | 2.3 | 9,536 | 825 | 11.6 |
| hydrolases | 59,021 | 3,924 | 15.0 | 597 | 178 | 3.4 | 45,913 | 3,118 | 14.7 |
| ion-channels | 31,287 | 2,945 | 10.6 | 208 | 69 | 3.0 | 27,656 | 2,647 | 10.4 |
| membrane-receptors | 197,919 | 8,256 | 24.0 | 627 | 252 | 2.5 | 118,914 | 6,169 | 19.3 |
| other-enzymes | 29,465 | 3,385 | 8.7 | 433 | 157 | 2.8 | 15,428 | 2,628 | 5.9 |
| oxidoreductases | 52,277 | 2,201 | 23.8 | 396 | 107 | 3.7 | 37,763 | 1,680 | 22.5 |
| proteases | 83,561 | 3,700 | 22.6 | 383 | 130 | 2.9 | 53,493 | 2,826 | 18.9 |
| transcription-factors | 23,498 | 1,578 | 14.9 | 116 | 38 | 3.1 | 16,015 | 1,149 | 13.9 |
| transferases | 150,247 | 9,717 | 15.5 | 956 | 393 | 2.4 | 98,103 | 7,200 | 13.6 |
| transporters | 26,313 | 2,048 | 12.8 | 182 | 63 | 2.9 | 16,545 | 1,425 | 11.6 |

**Table S5.** Model performance scores (in terms of MCC) in the small-scale analysis (on the compound-centric datasets) for; **(a)** random forest, and **(b)** SVM models. The 3 best performances for each dataset are shown in bold font. For the results based on other performance metrics please see the supplementary spreadsheets attached as external files named: "small-scale_rf_overall_test_results.xlsx" and "small-scale_svm_overall_test_results.xlsx".

**(a)**

| Model | ChEMBL id (only the numeric part) of the center compound of each compound cluster | | | | | | | | | Mean | Standard error |
|---|---|---|---|---|---|---|---|---|---|---|---|
| | 44 | 50 | 83 | 91 | 104 | 633 | 808 | 116438 | 295698 | | |
| aac | 0.283 | 0.112 | 0.359 | 0.238 | 0.413 | 0.199 | 0.227 | 0.099 | 0.195 | 0.236 | 0.035 |
| aac_pssm | 0.226 | 0.278 | 0.395 | 0.432 | 0.170 | 0.212 | 0.399 | 0.153 | 0.270 | 0.282 | 0.035 |
| aadp_pssm | 0.166 | 0.211 | 0.396 | 0.453 | 0.177 | 0.347 | 0.330 | 0.240 | 0.380 | 0.300 | 0.035 |
| aatp_pssm | 0.231 | 0.312 | 0.432 | 0.477 | 0.332 | 0.280 | 0.398 | 0.254 | 0.387 | 0.345 | 0.028 |
| ab_pssm | 0.237 | 0.231 | 0.397 | 0.408 | 0.157 | 0.323 | 0.487 | 0.206 | 0.364 | 0.312 | 0.037 |
| apaac | 0.263 | 0.262 | 0.463 | 0.346 | 0.303 | 0.514 | 0.422 | **0.293** | 0.384 | 0.361 | 0.030 |
| cksaagp | 0.243 | 0.034 | 0.390 | 0.333 | 0.222 | 0.323 | 0.261 | 0.120 | 0.260 | 0.243 | 0.037 |
| cksaap | 0.324 | 0.268 | 0.510 | 0.410 | 0.297 | 0.489 | 0.405 | 0.143 | 0.431 | 0.364 | 0.039 |
| ctdc | 0.117 | -0.035 | 0.361 | 0.213 | 0.332 | 0.280 | 0.223 | 0.185 | 0.109 | 0.199 | 0.041 |
| ctdd | 0.176 | 0.209 | 0.446 | 0.204 | 0.142 | 0.512 | 0.471 | 0.185 | 0.390 | 0.304 | 0.049 |
| ctdt | 0.212 | 0.033 | 0.309 | 0.201 | 0.363 | 0.169 | 0.379 | 0.130 | 0.301 | 0.233 | 0.038 |
| ctriad | 0.231 | 0.179 | 0.355 | 0.230 | 0.243 | 0.449 | 0.285 | 0.280 | 0.396 | 0.294 | 0.029 |
| d_fpssm | **0.364** | 0.288 | 0.429 | 0.371 | 0.398 | 0.518 | 0.492 | 0.250 | 0.345 | 0.384 | 0.029 |
| dde | 0.302 | 0.293 | **0.528** | **0.492** | 0.356 | 0.512 | **0.536** | 0.174 | 0.446 | **0.404** | 0.043 |
| dp_pssm | 0.279 | **0.376** | 0.369 | 0.387 | 0.297 | 0.316 | 0.325 | 0.278 | 0.396 | 0.336 | 0.016 |
| dpc | 0.318 | 0.170 | 0.467 | 0.482 | 0.197 | 0.453 | 0.395 | 0.121 | 0.412 | 0.335 | 0.046 |
| dpc_pssm | 0.227 | 0.210 | 0.396 | 0.491 | 0.198 | 0.369 | 0.360 | 0.233 | 0.433 | 0.324 | 0.036 |
| edp_pssm | 0.144 | 0.172 | 0.375 | 0.388 | 0.164 | 0.141 | 0.358 | 0.152 | 0.273 | 0.241 | 0.036 |
| eedp_pssm | 0.186 | 0.225 | 0.419 | 0.433 | 0.164 | 0.310 | 0.336 | 0.022 | 0.262 | 0.262 | 0.043 |
| gaac | 0.089 | -0.081 | 0.266 | 0.020 | 0.088 | 0.200 | 0.135 | -0.057 | 0.129 | 0.088 | 0.038 |
| gdpc | 0.314 | 0.045 | 0.311 | 0.292 | 0.253 | 0.310 | 0.262 | 0.076 | 0.232 | 0.233 | 0.034 |
| geary | 0.337 | 0.182 | 0.390 | 0.394 | 0.294 | 0.339 | 0.360 | **0.345** | 0.322 | 0.329 | 0.021 |
| gtpc | 0.304 | 0.134 | 0.344 | 0.198 | 0.122 | 0.442 | 0.339 | 0.184 | 0.404 | 0.275 | 0.039 |
| k-sep_pssm | 0.330 | 0.312 | **0.561** | 0.326 | **0.452** | 0.506 | 0.313 | 0.209 | 0.355 | 0.374 | 0.037 |
| ksctriad | 0.295 | 0.190 | 0.394 | 0.268 | 0.316 | 0.448 | 0.315 | 0.172 | 0.387 | 0.310 | 0.031 |
| medp_pssm | 0.167 | 0.164 | 0.417 | 0.431 | 0.216 | 0.309 | 0.375 | 0.023 | 0.285 | 0.265 | 0.045 |
| moran | 0.325 | 0.236 | 0.367 | 0.391 | 0.264 | 0.412 | 0.415 | 0.247 | 0.377 | 0.337 | 0.024 |
| nmbroto | 0.254 | 0.190 | 0.365 | 0.444 | 0.277 | 0.327 | 0.344 | 0.195 | 0.391 | 0.310 | 0.029 |
| paac | 0.230 | 0.181 | 0.502 | 0.417 | 0.255 | 0.505 | 0.500 | 0.262 | 0.347 | 0.356 | 0.043 |
| pfam | 0.325 | 0.286 | 0.486 | 0.432 | 0.430 | 0.203 | 0.273 | 0.132 | 0.399 | 0.329 | 0.039 |
| pse_pssm | 0.209 | 0.267 | 0.438 | 0.327 | 0.386 | 0.228 | 0.340 | **0.314** | 0.396 | 0.323 | 0.026 |
| pssm_ac | **0.371** | **0.379** | 0.393 | 0.426 | 0.324 | **0.578** | 0.451 | 0.210 | **0.478** | **0.401** | 0.034 |
| pssm_cc | 0.293 | 0.276 | 0.401 | 0.413 | 0.302 | 0.483 | 0.400 | 0.274 | 0.398 | 0.360 | 0.025 |
| pssm_composition | 0.220 | 0.221 | 0.455 | 0.451 | 0.333 | 0.373 | **0.507** | 0.142 | 0.437 | 0.349 | 0.043 |
| qso | 0.229 | 0.247 | 0.410 | 0.224 | 0.378 | 0.433 | 0.291 | 0.141 | 0.348 | 0.300 | 0.033 |
| random200 | 0.071 | 0.080 | -0.035 | -0.033 | -0.046 | -0.100 | -0.055 | -0.111 | 0.054 | -0.019 | 0.024 |
| rpm_pssm | 0.289 | 0.132 | 0.401 | 0.471 | **0.440** | 0.410 | 0.358 | 0.189 | 0.422 | 0.346 | 0.039 |
| rpssm | 0.055 | 0.197 | 0.355 | 0.456 | 0.187 | 0.340 | 0.418 | 0.087 | 0.338 | 0.270 | 0.048 |
| spmap | 0.338 | 0.259 | 0.431 | 0.405 | 0.194 | **0.556** | 0.489 | 0.153 | **0.457** | 0.365 | 0.046 |
| taap | 0.174 | 0.179 | 0.455 | 0.283 | **0.431** | 0.442 | 0.402 | 0.102 | 0.309 | 0.309 | 0.044 |
| tpc | 0.245 | 0.268 | 0.476 | **0.499** | 0.270 | 0.529 | **0.528** | 0.124 | **0.448** | 0.376 | 0.050 |
| tpc_pssm | 0.220 | 0.263 | 0.330 | **0.492** | 0.352 | 0.241 | 0.356 | 0.241 | 0.412 | 0.323 | 0.030 |
| tri-gram_pssm | **0.350** | **0.349** | **0.541** | 0.362 | 0.426 | **0.545** | 0.386 | 0.274 | 0.352 | **0.398** | 0.030 |

**(b)**

| Model | ChEMBL id (only the numeric part) of the center compound of each compound cluster | | | | | | | | | Mean | Standard error |
|---|---|---|---|---|---|---|---|---|---|---|---|
| | 44 | 50 | 83 | 91 | 104 | 633 | 808 | 116438 | 295698 | | |
| aac | 0.122 | 0.049 | 0.183 | 0.102 | 0.243 | 0.279 | 0.091 | 0.161 | 0.136 | 0.152 | 0.025 |
| aac_pssm | 0.178 | 0.146 | 0.396 | **0.488** | 0.285 | 0.306 | 0.378 | 0.144 | 0.242 | 0.285 | 0.040 |
| aadp_pssm | 0.195 | 0.142 | 0.360 | 0.287 | 0.218 | 0.254 | 0.394 | 0.132 | 0.359 | 0.260 | 0.032 |
| aatp_pssm | 0.096 | 0.180 | 0.390 | 0.392 | 0.311 | 0.305 | 0.471 | 0.132 | 0.262 | 0.282 | 0.042 |
| ab_pssm | 0.296 | 0.311 | 0.385 | 0.372 | 0.201 | 0.470 | 0.370 | 0.195 | 0.295 | 0.322 | 0.030 |
| apaac | 0.159 | 0.292 | **0.508** | 0.430 | 0.358 | **0.559** | 0.438 | 0.200 | 0.300 | 0.360 | 0.045 |
| cksaagp | 0.215 | 0.045 | 0.238 | 0.248 | 0.240 | 0.216 | 0.146 | 0.229 | 0.283 | 0.207 | 0.024 |
| cksaap | 0.329 | 0.260 | 0.336 | 0.380 | 0.433 | 0.316 | **0.491** | 0.164 | **0.506** | 0.357 | 0.036 |
| ctdc | 0.197 | -0.055 | 0.280 | 0.225 | 0.353 | 0.158 | 0.134 | 0.079 | 0.121 | 0.166 | 0.039 |
| ctdd | 0.151 | 0.226 | 0.375 | 0.357 | 0.225 | 0.310 | 0.414 | 0.237 | 0.024 | 0.258 | 0.041 |
| ctdt | 0.229 | -0.013 | 0.233 | 0.099 | **0.498** | 0.300 | 0.284 | 0.096 | 0.053 | 0.198 | 0.052 |
| ctriad | 0.171 | 0.174 | 0.293 | 0.215 | 0.408 | 0.329 | 0.233 | 0.164 | 0.271 | 0.251 | 0.027 |
| d_fpssm | 0.205 | 0.123 | 0.242 | 0.388 | 0.025 | 0.140 | 0.394 | 0.033 | 0.362 | 0.212 | 0.048 |
| dde | 0.307 | 0.301 | 0.360 | 0.283 | 0.389 | 0.386 | 0.358 | 0.238 | 0.409 | 0.337 | 0.019 |
| dp_pssm | 0.228 | **0.434** | 0.370 | 0.363 | 0.229 | 0.329 | 0.335 | 0.186 | 0.313 | 0.310 | 0.027 |
| dpc | 0.202 | 0.106 | 0.308 | 0.291 | 0.236 | 0.333 | 0.263 | 0.193 | 0.337 | 0.252 | 0.025 |
| dpc_pssm | 0.195 | 0.142 | 0.418 | 0.287 | 0.218 | 0.254 | 0.394 | 0.132 | 0.358 | 0.266 | 0.035 |
| edp_pssm | 0.131 | 0.253 | 0.391 | 0.331 | 0.215 | 0.256 | 0.262 | 0.004 | 0.334 | 0.242 | 0.039 |
| eedp_pssm | 0.220 | 0.248 | 0.373 | 0.415 | 0.080 | 0.266 | **0.491** | 0.059 | 0.283 | 0.271 | 0.048 |
| gaac | 0.151 | 0.067 | 0.172 | 0.120 | -0.089 | 0.230 | 0.111 | 0.065 | 0.206 | 0.115 | 0.032 |
| gdpc | 0.088 | 0.136 | 0.273 | 0.319 | 0.055 | 0.134 | 0.018 | 0.139 | 0.234 | 0.155 | 0.034 |
| geary | 0.277 | 0.201 | 0.357 | 0.327 | 0.308 | 0.288 | 0.280 | 0.195 | 0.358 | 0.288 | 0.020 |
| gtpc | 0.140 | 0.056 | 0.135 | 0.060 | 0.105 | 0.212 | 0.243 | **0.259** | 0.362 | 0.175 | 0.034 |
| k-sep_pssm | 0.277 | 0.347 | **0.513** | 0.288 | 0.418 | 0.474 | 0.456 | 0.241 | 0.312 | **0.370** | 0.033 |
| ksctriad | 0.124 | 0.231 | 0.406 | 0.277 | 0.031 | 0.402 | 0.338 | 0.148 | 0.262 | 0.247 | 0.043 |
| medp_pssm | 0.198 | 0.270 | 0.365 | 0.415 | 0.080 | 0.280 | **0.491** | 0.091 | 0.271 | 0.274 | 0.046 |
| moran | 0.264 | 0.215 | 0.345 | 0.347 | 0.181 | 0.315 | 0.384 | 0.184 | 0.303 | 0.282 | 0.025 |
| nmbroto | 0.231 | 0.228 | 0.354 | 0.289 | 0.439 | 0.282 | 0.386 | 0.236 | 0.258 | 0.300 | 0.025 |
| paac | 0.257 | 0.268 | 0.480 | 0.314 | 0.246 | **0.528** | 0.443 | 0.167 | 0.338 | 0.338 | 0.040 |
| pfam | 0.288 | 0.328 | 0.478 | **0.452** | 0.327 | 0.353 | 0.432 | **0.367** | 0.398 | **0.380** | 0.021 |
| pse_pssm | 0.248 | 0.280 | 0.367 | 0.334 | 0.144 | 0.348 | 0.342 | 0.176 | 0.343 | 0.287 | 0.027 |
| pssm_ac | 0.331 | **0.353** | 0.290 | 0.392 | 0.351 | 0.423 | 0.357 | 0.164 | **0.428** | 0.343 | 0.027 |
| pssm_cc | 0.294 | **0.368** | 0.484 | 0.311 | **0.517** | 0.446 | 0.373 | 0.192 | 0.349 | **0.370** | 0.034 |
| pssm_composition | 0.154 | 0.204 | 0.386 | 0.411 | 0.257 | 0.419 | 0.376 | 0.114 | 0.348 | 0.297 | 0.039 |
| qso | 0.220 | 0.078 | 0.249 | 0.211 | 0.346 | 0.268 | 0.073 | 0.213 | 0.113 | 0.197 | 0.031 |
| random200 | -0.044 | -0.054 | -0.110 | -0.206 | -0.046 | -0.083 | 0.248 | -0.140 | 0.205 | -0.026 | 0.051 |
| rpm_pssm | 0.331 | 0.177 | 0.376 | 0.426 | 0.300 | 0.430 | 0.435 | 0.062 | 0.377 | 0.324 | 0.042 |
| rpssm | 0.133 | 0.312 | 0.260 | **0.431** | 0.249 | 0.305 | 0.361 | 0.076 | 0.361 | 0.276 | 0.038 |
| spmap | **0.342** | 0.260 | 0.486 | 0.254 | 0.249 | **0.492** | 0.394 | 0.026 | 0.370 | 0.319 | 0.048 |
| taap | 0.211 | 0.284 | 0.456 | 0.325 | 0.210 | 0.408 | 0.337 | 0.167 | 0.289 | 0.299 | 0.032 |
| tpc | **0.356** | 0.218 | 0.417 | 0.354 | 0.300 | 0.419 | 0.269 | 0.239 | 0.418 | 0.332 | 0.026 |
| tpc_pssm | 0.111 | 0.260 | 0.285 | 0.232 | 0.237 | 0.276 | 0.264 | 0.209 | **0.425** | 0.256 | 0.027 |
| tri-gram_pssm | **0.340** | 0.309 | **0.487** | 0.368 | **0.444** | 0.485 | 0.490 | **0.284** | 0.331 | **0.393** | 0.028 |

**Table S6.** Model performance scores in the medium-scale analysis (on the mDavis dataset). The best performance for each metric is shown in bold font.

| **Model** | **RMSE** | **Spearman** | **F1-score** | **MCC** |
|---|---|---|---|---|
| seqvec | **0.794** | **0.571** | 0.530 | **0.445** |
| k-sep_pssm | 0.817 | 0.545 | **0.531** | 0.434 |
| unirep1900 | 0.823 | 0.541 | 0.510 | 0.418 |
| apaac | 0.831 | 0.532 | 0.519 | 0.418 |
| unirep5700 | 0.831 | 0.531 | 0.506 | 0.412 |
| transformer-avg | 0.839 | 0.519 | 0.508 | 0.410 |
| transformer-pool | 0.840 | 0.515 | 0.506 | 0.412 |
| qso | 0.843 | 0.519 | 0.486 | 0.384 |
| dde | 0.845 | 0.508 | 0.480 | 0.384 |
| geary | 0.847 | 0.519 | 0.473 | 0.377 |
| protvec | 0.850 | 0.503 | 0.506 | 0.403 |
| ctdd | 0.851 | 0.503 | 0.484 | 0.376 |
| ctriad | 0.852 | 0.508 | 0.476 | 0.387 |
| pfam | 0.854 | 0.497 | 0.538 | 0.410 |
| taap | 0.863 | 0.492 | 0.467 | 0.349 |
| spmap | 0.871 | 0.491 | 0.477 | 0.362 |
| random200 | 0.957 | 0.403 | 0.368 | 0.251 |
| random200_random-ecfp4 | 0.968 | 0.388 | 0.346 | 0.235 |

**Table S7.** Model performance scores (in terms of the median corrected MCC) in the large-scale analysis on the protein family specific datasets of; **(a)** the random-split, **(b)** dissimilar-compound-split, and **(c)** the fully-dissimilar-split. The 3 best performances for each protein family are shown in bold font (ran200_ran-ecfp4: random200_ random-ecfp4, only-ran-ecfp4: only-random-ecfp4). For the results based on other performance metrics please see the supplementary spreadsheet attached as an external file named: "large-scale_overall_test_results.xlsx".

**(a)**

| **Random-split** | **epigenetic-regulators** | **hydrolases** | **ion-channels** | **membrane-receptors** | **other-enzymes** | **oxidoreductases** | **proteases** | **transcription-factors** | **transferases** | **transporters** |
|---|---|---|---|---|---|---|---|---|---|---|
| apaac | **0.745** | 0.755 | 0.697 | 0.689 | 0.754 | 0.692 | **0.735** | 0.714 | 0.696 | 0.728 |
| ctdd | 0.741 | 0.747 | 0.700 | 0.686 | **0.757** | **0.694** | 0.730 | 0.711 | 0.694 | **0.732** |
| ctriad | 0.734 | 0.749 | 0.701 | 0.686 | 0.752 | **0.694** | 0.731 | 0.706 | 0.694 | 0.726 |
| dde | 0.741 | **0.756** | 0.703 | 0.689 | 0.754 | 0.692 | **0.735** | 0.709 | 0.691 | 0.722 |
| geary | 0.733 | **0.754** | 0.701 | **0.694** | 0.752 | 0.681 | **0.735** | **0.721** | 0.696 | 0.728 |
| k-sep_pssm | **0.757** | 0.749 | **0.709** | 0.688 | 0.754 | 0.690 | **0.735** | 0.706 | **0.704** | 0.720 |
| pfam | 0.678 | 0.694 | 0.679 | 0.458 | 0.609 | 0.561 | 0.635 | 0.645 | 0.628 | 0.622 |
| qso | 0.734 | **0.757** | 0.700 | 0.685 | 0.754 | 0.690 | 0.733 | 0.704 | 0.691 | 0.728 |
| random200 | 0.728 | 0.751 | 0.687 | 0.680 | 0.746 | 0.685 | 0.734 | 0.709 | 0.687 | 0.726 |
| spmap | 0.737 | 0.748 | 0.697 | 0.682 | **0.757** | 0.680 | 0.728 | 0.709 | 0.683 | 0.720 |
| taap | **0.760** | 0.747 | **0.712** | 0.687 | 0.750 | 0.693 | **0.736** | **0.721** | 0.700 | **0.730** |
| protvec | 0.741 | 0.742 | 0.703 | **0.693** | **0.758** | **0.696** | 0.733 | 0.714 | 0.696 | 0.726 |
| seqvec | **0.745** | 0.749 | 0.699 | 0.690 | **0.757** | 0.678 | 0.728 | 0.709 | 0.700 | 0.724 |
| transformer-avg | 0.736 | 0.748 | **0.707** | **0.691** | **0.760** | 0.681 | 0.734 | 0.701 | 0.702 | 0.718 |
| transformer-pool | 0.734 | 0.746 | 0.695 | 0.689 | 0.741 | 0.684 | 0.733 | 0.714 | 0.694 | **0.730** |
| unirep1900 | 0.744 | 0.745 | 0.703 | 0.686 | 0.753 | **0.696** | 0.731 | **0.716** | **0.703** | 0.728 |
| unirep5700 | 0.729 | 0.749 | 0.690 | 0.688 | 0.755 | 0.690 | 0.734 | 0.706 | **0.705** | 0.726 |
| only-ecfp4 | 0.591 | 0.665 | 0.643 | 0.426 | 0.600 | 0.514 | 0.519 | 0.534 | 0.576 | 0.503 |
| ran200_ran-ecfp4 | 0.382 | 0.481 | 0.400 | 0.256 | 0.449 | 0.401 | 0.320 | 0.265 | 0.319 | 0.235 |
| only-ran-ecfp4 | 0.296 | 0.175 | 0.082 | 0.165 | 0.358 | 0.137 | 0.189 | 0.171 | 0.224 | 0.173 |

**(b)**

| Dissimilar-compound-split | epigenetic-regulators | hydrolases | ion-channels | membrane-receptors | other-enzymes | oxidoreductases | proteases | transcription-factors | transferases | transporters |
|---|---|---|---|---|---|---|---|---|---|---|
| apaac | 0.137 | 0.355 | 0.342 | 0.249 | 0.419 | 0.391 | 0.381 | 0.058 | **0.358** | 0.362 |
| ctdd | 0.021 | **0.407** | 0.311 | 0.241 | 0.386 | 0.423 | **0.391** | 0.048 | 0.342 | 0.405 |
| ctriad | 0.045 | 0.351 | 0.276 | 0.243 | 0.354 | 0.405 | 0.353 | 0.006 | 0.346 | **0.425** |
| dde | 0.089 | 0.371 | 0.327 | 0.223 | 0.359 | 0.398 | 0.341 | 0.071 | 0.340 | 0.403 |
| geary | 0.044 | 0.375 | 0.291 | 0.242 | 0.397 | 0.400 | **0.394** | 0.036 | **0.347** | 0.417 |
| k-sep_pssm | 0.239 | **0.382** | 0.419 | **0.298** | 0.381 | **0.449** | 0.354 | 0.071 | 0.318 | 0.368 |
| pfam | **0.455** | 0.329 | **0.448** | 0.146 | **0.452** | 0.339 | 0.319 | **0.257** | 0.308 | 0.366 |
| qso | 0.247 | 0.373 | 0.338 | **0.278** | 0.345 | 0.356 | 0.369 | 0.071 | 0.324 | **0.419** |
| random200 | 0.152 | **0.386** | 0.273 | 0.266 | 0.348 | 0.409 | 0.341 | 0.103 | 0.341 | 0.388 |
| spmap | 0.158 | 0.351 | 0.289 | **0.274** | 0.361 | 0.395 | 0.383 | 0.036 | 0.335 | 0.369 |
| taap | **0.289** | 0.371 | **0.434** | 0.243 | **0.443** | 0.360 | **0.398** | **0.187** | 0.322 | **0.438** |
| protvec | -0.024 | 0.348 | **0.437** | 0.222 | 0.363 | 0.381 | 0.366 | **0.118** | 0.344 | 0.390 |
| seqvec | 0.192 | 0.349 | 0.310 | 0.228 | 0.374 | **0.435** | 0.373 | 0.033 | **0.359** | 0.378 |
| transformer-avg | 0.075 | 0.349 | 0.288 | 0.250 | **0.447** | **0.428** | **0.391** | 0.043 | 0.328 | 0.390 |
| transformer-pool | 0.104 | 0.348 | 0.364 | 0.258 | 0.411 | 0.402 | 0.357 | 0.061 | 0.316 | 0.376 |
| unirep1900 | 0.161 | 0.334 | 0.277 | 0.256 | 0.402 | 0.400 | 0.376 | 0.076 | 0.329 | 0.370 |
| unirep5700 | 0.061 | 0.350 | 0.289 | 0.255 | 0.387 | 0.374 | 0.346 | 0.090 | 0.320 | 0.376 |
| only-ecfp4 | **0.428** | 0.244 | 0.243 | 0.168 | 0.419 | 0.281 | 0.307 | 0.058 | 0.306 | 0.309 |
| ran200_ran-ecfp4 | -0.076 | 0.254 | 0.293 | 0.133 | 0.210 | 0.270 | 0.138 | 0.083 | 0.178 | 0.284 |
| only-ran-ecfp4 | 0.004 | -0.020 | 0.018 | 0.001 | -0.028 | -0.015 | -0.008 | 0.016 | -0.020 | -0.035 |

**(c)**

| Fully-dissimilar-split | epigenetic-regulators | hydrolases | ion-channels | membrane-receptors | other-enzymes | oxidoreductases | proteases | transcription-factors | transferases | transporters |
|---|---|---|---|---|---|---|---|---|---|---|
| apaac | 0.403 | 0.156 | 0.146 | 0.243 | 0.129 | -0.044 | 0.192 | 0.063 | 0.300 | **0.240** |
| ctdd | 0.396 | 0.132 | -0.074 | 0.253 | 0.162 | 0.074 | 0.207 | 0.101 | 0.247 | -0.017 |
| ctriad | **0.420** | **0.203** | 0.086 | 0.220 | 0.125 | 0.030 | 0.212 | **0.238** | 0.273 | **0.276** |
| dde | 0.375 | 0.170 | 0.124 | 0.206 | **0.232** | -0.050 | 0.195 | 0.150 | 0.295 | 0.029 |
| geary | 0.319 | 0.160 | 0.155 | 0.195 | 0.172 | 0.044 | **0.268** | 0.066 | 0.275 | -0.027 |
| k-sep_pssm | 0.252 | 0.181 | 0.157 | 0.137 | 0.043 | 0.052 | -0.134 | 0.086 | 0.300 | **0.297** |
| pfam | **0.446** | **0.208** | **0.174** | **0.270** | 0.221 | 0.088 | 0.142 | 0.156 | 0.301 | 0.198 |
| qso | 0.397 | 0.111 | 0.044 | 0.166 | 0.187 | 0.141 | 0.215 | 0.129 | 0.300 | 0.202 |
| random200 | 0.289 | 0.040 | 0.146 | 0.226 | **0.282** | 0.070 | 0.149 | 0.051 | 0.284 | -0.194 |
| spmap | 0.361 | 0.103 | **0.182** | 0.213 | 0.114 | 0.015 | 0.209 | 0.091 | 0.287 | 0.118 |
| taap | 0.289 | 0.155 | **0.181** | **0.286** | 0.208 | -0.028 | 0.205 | 0.129 | 0.310 | 0.194 |
| protvec | 0.275 | 0.146 | **0.174** | 0.235 | 0.131 | 0.077 | 0.184 | **0.160** | 0.301 | 0.204 |
| seqvec | 0.372 | 0.154 | 0.032 | 0.155 | 0.199 | 0.018 | 0.046 | 0.150 | 0.276 | -0.023 |
| transformer-avg | 0.367 | 0.129 | 0.058 | **0.265** | 0.176 | 0.092 | **0.227** | 0.144 | 0.311 | 0.187 |
| transformer-pool | 0.403 | 0.148 | -0.052 | 0.244 | 0.079 | **0.133** | 0.170 | 0.117 | **0.313** | -0.040 |
| unirep1900 | 0.325 | 0.186 | 0.143 | 0.217 | 0.205 | **0.132** | 0.159 | -0.008 | **0.332** | 0.004 |
| unirep5700 | 0.339 | 0.154 | 0.143 | 0.204 | 0.229 | **0.226** | 0.189 | 0.127 | **0.324** | 0.114 |
| only-ecfp4 | **0.426** | **0.205** | 0.133 | 0.254 | **0.240** | 0.016 | **0.287** | **0.169** | 0.294 | 0.172 |
| ran200_ran-ecfp4 | 0.087 | 0.067 | -0.012 | 0.022 | -0.001 | 0.123 | 0.061 | -0.003 | 0.057 | 0.032 |
| only-ran-ecfp4 | 0.041 | -0.003 | -0.013 | 0.001 | 0.010 | -0.031 | -0.019 | 0.001 | 0.018 | 0.025 |

### Supplementary Figures

**(a)**

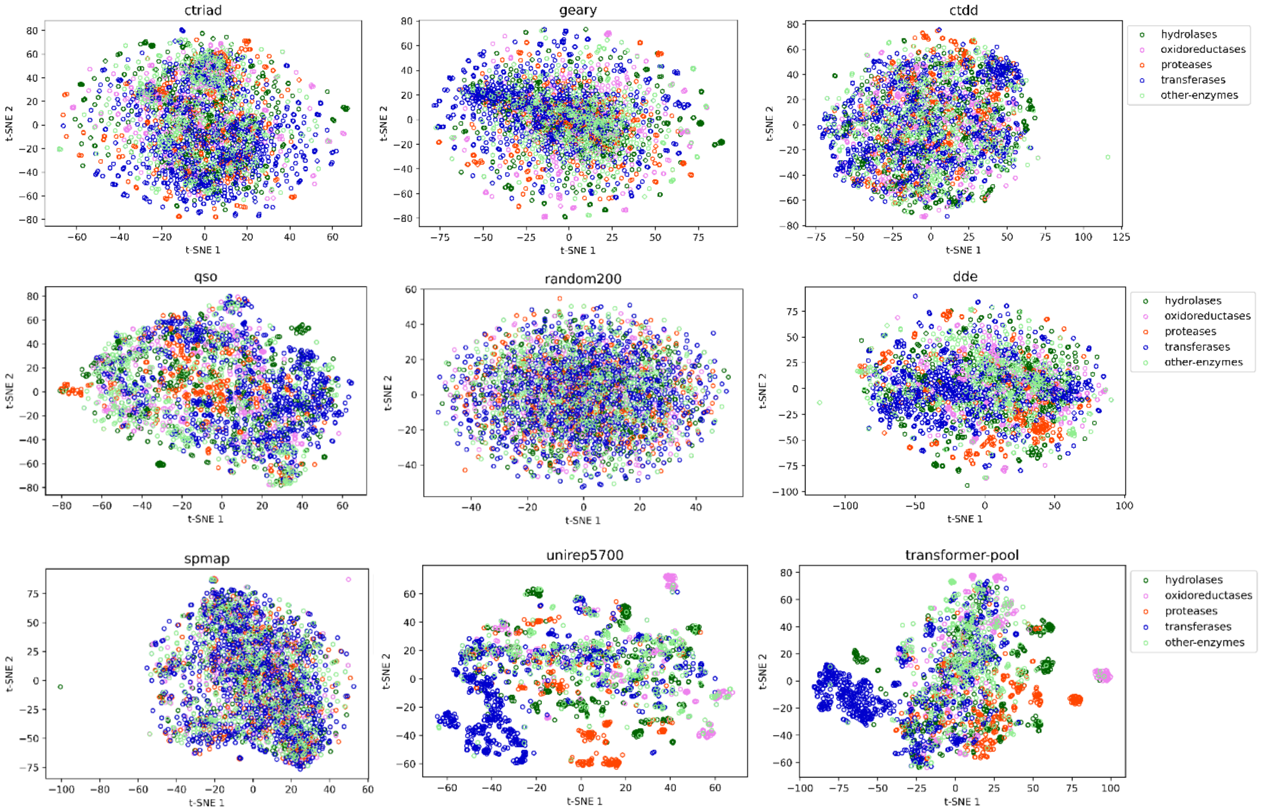


**(b)**

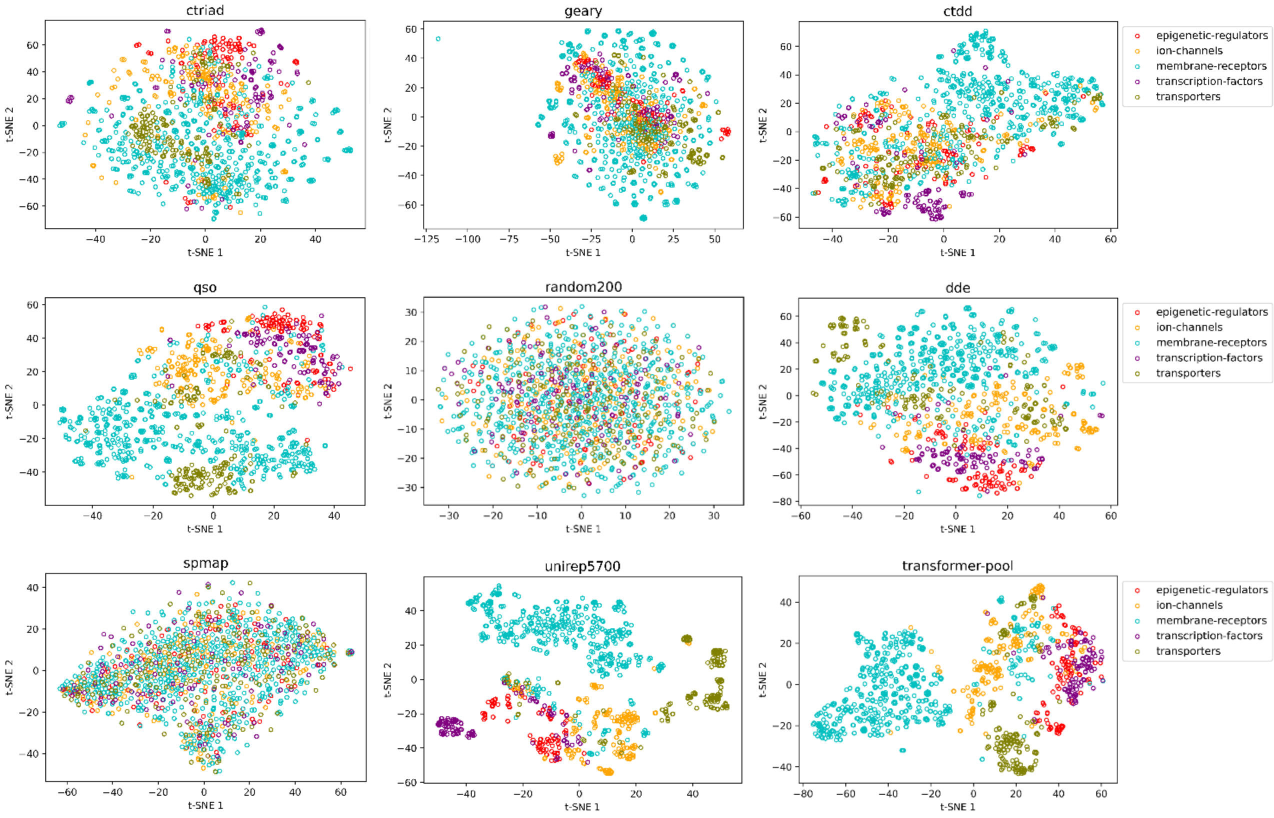


**Figure S1.** t-SNE based visualization of conventional and learned protein representations on; **(a)** enzymes including hydrolases, oxidoreductases, proteases, transferases, and other-enzymes, and **(b)** other protein families (non-enzymes) including epigenetic regulators, ion channels, membrane receptors, transcription factors and transporters.

(a)

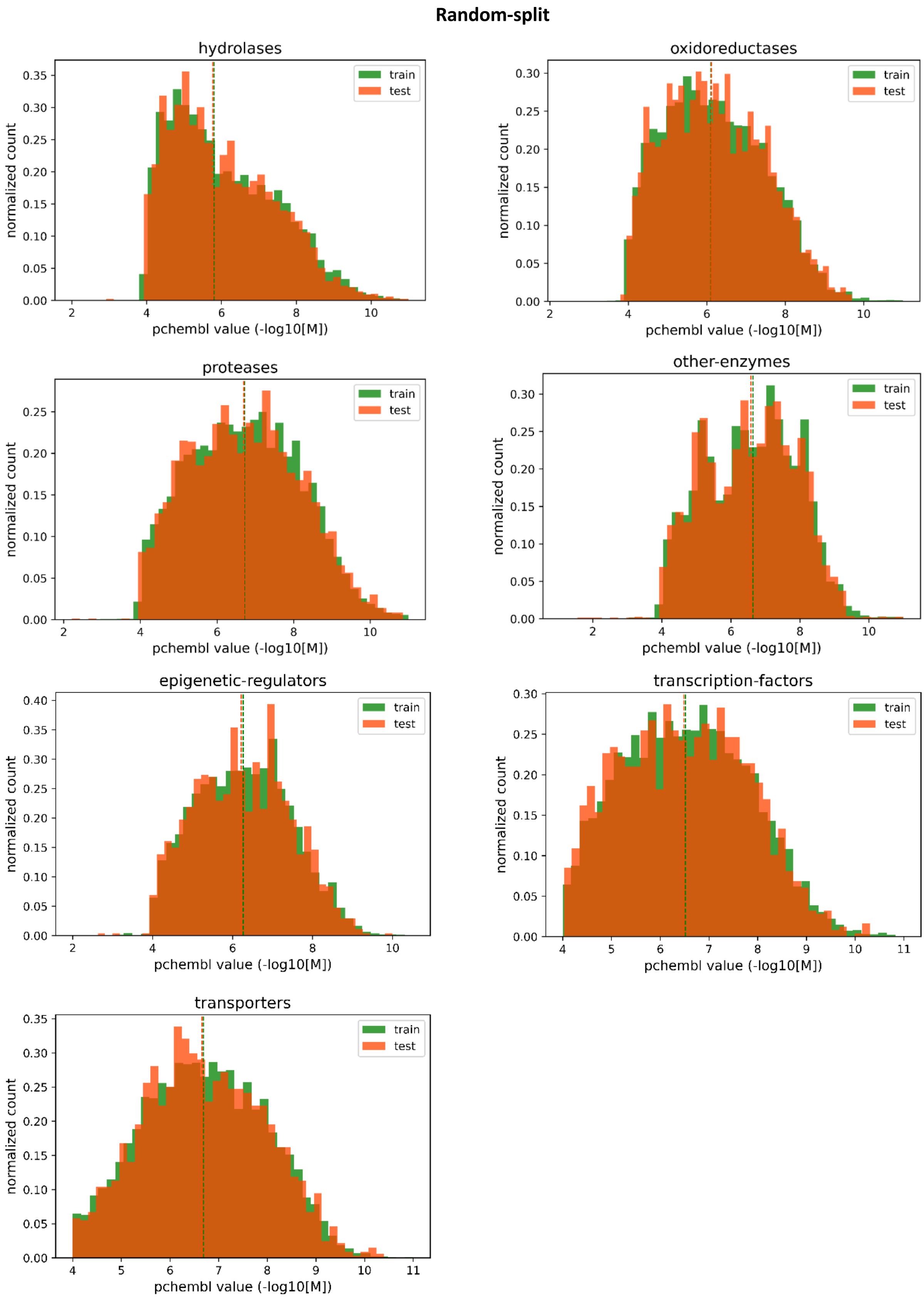
Random-split
hydrolases
oxidoreductases
proteases
other-enzymes
epigenetic-regulators
transcription-factors
transporters
train
test
normalized count
pchembl value (-log10[M])

**(b)**

**Dissimilar-compound-split**

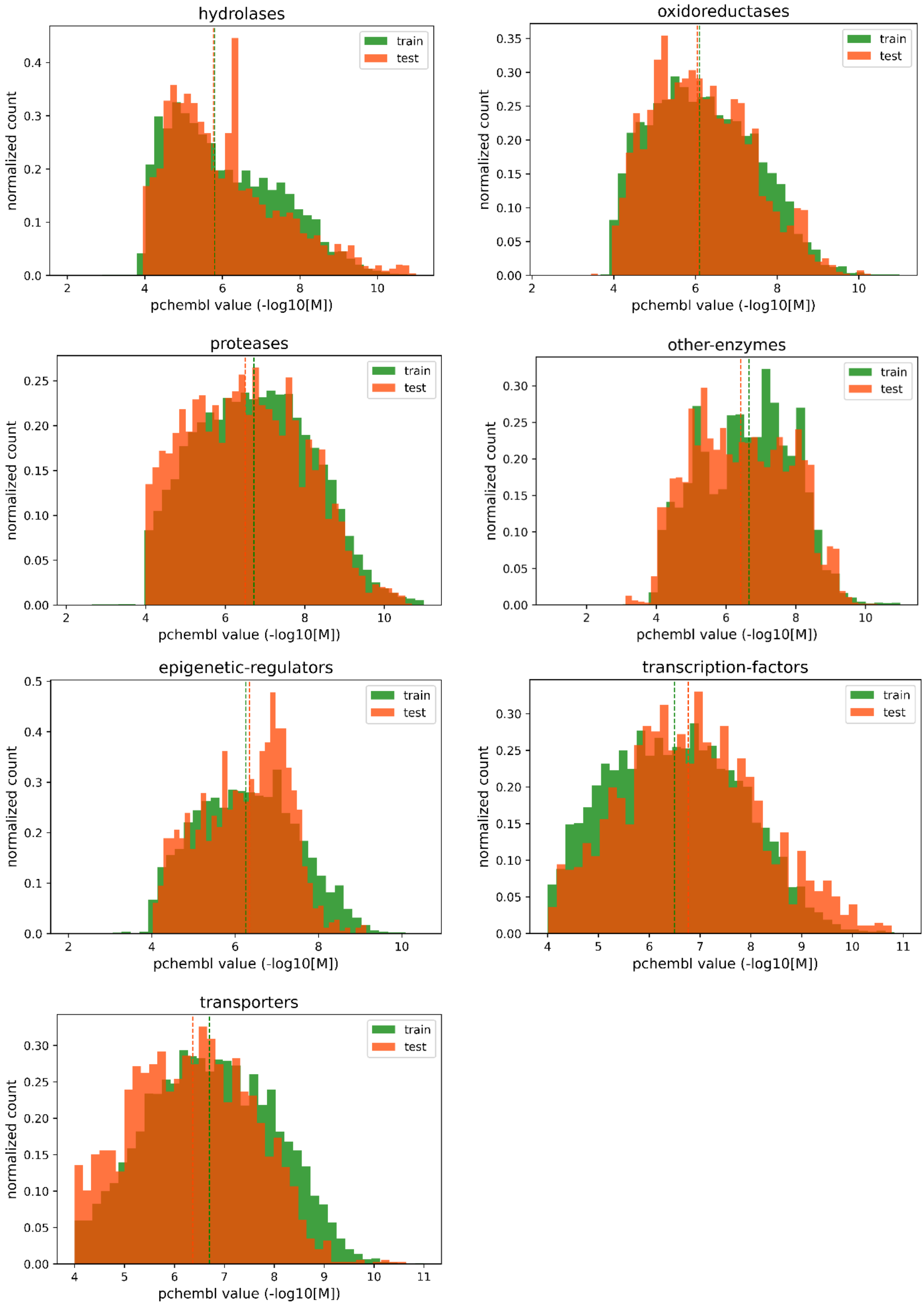

**(c)**

**Fully-dissimilar-split**

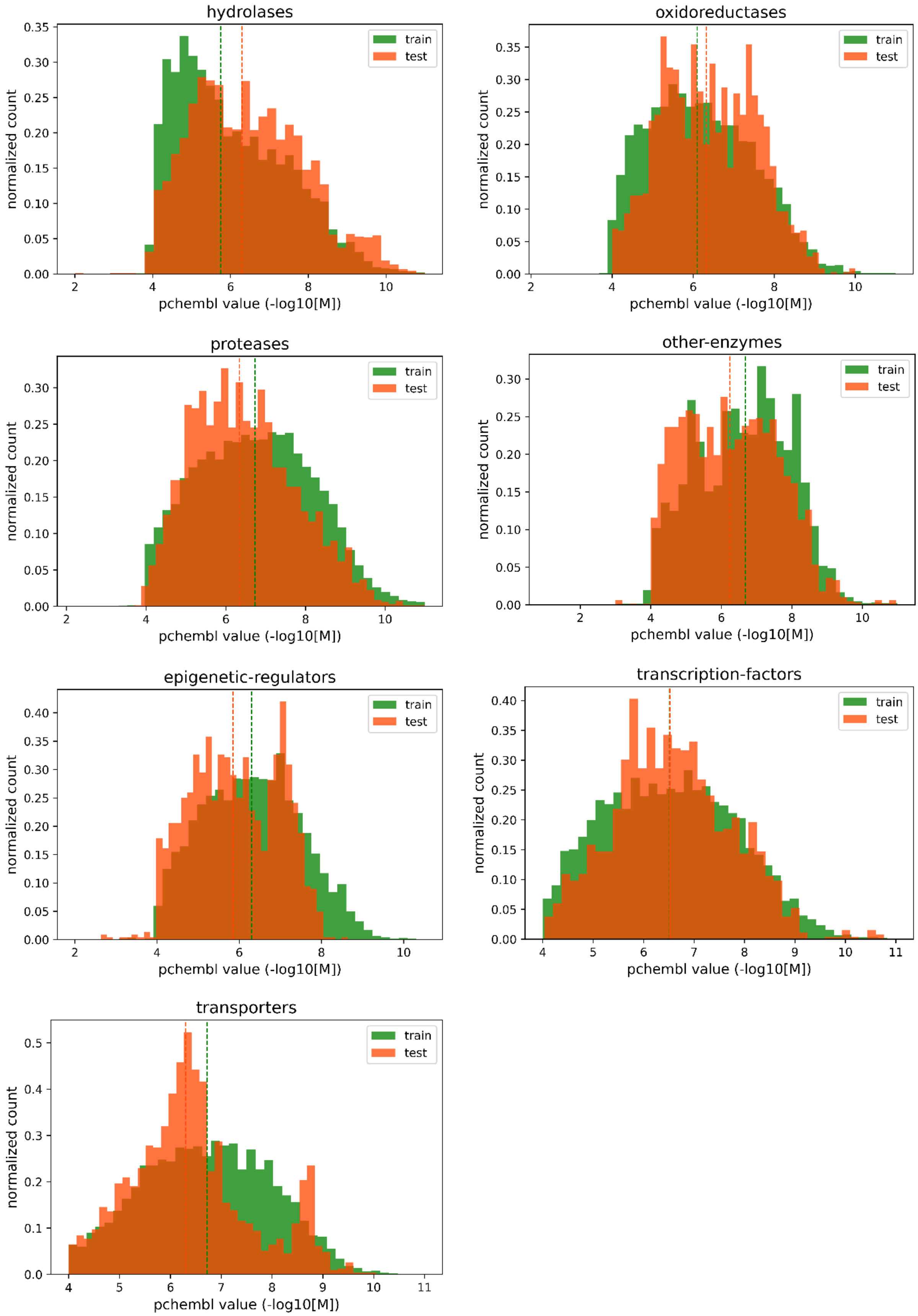


**Figure S2.** Bioactivity distributions of protein family-specific datasets in terms of; **(a)** random split, **(b)** dissimilar-compound split, and **(c)** fully-dissimilar split sets, together with the median values shown as vertical dashed lines.

**(a)**

### Random-split

hydrolases

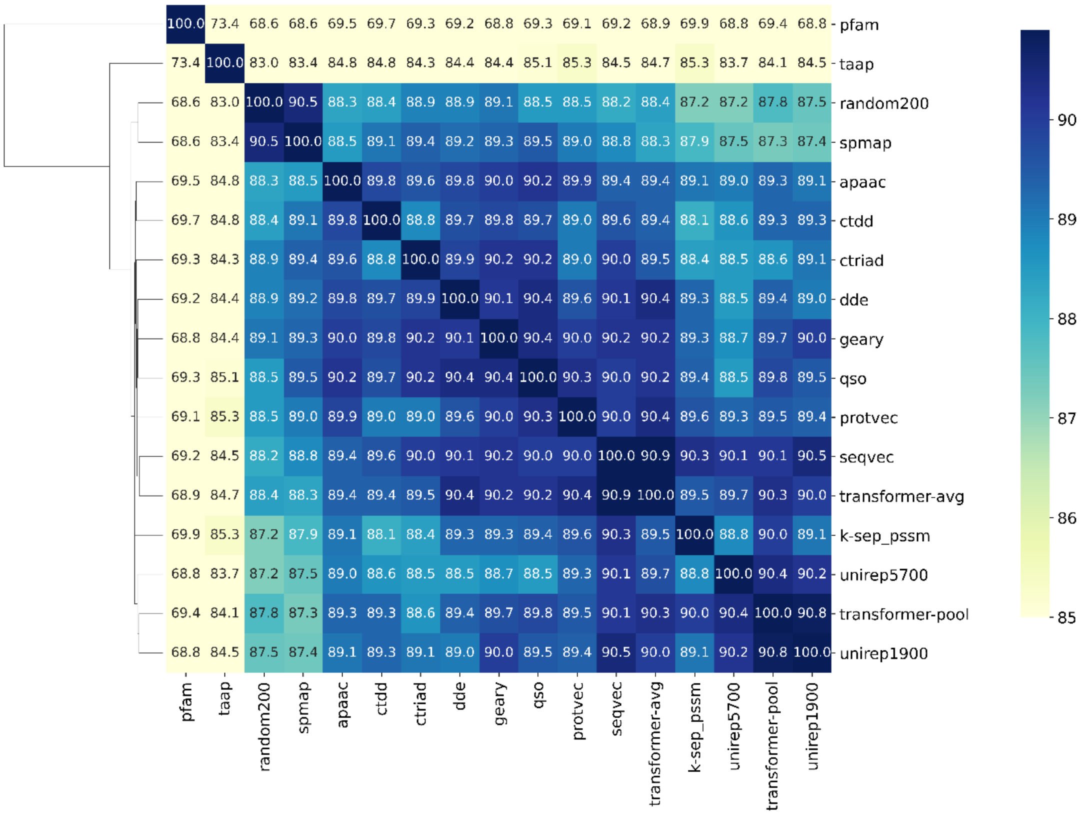


oxidoreductases

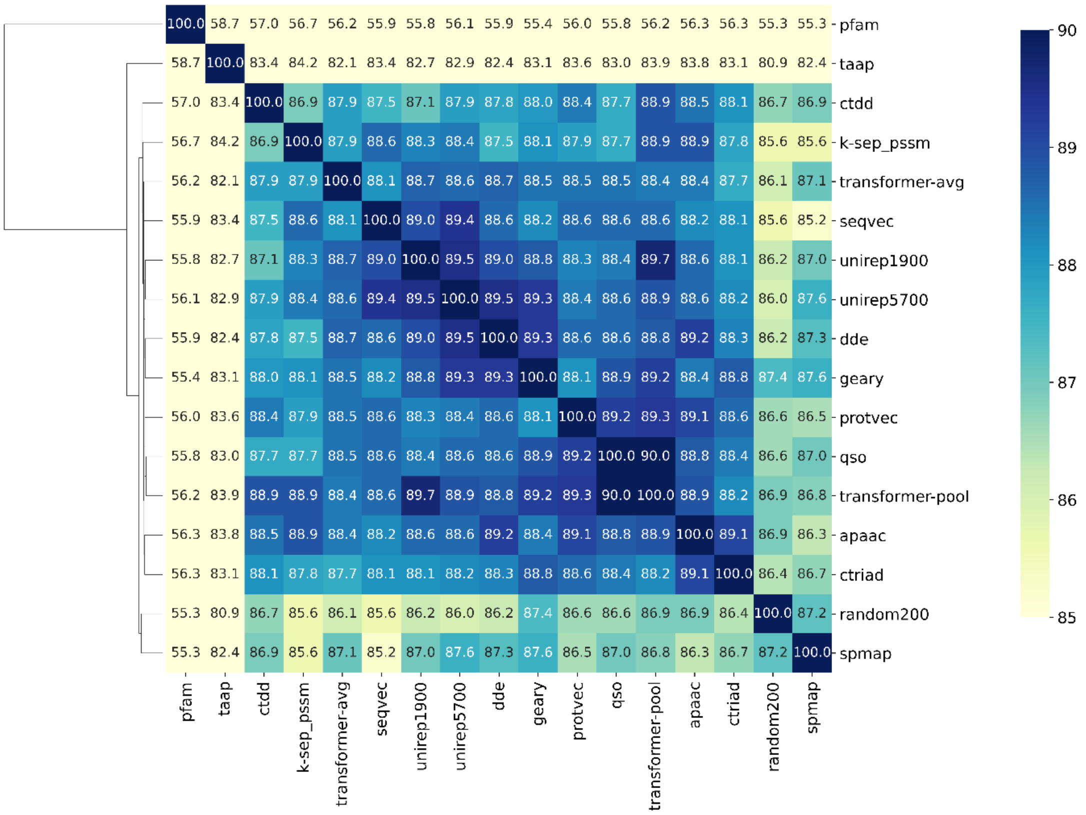

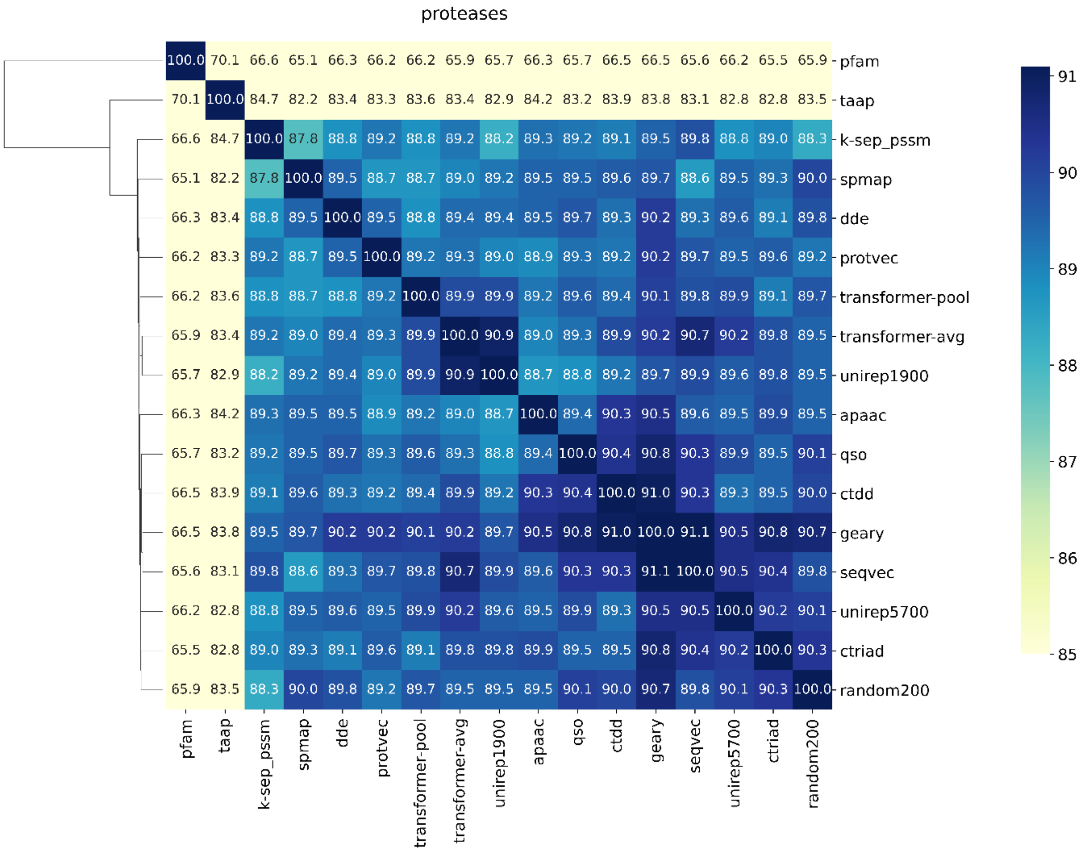
proteases
pfam 100.0 70.1 66.6 65.1 66.3 66.2 66.2 65.9 65.7 66.3 65.7 66.5 66.5 65.6 66.2 65.5 65.9
taap 70.1 100.0 84.7 82.2 83.4 83.3 83.6 83.4 82.9 84.2 83.2 83.9 83.8 83.1 82.8 82.8 83.5
k-sep_pssm 66.6 84.7 100.0 87.8 88.8 89.2 88.8 89.2 88.2 89.3 89.2 89.1 89.5 89.8 88.8 89.0 88.3
spmap 65.1 82.2 87.8 100.0 89.5 88.7 88.7 89.0 89.2 89.5 89.5 89.6 89.7 88.6 89.5 89.3 90.0
dde 66.3 83.4 88.8 89.5 100.0 89.5 88.8 89.4 89.4 89.5 89.7 89.3 90.2 89.3 89.6 89.1 89.8
protvec 66.2 83.3 89.2 88.7 89.5 100.0 89.2 89.3 89.0 88.9 89.3 89.2 90.2 89.7 89.5 89.6 89.2
transformer-pool 66.2 83.6 88.8 88.7 88.8 89.2 100.0 89.9 89.9 89.2 89.6 89.4 90.1 89.8 89.9 89.1 89.7
transformer-avg 65.9 83.4 89.2 89.0 89.4 89.3 89.9 100.0 90.9 89.0 89.3 89.9 90.2 90.7 90.2 89.8 89.5
unirep1900 65.7 82.9 88.2 89.2 89.4 89.0 89.9 90.9 100.0 88.7 88.8 89.2 89.7 89.9 89.6 89.8 89.5
apaac 66.3 84.2 89.3 89.5 89.5 88.9 89.2 89.0 88.7 100.0 89.4 90.3 90.5 89.6 89.5 89.9 89.5
qso 65.7 83.2 89.2 89.5 89.7 89.3 89.6 89.3 88.8 89.4 100.0 90.4 90.8 90.3 89.9 89.5 90.1
ctdd 66.5 83.9 89.1 89.6 89.3 89.2 89.4 89.9 89.2 90.3 90.4 100.0 91.0 90.3 89.3 89.5 90.0
geary 66.5 83.8 89.5 89.7 90.2 90.2 90.1 90.2 89.7 90.5 90.8 91.0 100.0 91.1 90.5 90.8 90.7
seqvec 65.6 83.1 89.8 88.6 89.3 89.7 89.8 90.7 89.9 89.6 90.3 90.3 91.1 100.0 90.5 90.4 89.8
unirep5700 66.2 82.8 88.8 89.5 89.6 89.5 89.9 90.2 89.6 89.5 89.9 89.3 90.5 90.5 100.0 90.2 90.1
ctriad 65.5 82.8 89.0 89.3 89.1 89.6 89.1 89.8 89.8 89.9 89.5 89.5 90.8 90.4 90.2 100.0 90.3
random200 65.9 83.5 88.3 90.0 89.8 89.2 89.7 89.5 89.5 89.5 90.1 90.0 90.7 89.8 90.1 90.3 100.0
pfam taap k-sep_pssm spmap dde protvec transformer-pool transformer-avg unirep1900 apaac qso ctdd geary seqvec unirep5700 ctriad random200
91 90 89 88 87 86 85

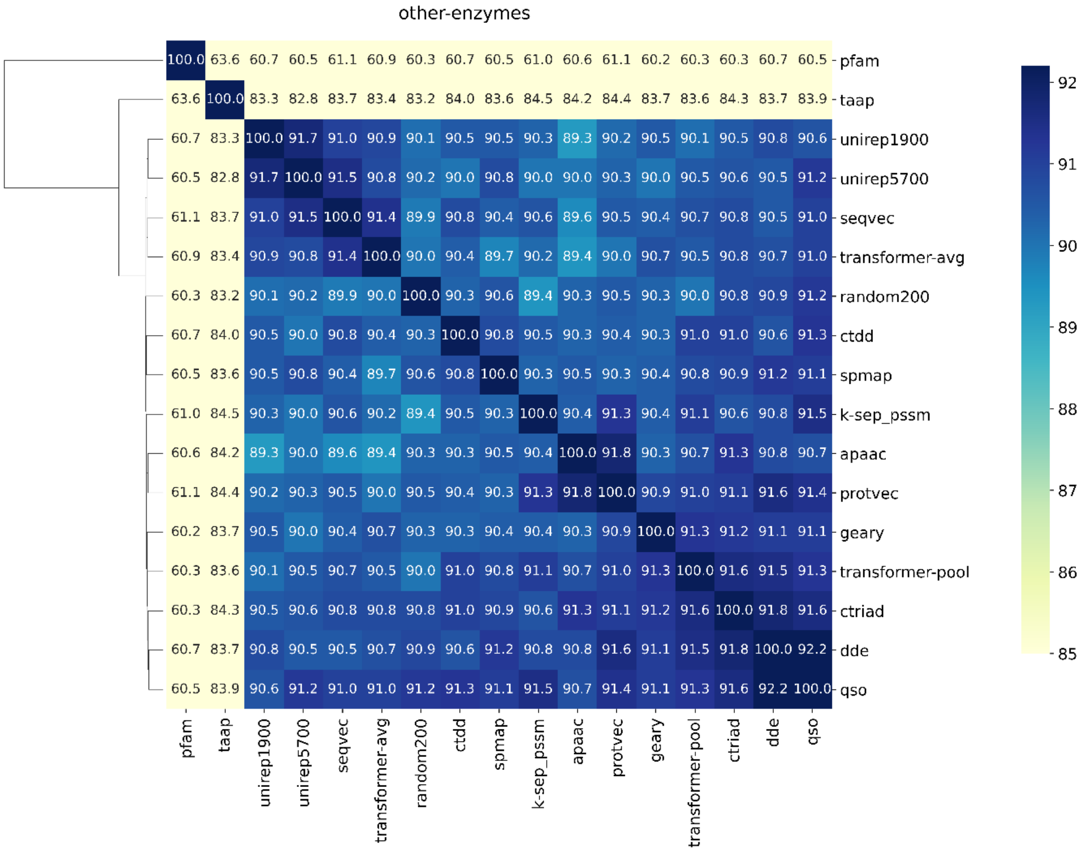
other-enzymes
pfam 100.0 63.6 60.7 60.5 61.1 60.9 60.3 60.7 60.5 61.0 60.6 61.1 60.2 60.3 60.3 60.7 60.5
taap 63.6 100.0 83.3 82.8 83.7 83.4 83.2 84.0 83.6 84.5 84.2 84.4 83.7 83.6 84.3 83.7 83.9
unirep1900 60.7 83.3 100.0 91.7 91.0 90.9 90.1 90.5 90.5 90.3 89.3 90.2 90.5 90.1 90.5 90.8 90.6
unirep5700 60.5 82.8 91.7 100.0 91.5 90.8 90.2 90.0 90.8 90.0 90.0 90.3 90.0 90.5 90.6 90.5 91.2
seqvec 61.1 83.7 91.0 91.5 100.0 91.4 89.9 90.8 90.4 90.6 89.6 90.5 90.4 90.7 90.8 90.5 91.0
transformer-avg 60.9 83.4 90.9 90.8 91.4 100.0 90.0 90.4 89.7 90.2 89.4 90.0 90.7 90.5 90.8 90.7 91.0
random200 60.3 83.2 90.1 90.2 89.9 90.0 100.0 90.3 90.6 89.4 90.3 90.5 90.3 90.0 90.8 90.9 91.2
ctdd 60.7 84.0 90.5 90.0 90.8 90.4 90.3 100.0 90.8 90.5 90.3 90.4 90.3 91.0 91.0 90.6 91.3
spmap 60.5 83.6 90.5 90.8 90.4 89.7 90.6 90.8 100.0 90.3 90.5 90.3 90.4 90.8 90.9 91.2 91.1
k-sep_pssm 61.0 84.5 90.3 90.0 90.6 90.2 89.4 90.5 90.3 100.0 90.4 91.3 90.4 91.1 90.6 90.8 91.5
apaac 60.6 84.2 89.3 90.0 89.6 89.4 90.3 90.3 90.5 90.4 100.0 91.8 90.3 90.7 91.3 90.8 90.7
protvec 61.1 84.4 90.2 90.3 90.5 90.0 90.5 90.4 90.3 91.3 91.8 100.0 90.9 91.0 91.1 91.6 91.4
geary 60.2 83.7 90.5 90.0 90.4 90.7 90.3 90.3 90.4 90.4 90.3 90.9 100.0 91.3 91.2 91.1 91.1
transformer-pool 60.3 83.6 90.1 90.5 90.7 90.5 90.0 91.0 90.8 91.1 90.7 91.0 91.3 100.0 91.6 91.5 91.3
ctriad 60.3 84.3 90.5 90.6 90.8 90.8 90.8 91.0 90.9 90.6 91.3 91.1 91.2 91.6 100.0 91.8 91.6
dde 60.7 83.7 90.8 90.5 90.5 90.7 90.9 90.6 91.2 90.8 90.8 91.6 91.1 91.5 91.8 100.0 92.2
qso 60.5 83.9 90.6 91.2 91.0 91.0 91.2 91.3 91.1 91.5 90.7 91.4 91.1 91.3 91.6 92.2 100.0
pfam taap unirep1900 unirep5700 seqvec transformer-avg random200 ctdd spmap k-sep_pssm apaac protvec geary transformer-pool ctriad dde qso
92 91 90 89 88 87 86 85

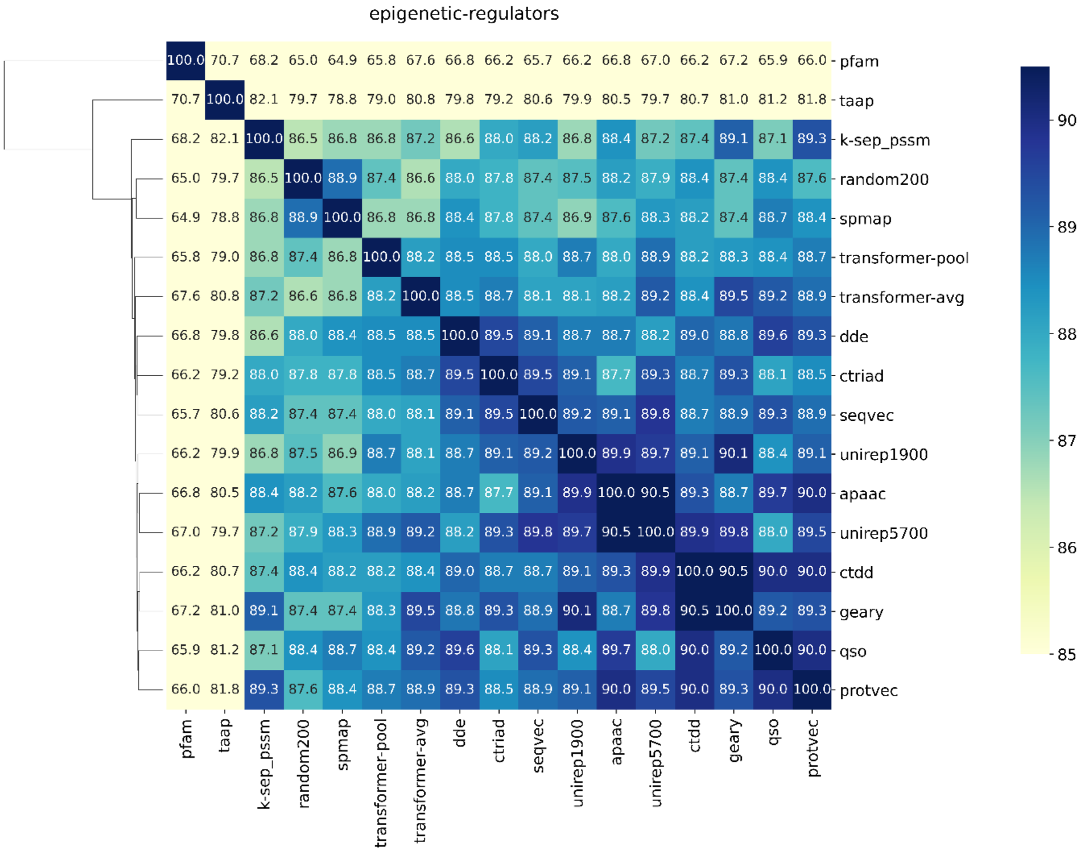
epigenetic-regulators
pfam
taap
k-sep_pssm
random200
spmap
transformer-pool
transformer-avg
dde
ctriad
seqvec
unirep1900
apaac
unirep5700
ctdd
geary
qso
protvec
90
89
88
87
86
85

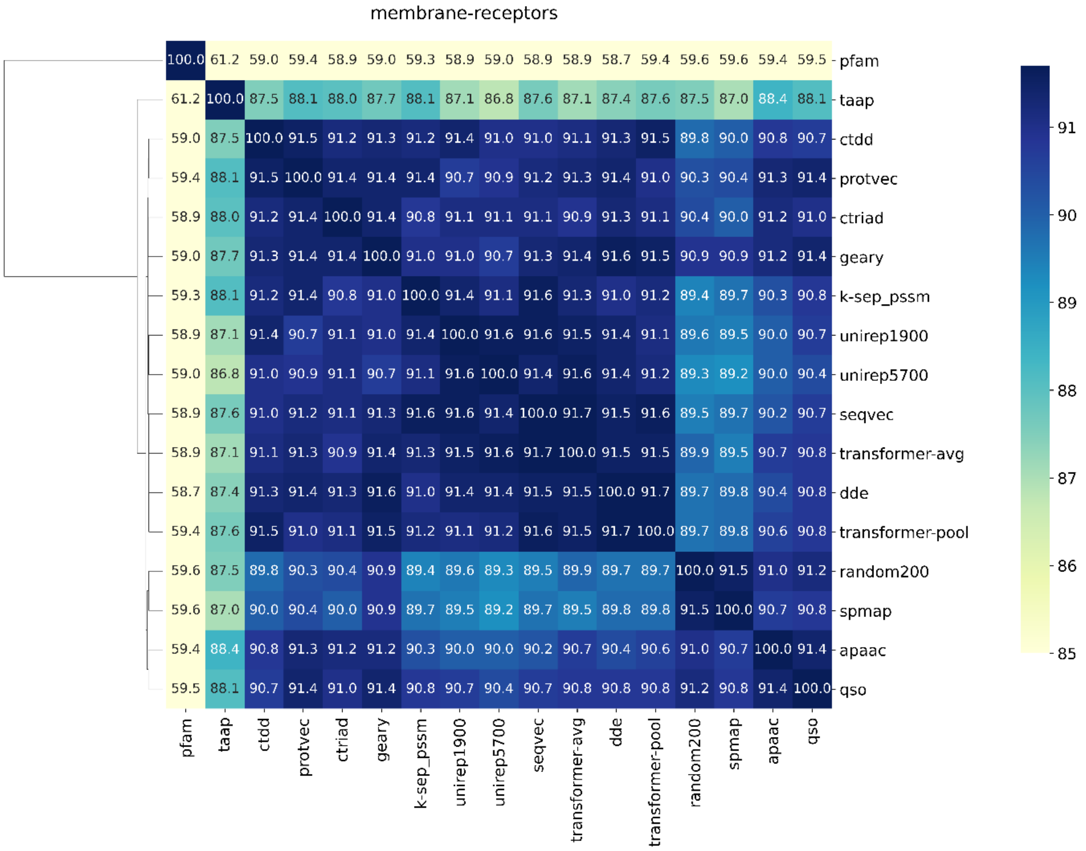
membrane-receptors
pfam
taap
ctdd
protvec
ctriad
geary
k-sep_pssm
unirep1900
unirep5700
seqvec
transformer-avg
dde
transformer-pool
random200
spmap
apaac
qso
91
90
89
88
87
86
85

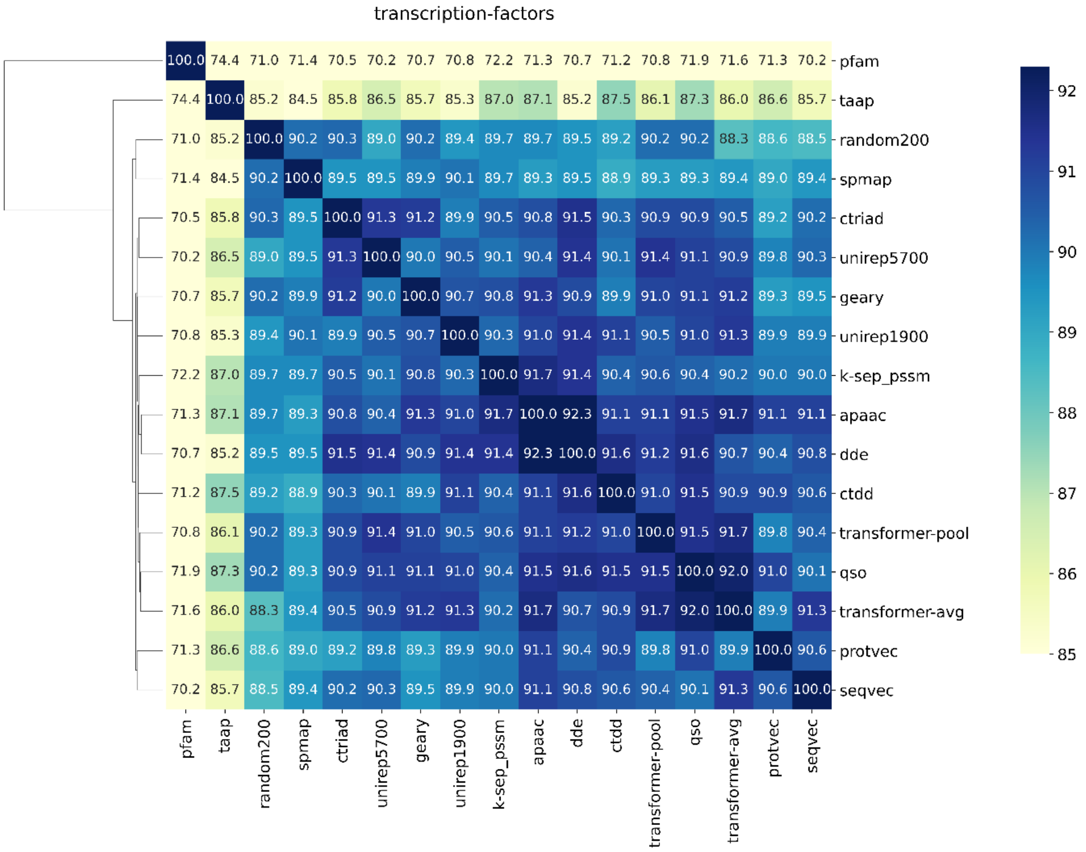
transcription-factors
pfam
taap
random200
spmap
ctriad
unirep5700
geary
unirep1900
k-sep_pssm
apaac
dde
ctdd
transformer-pool
qso
transformer-avg
protvec
seqvec
92
91
90
89
88
87
86
85

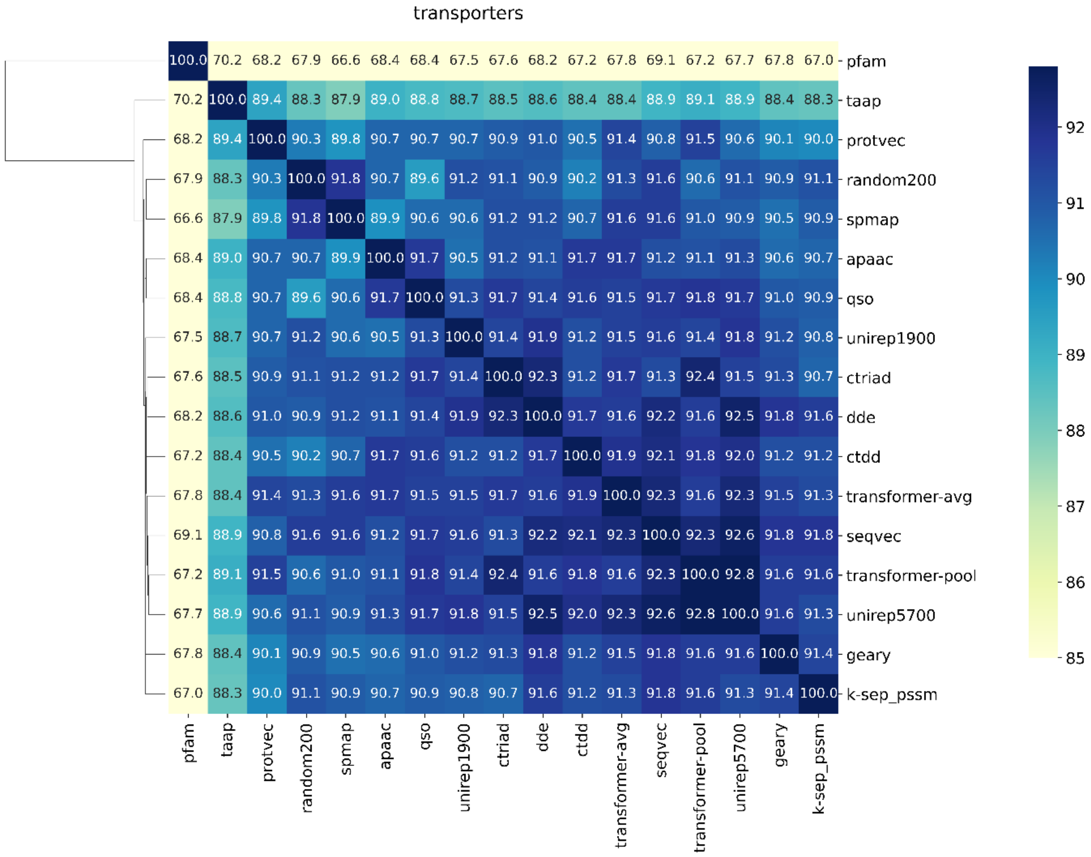
transporters
pfam
taap
protvec
random200
spmap
apaac
qso
unirep1900
ctriad
dde
ctdd
transformer-avg
seqvec
transformer-pool
unirep5700
geary
k-sep_pssm
92
91
90
89
88
87
86
85

**(b)**

### Dissimilar-compound-split

hydrolases

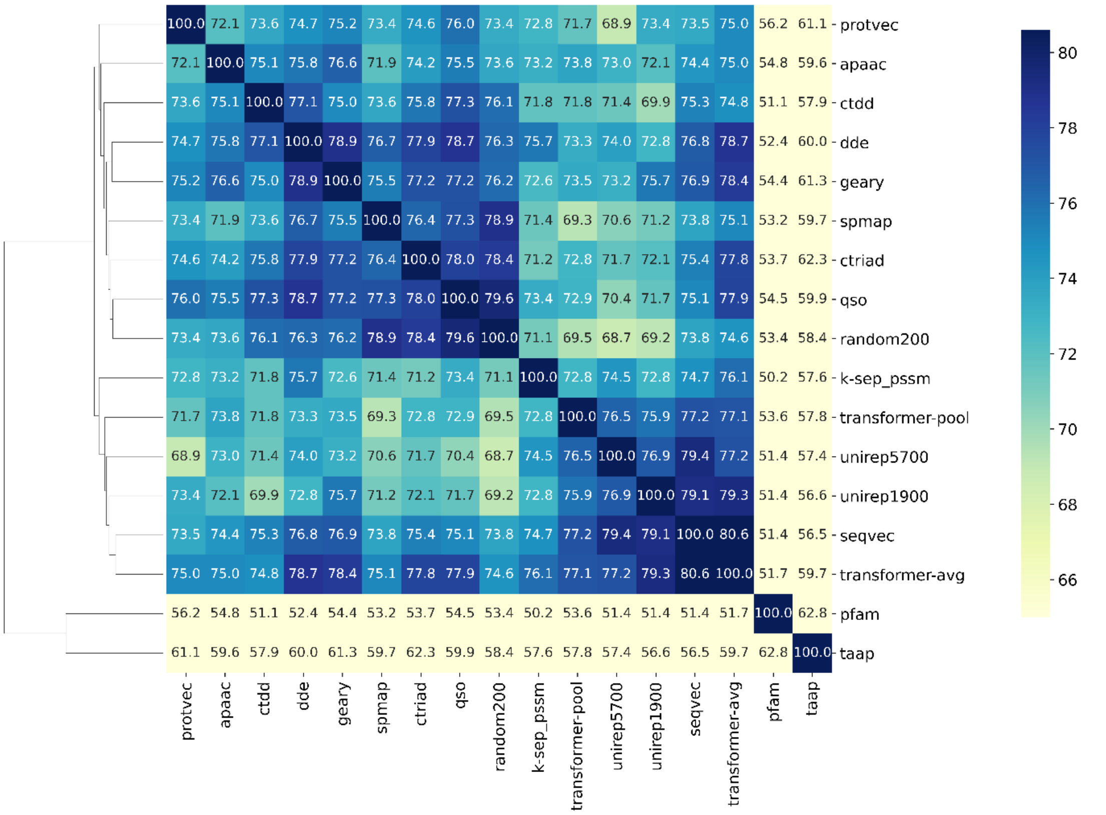


oxidoreductases

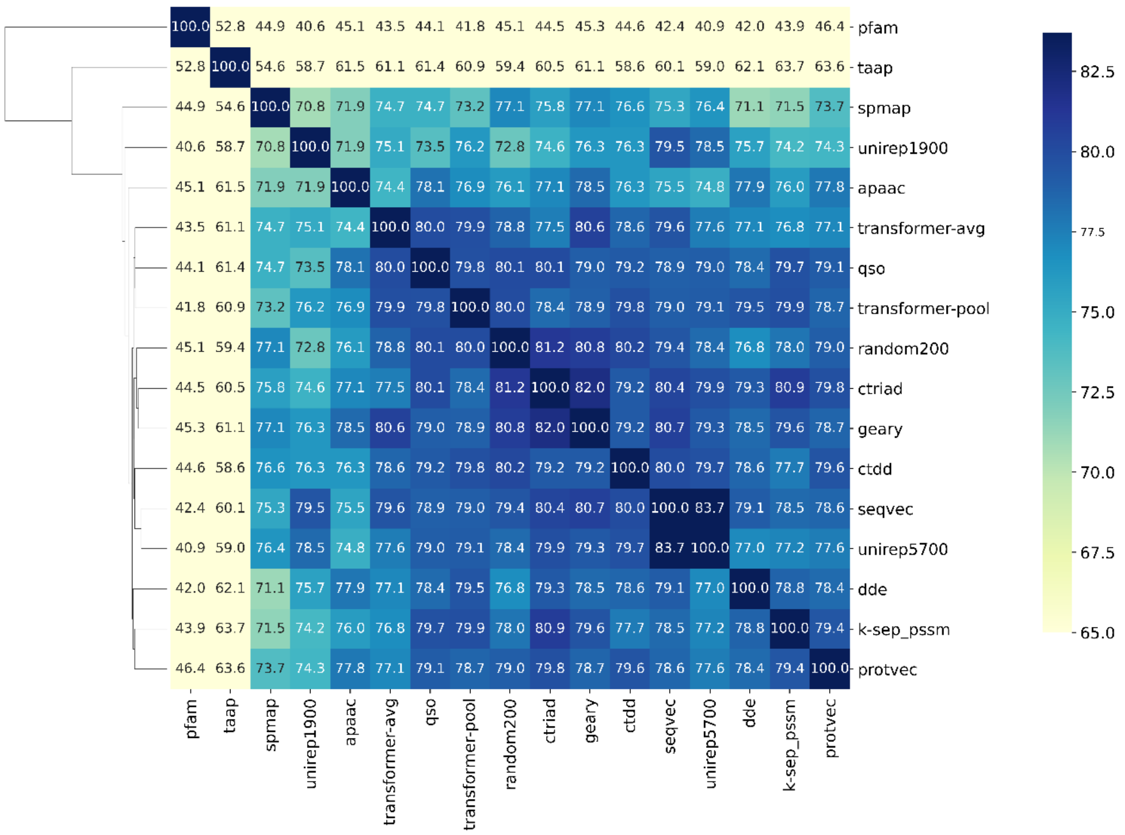

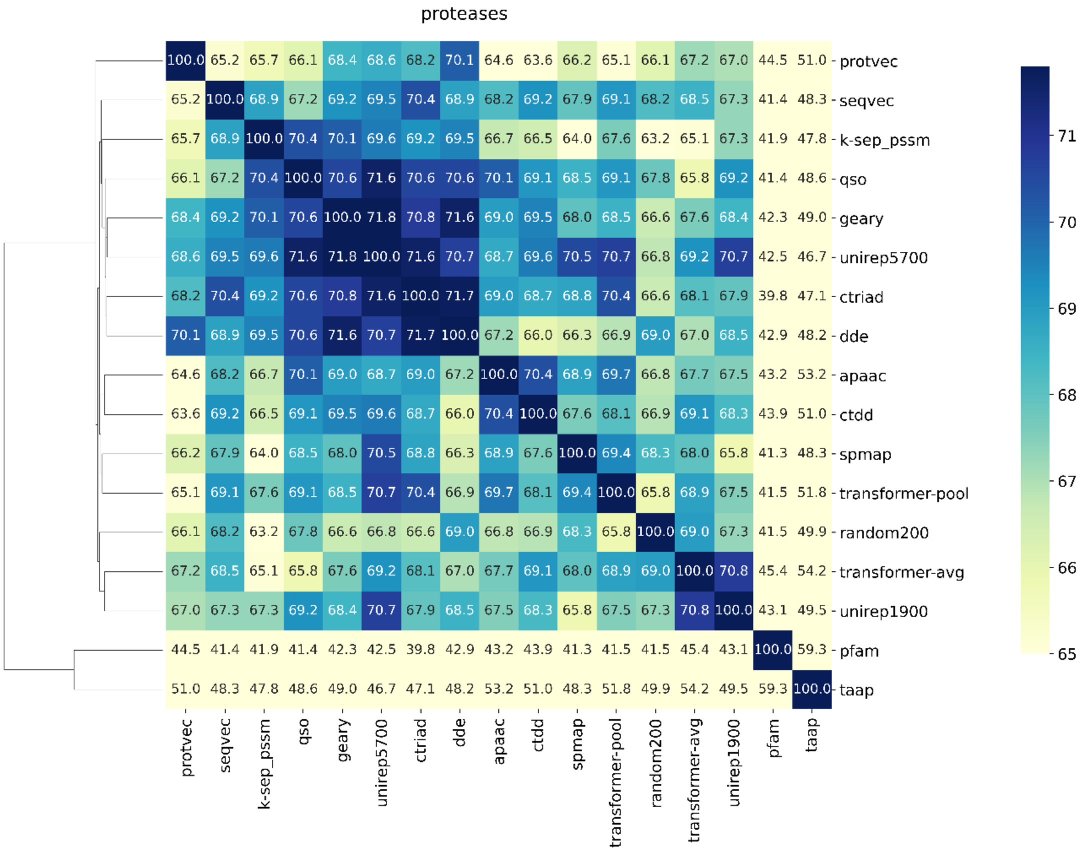
proteases
100.0 65.2 65.7 66.1 68.4 68.6 68.2 70.1 64.6 63.6 66.2 65.1 66.1 67.2 67.0 44.5 51.0 - protvec
65.2 100.0 68.9 67.2 69.2 69.5 70.4 68.9 68.2 69.2 67.9 69.1 68.2 68.5 67.3 41.4 48.3 - seqvec
65.7 68.9 100.0 70.4 70.1 69.6 69.2 69.5 66.7 66.5 64.0 67.6 63.2 65.1 67.3 41.9 47.8 - k-sep_pssm
66.1 67.2 70.4 100.0 70.6 71.6 70.6 70.6 70.1 69.1 68.5 69.1 67.8 65.8 69.2 41.4 48.6 - qso
68.4 69.2 70.1 70.6 100.0 71.8 70.8 71.6 69.0 69.5 68.0 68.5 66.6 67.6 68.4 42.3 49.0 - geary
68.6 69.5 69.6 71.6 71.8 100.0 71.6 70.7 68.7 69.6 70.5 70.7 66.8 69.2 70.7 42.5 46.7 - unirep5700
68.2 70.4 69.2 70.6 70.8 71.6 100.0 71.7 69.0 68.7 68.8 70.4 66.6 68.1 67.9 39.8 47.1 - ctriad
70.1 68.9 69.5 70.6 71.6 70.7 71.7 100.0 67.2 66.0 66.3 66.9 69.0 67.0 68.5 42.9 48.2 - dde
64.6 68.2 66.7 70.1 69.0 68.7 69.0 67.2 100.0 70.4 68.9 69.7 66.8 67.7 67.5 43.2 53.2 - apaac
63.6 69.2 66.5 69.1 69.5 69.6 68.7 66.0 70.4 100.0 67.6 68.1 66.9 69.1 68.3 43.9 51.0 - ctdd
66.2 67.9 64.0 68.5 68.0 70.5 68.8 66.3 68.9 67.6 100.0 69.4 68.3 68.0 65.8 41.3 48.3 - spmap
65.1 69.1 67.6 69.1 68.5 70.7 70.4 66.9 69.7 68.1 69.4 100.0 65.8 68.9 67.5 41.5 51.8 - transformer-pool
66.1 68.2 63.2 67.8 66.6 66.8 66.6 69.0 66.8 66.9 68.3 65.8 100.0 69.0 67.3 41.5 49.9 - random200
67.2 68.5 65.1 65.8 67.6 69.2 68.1 67.0 67.7 69.1 68.0 68.9 69.0 100.0 70.8 45.4 54.2 - transformer-avg
67.0 67.3 67.3 69.2 68.4 70.7 67.9 68.5 67.5 68.3 65.8 67.5 67.3 70.8 100.0 43.1 49.5 - unirep1900
44.5 41.4 41.9 41.4 42.3 42.5 39.8 42.9 43.2 43.9 41.3 41.5 41.5 45.4 43.1 100.0 59.3 - pfam
51.0 48.3 47.8 48.6 49.0 46.7 47.1 48.2 53.2 51.0 48.3 51.8 49.9 54.2 49.5 59.3 100.0 - taap
protvec
seqvec
k-sep_pssm
qso
geary
unirep5700
ctriad
dde
apaac
ctdd
spmap
transformer-pool
random200
transformer-avg
unirep1900
pfam
taap
71
70
69
68
67
66
65

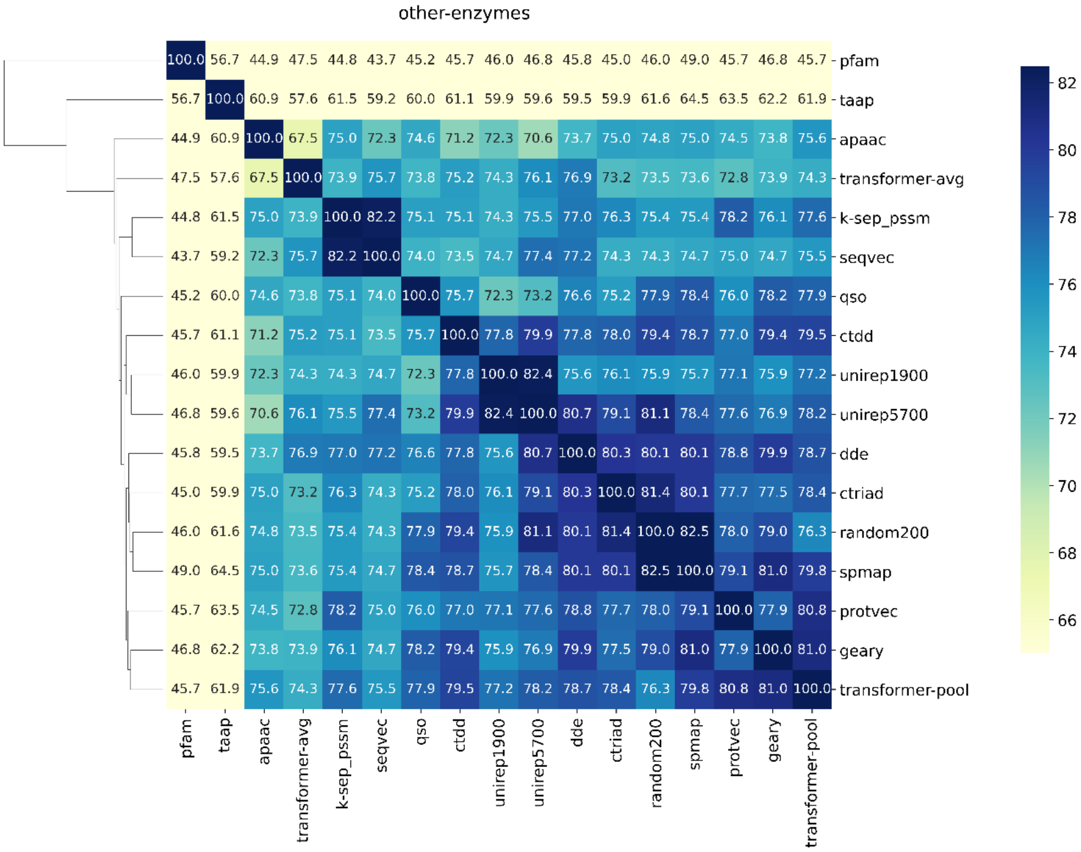
other-enzymes
100.0 56.7 44.9 47.5 44.8 43.7 45.2 45.7 46.0 46.8 45.8 45.0 46.0 49.0 45.7 46.8 45.7 - pfam
56.7 100.0 60.9 57.6 61.5 59.2 60.0 61.1 59.9 59.6 59.5 59.9 61.6 64.5 63.5 62.2 61.9 - taap
44.9 60.9 100.0 67.5 75.0 72.3 74.6 71.2 72.3 70.6 73.7 75.0 74.8 75.0 74.5 73.8 75.6 - apaac
47.5 57.6 67.5 100.0 73.9 75.7 73.8 75.2 74.3 76.1 76.9 73.2 73.5 73.6 72.8 73.9 74.3 - transformer-avg
44.8 61.5 75.0 73.9 100.0 82.2 75.1 75.1 74.3 75.5 77.0 76.3 75.4 75.4 78.2 76.1 77.6 - k-sep_pssm
43.7 59.2 72.3 75.7 82.2 100.0 74.0 73.5 74.7 77.4 77.2 74.3 74.3 74.7 75.0 74.7 75.5 - seqvec
45.2 60.0 74.6 73.8 75.1 74.0 100.0 75.7 72.3 73.2 76.6 75.2 77.9 78.4 76.0 78.2 77.9 - qso
45.7 61.1 71.2 75.2 75.1 73.5 75.7 100.0 77.8 79.9 77.8 78.0 79.4 78.7 77.0 79.4 79.5 - ctdd
46.0 59.9 72.3 74.3 74.3 74.7 72.3 77.8 100.0 82.4 75.6 76.1 75.9 75.7 77.1 75.9 77.2 - unirep1900
46.8 59.6 70.6 76.1 75.5 77.4 73.2 79.9 82.4 100.0 80.7 79.1 81.1 78.4 77.6 76.9 78.2 - unirep5700
45.8 59.5 73.7 76.9 77.0 77.2 76.6 77.8 75.6 80.7 100.0 80.3 80.1 80.1 78.8 79.9 78.7 - dde
45.0 59.9 75.0 73.2 76.3 74.3 75.2 78.0 76.1 79.1 80.3 100.0 81.4 80.1 77.7 77.5 78.4 - ctriad
46.0 61.6 74.8 73.5 75.4 74.3 77.9 79.4 75.9 81.1 80.1 81.4 100.0 82.5 78.0 79.0 76.3 - random200
49.0 64.5 75.0 73.6 75.4 74.7 78.4 78.7 75.7 78.4 80.1 80.1 82.5 100.0 79.1 81.0 79.8 - spmap
45.7 63.5 74.5 72.8 78.2 75.0 76.0 77.0 77.1 77.6 78.8 77.7 78.0 79.1 100.0 77.9 80.8 - protvec
46.8 62.2 73.8 73.9 76.1 74.7 78.2 79.4 75.9 76.9 79.9 77.5 79.0 81.0 77.9 100.0 81.0 - geary
45.7 61.9 75.6 74.3 77.6 75.5 77.9 79.5 77.2 78.2 78.7 78.4 76.3 79.8 80.8 81.0 100.0 - transformer-pool
pfam
taap
apaac
transformer-avg
k-sep_pssm
seqvec
qso
ctdd
unirep1900
unirep5700
dde
ctriad
random200
spmap
protvec
geary
transformer-pool
82
80
78
76
74
72
70
68
66

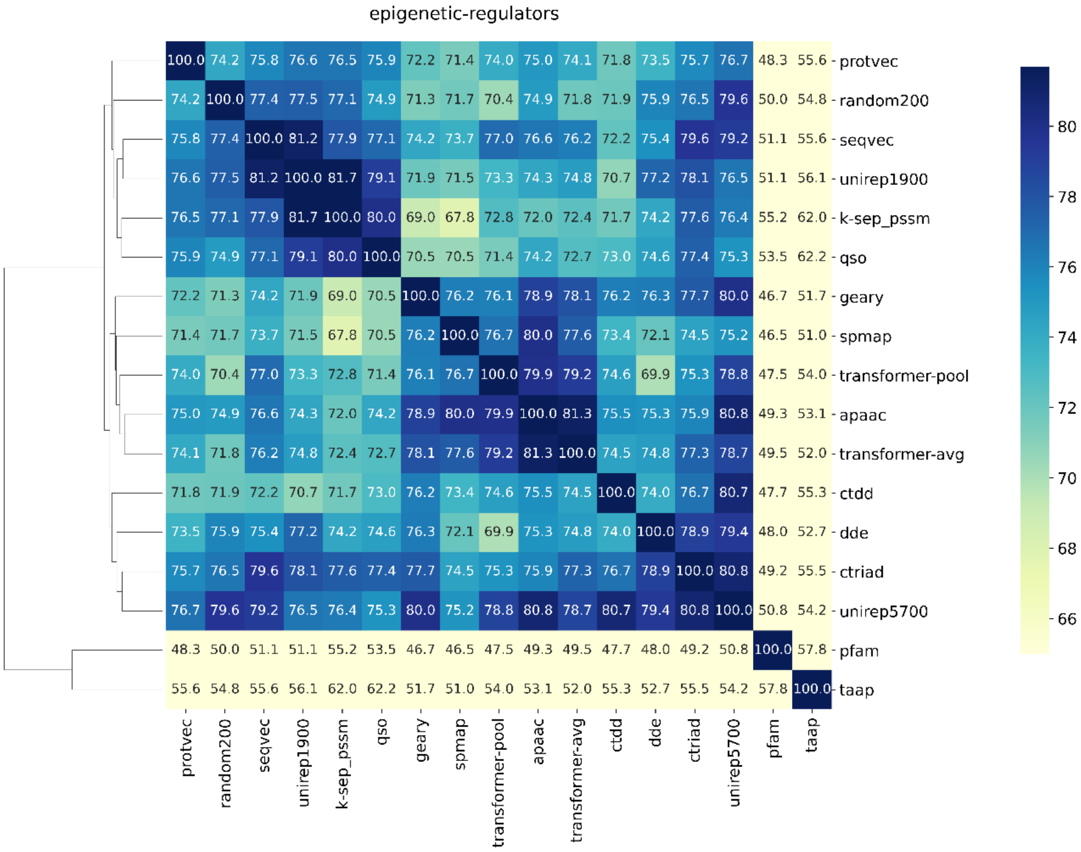
epigenetic-regulators
protvec: 100.0 74.2 75.8 76.6 76.5 75.9 72.2 71.4 74.0 75.0 74.1 71.8 73.5 75.7 76.7 48.3 55.6
random200: 74.2 100.0 77.4 77.5 77.1 74.9 71.3 71.7 70.4 74.9 71.8 71.9 75.9 76.5 79.6 50.0 54.8
seqvec: 75.8 77.4 100.0 81.2 77.9 77.1 74.2 73.7 77.0 76.6 76.2 72.2 75.4 79.6 79.2 51.1 55.6
unirep1900: 76.6 77.5 81.2 100.0 81.7 79.1 71.9 71.5 73.3 74.3 74.8 70.7 77.2 78.1 76.5 51.1 56.1
k-sep_pssm: 76.5 77.1 77.9 81.7 100.0 80.0 69.0 67.8 72.8 72.0 72.4 71.7 74.2 77.6 76.4 55.2 62.0
qso: 75.9 74.9 77.1 79.1 80.0 100.0 70.5 70.5 71.4 74.2 72.7 73.0 74.6 77.4 75.3 53.5 62.2
geary: 72.2 71.3 74.2 71.9 69.0 70.5 100.0 76.2 76.1 78.9 78.1 76.2 76.3 77.7 80.0 46.7 51.7
spmap: 71.4 71.7 73.7 71.5 67.8 70.5 76.2 100.0 76.7 80.0 77.6 73.4 72.1 74.5 75.2 46.5 51.0
transformer-pool: 74.0 70.4 77.0 73.3 72.8 71.4 76.1 76.7 100.0 79.9 79.2 74.6 69.9 75.3 78.8 47.5 54.0
apaac: 75.0 74.9 76.6 74.3 72.0 74.2 78.9 80.0 79.9 100.0 81.3 75.5 75.3 75.9 80.8 49.3 53.1
transformer-avg: 74.1 71.8 76.2 74.8 72.4 72.7 78.1 77.6 79.2 81.3 100.0 74.5 74.8 77.3 78.7 49.5 52.0
ctdd: 71.8 71.9 72.2 70.7 71.7 73.0 76.2 73.4 74.6 75.5 74.5 100.0 74.0 76.7 80.7 47.7 55.3
dde: 73.5 75.9 75.4 77.2 74.2 74.6 76.3 72.1 69.9 75.3 74.8 74.0 100.0 78.9 79.4 48.0 52.7
ctriad: 75.7 76.5 79.6 78.1 77.6 77.4 77.7 74.5 75.3 75.9 77.3 76.7 78.9 100.0 80.8 49.2 55.5
unirep5700: 76.7 79.6 79.2 76.5 76.4 75.3 80.0 75.2 78.8 80.8 78.7 80.7 79.4 80.8 100.0 50.8 54.2
pfam: 48.3 50.0 51.1 51.1 55.2 53.5 46.7 46.5 47.5 49.3 49.5 47.7 48.0 49.2 50.8 100.0 57.8
taap: 55.6 54.8 55.6 56.1 62.0 62.2 51.7 51.0 54.0 53.1 52.0 55.3 52.7 55.5 54.2 57.8 100.0
protvec random200 seqvec unirep1900 k-sep_pssm qso geary spmap transformer-pool apaac transformer-avg ctdd dde ctriad unirep5700 pfam taap
80 78 76 74 72 70 68 66

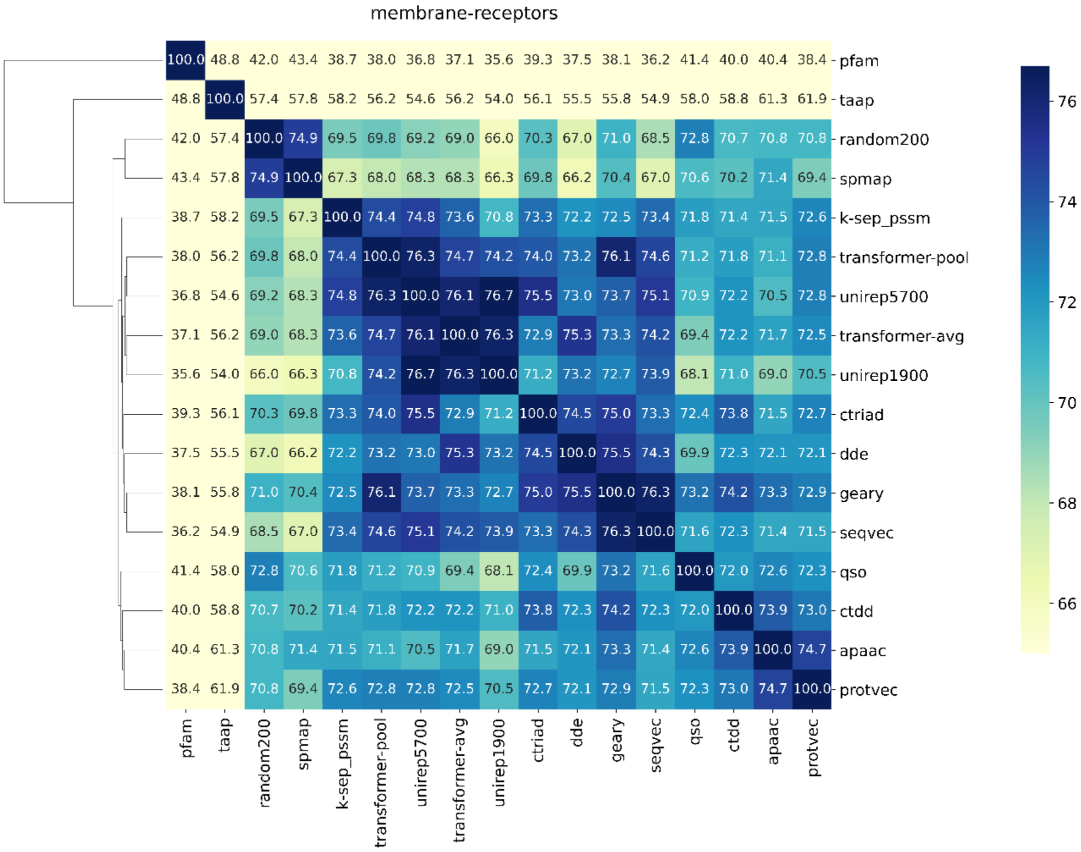
membrane-receptors
pfam: 100.0 48.8 42.0 43.4 38.7 38.0 36.8 37.1 35.6 39.3 37.5 38.1 36.2 41.4 40.0 40.4 38.4
taap: 48.8 100.0 57.4 57.8 58.2 56.2 54.6 56.2 54.0 56.1 55.5 55.8 54.9 58.0 58.8 61.3 61.9
random200: 42.0 57.4 100.0 74.9 69.5 69.8 69.2 69.0 66.0 70.3 67.0 71.0 68.5 72.8 70.7 70.8 70.8
spmap: 43.4 57.8 74.9 100.0 67.3 68.0 68.3 68.3 66.3 69.8 66.2 70.4 67.0 70.6 70.2 71.4 69.4
k-sep_pssm: 38.7 58.2 69.5 67.3 100.0 74.4 74.8 73.6 70.8 73.3 72.2 72.5 73.4 71.8 71.4 71.5 72.6
transformer-pool: 38.0 56.2 69.8 68.0 74.4 100.0 76.3 74.7 74.2 74.0 73.2 76.1 74.6 71.2 71.8 71.1 72.8
unirep5700: 36.8 54.6 69.2 68.3 74.8 76.3 100.0 76.1 76.7 75.5 73.0 73.7 75.1 70.9 72.2 70.5 72.8
transformer-avg: 37.1 56.2 69.0 68.3 73.6 74.7 76.1 100.0 76.3 72.9 75.3 73.3 74.2 69.4 72.2 71.7 72.5
unirep1900: 35.6 54.0 66.0 66.3 70.8 74.2 76.7 76.3 100.0 71.2 73.2 72.7 73.9 68.1 71.0 69.0 70.5
ctriad: 39.3 56.1 70.3 69.8 73.3 74.0 75.5 72.9 71.2 100.0 74.5 75.0 73.3 72.4 73.8 71.5 72.7
dde: 37.5 55.5 67.0 66.2 72.2 73.2 73.0 75.3 73.2 74.5 100.0 75.5 74.3 69.9 72.3 72.1 72.1
geary: 38.1 55.8 71.0 70.4 72.5 76.1 73.7 73.3 72.7 75.0 75.5 100.0 76.3 73.2 74.2 73.3 72.9
seqvec: 36.2 54.9 68.5 67.0 73.4 74.6 75.1 74.2 73.9 73.3 74.3 76.3 100.0 71.6 72.3 71.4 71.5
qso: 41.4 58.0 72.8 70.6 71.8 71.2 70.9 69.4 68.1 72.4 69.9 73.2 71.6 100.0 72.0 72.6 72.3
ctdd: 40.0 58.8 70.7 70.2 71.4 71.8 72.2 72.2 71.0 73.8 72.3 74.2 72.3 72.0 100.0 73.9 73.0
apaac: 40.4 61.3 70.8 71.4 71.5 71.1 70.5 71.7 69.0 71.5 72.1 73.3 71.4 72.6 73.9 100.0 74.7
protvec: 38.4 61.9 70.8 69.4 72.6 72.8 72.8 72.5 70.5 72.7 72.1 72.9 71.5 72.3 73.0 74.7 100.0
pfam taap random200 spmap k-sep_pssm transformer-pool unirep5700 transformer-avg unirep1900 ctriad dde geary seqvec qso ctdd apaac protvec
76 74 72 70 68 66

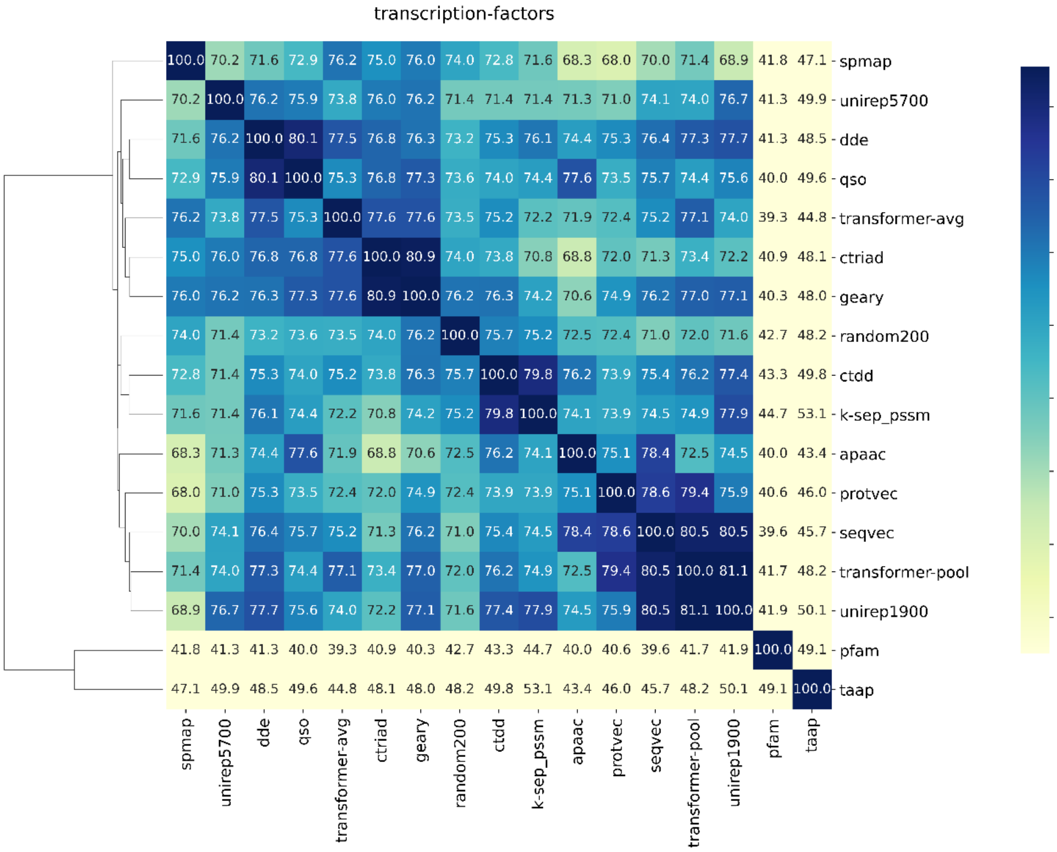
transcription-factors
spmap
unirep5700
dde
qso
transformer-avg
ctriad
geary
random200
ctdd
k-sep_pssm
apaac
protvec
seqvec
transformer-pool
unirep1900
pfam
taap
80
78
76
74
72
70
68
66

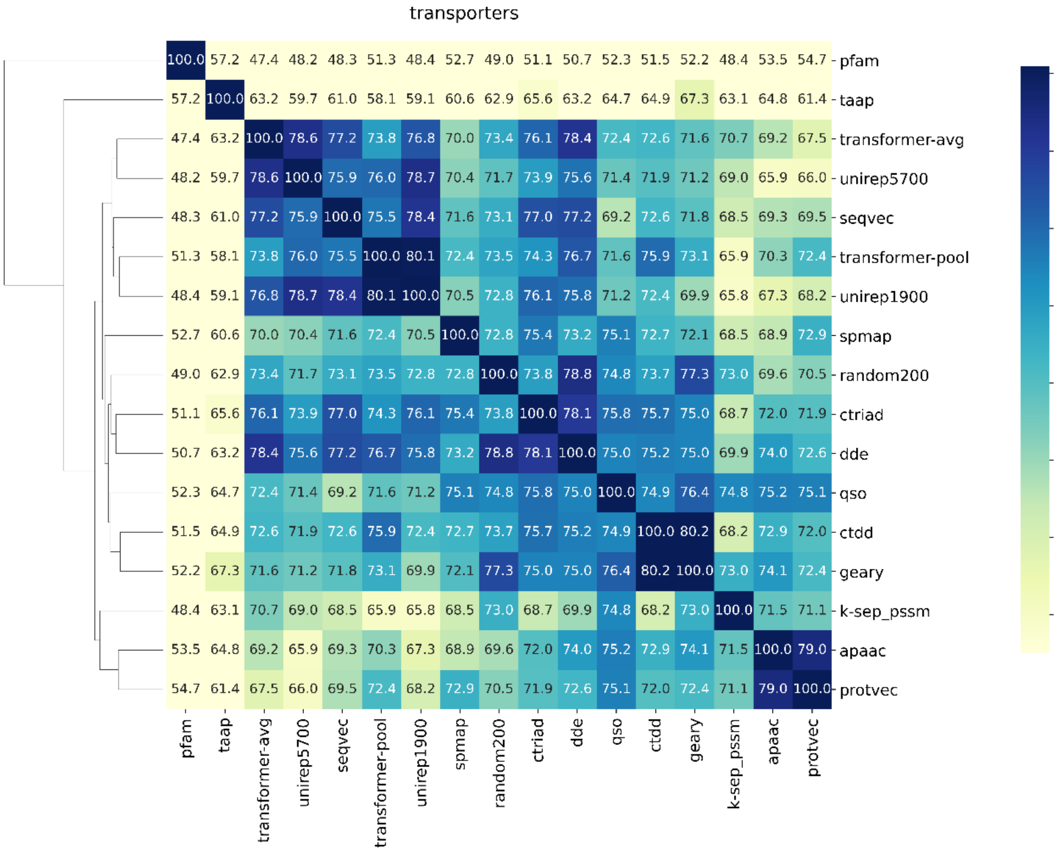
transporters
pfam
taap
transformer-avg
unirep5700
seqvec
transformer-pool
unirep1900
spmap
random200
ctriad
dde
qso
ctdd
geary
k-sep_pssm
apaac
protvec
80
78
76
74
72
70
68
66

**(c)**

### Fully-dissimilar-split

hydrolases

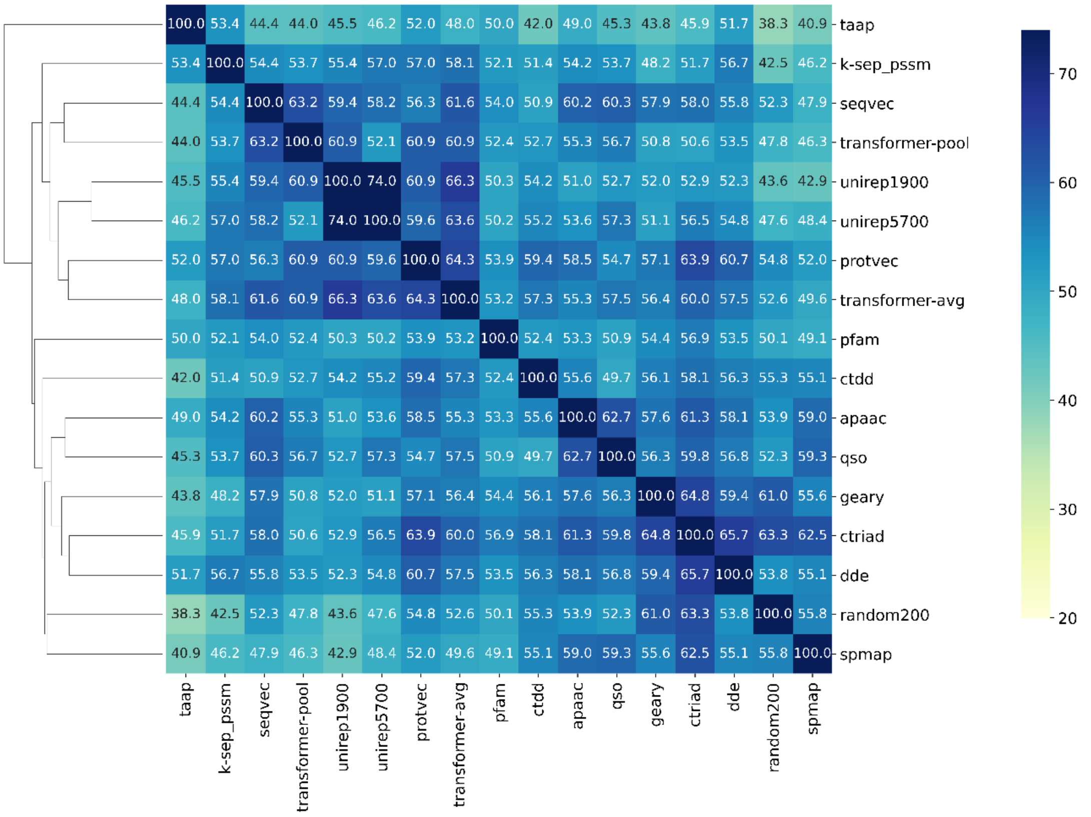


oxidoreductases

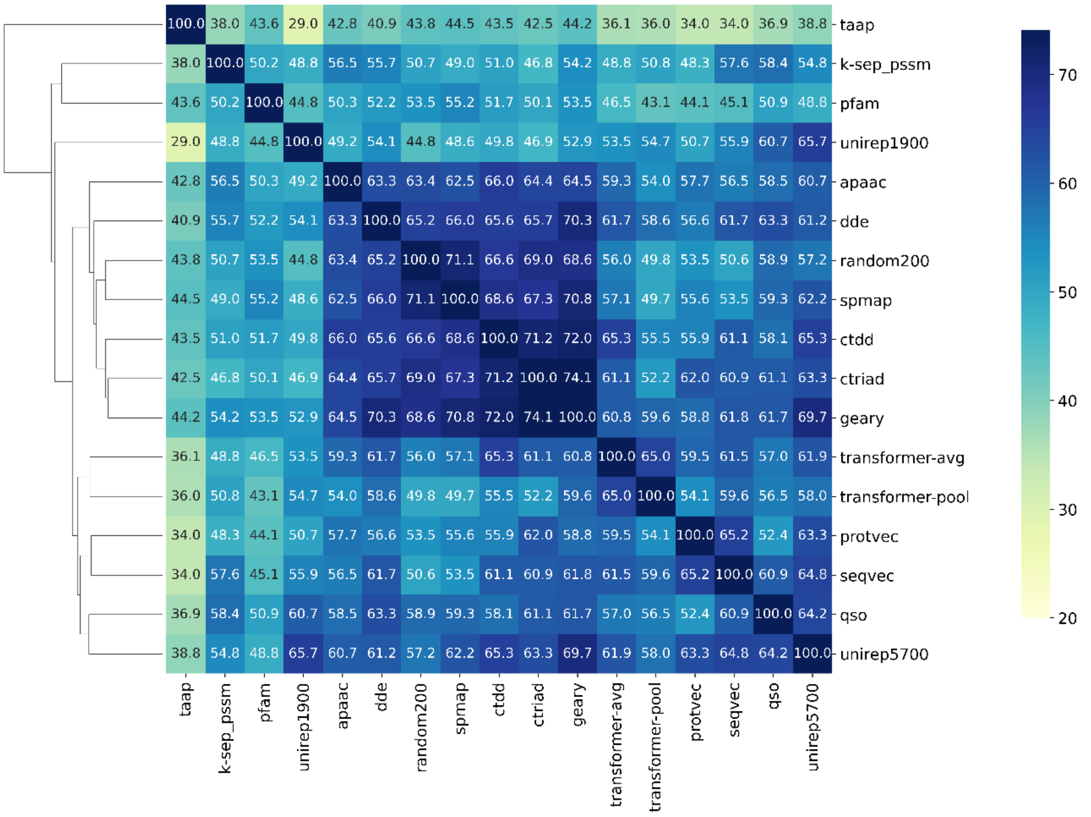

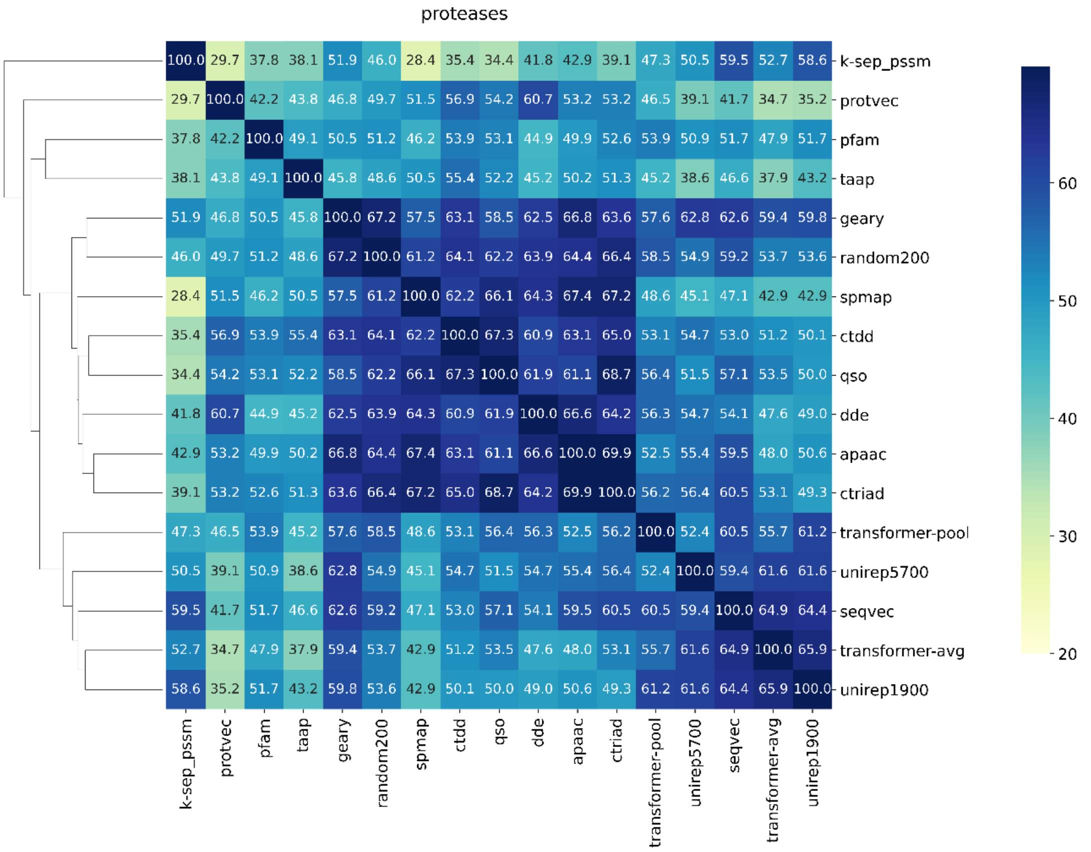
proteases
k-sep_pssm 100.0 29.7 37.8 38.1 51.9 46.0 28.4 35.4 34.4 41.8 42.9 39.1 47.3 50.5 59.5 52.7 58.6
protvec 29.7 100.0 42.2 43.8 46.8 49.7 51.5 56.9 54.2 60.7 53.2 53.2 46.5 39.1 41.7 34.7 35.2
pfam 37.8 42.2 100.0 49.1 50.5 51.2 46.2 53.9 53.1 44.9 49.9 52.6 53.9 50.9 51.7 47.9 51.7
taap 38.1 43.8 49.1 100.0 45.8 48.6 50.5 55.4 52.2 45.2 50.2 51.3 45.2 38.6 46.6 37.9 43.2
geary 51.9 46.8 50.5 45.8 100.0 67.2 57.5 63.1 58.5 62.5 66.8 63.6 57.6 62.8 62.6 59.4 59.8
random200 46.0 49.7 51.2 48.6 67.2 100.0 61.2 64.1 62.2 63.9 64.4 66.4 58.5 54.9 59.2 53.7 53.6
spmap 28.4 51.5 46.2 50.5 57.5 61.2 100.0 62.2 66.1 64.3 67.4 67.2 48.6 45.1 47.1 42.9 42.9
ctdd 35.4 56.9 53.9 55.4 63.1 64.1 62.2 100.0 67.3 60.9 63.1 65.0 53.1 54.7 53.0 51.2 50.1
qso 34.4 54.2 53.1 52.2 58.5 62.2 66.1 67.3 100.0 61.9 61.1 68.7 56.4 51.5 57.1 53.5 50.0
dde 41.8 60.7 44.9 45.2 62.5 63.9 64.3 60.9 61.9 100.0 66.6 64.2 56.3 54.7 54.1 47.6 49.0
apaac 42.9 53.2 49.9 50.2 66.8 64.4 67.4 63.1 61.1 66.6 100.0 69.9 52.5 55.4 59.5 48.0 50.6
ctriad 39.1 53.2 52.6 51.3 63.6 66.4 67.2 65.0 68.7 64.2 69.9 100.0 56.2 56.4 60.5 53.1 49.3
transformer-pool 47.3 46.5 53.9 45.2 57.6 58.5 48.6 53.1 56.4 56.3 52.5 56.2 100.0 52.4 60.5 55.7 61.2
unirep5700 50.5 39.1 50.9 38.6 62.8 54.9 45.1 54.7 51.5 54.7 55.4 56.4 52.4 100.0 59.4 61.6 61.6
seqvec 59.5 41.7 51.7 46.6 62.6 59.2 47.1 53.0 57.1 54.1 59.5 60.5 60.5 59.4 100.0 64.9 64.4
transformer-avg 52.7 34.7 47.9 37.9 59.4 53.7 42.9 51.2 53.5 47.6 48.0 53.1 55.7 61.6 64.9 100.0 65.9
unirep1900 58.6 35.2 51.7 43.2 59.8 53.6 42.9 50.1 50.0 49.0 50.6 49.3 61.2 61.6 64.4 65.9 100.0
k-sep_pssm protvec pfam taap geary random200 spmap ctdd qso dde apaac ctriad transformer-pool unirep5700 seqvec transformer-avg unirep1900
60
50
40
30
20

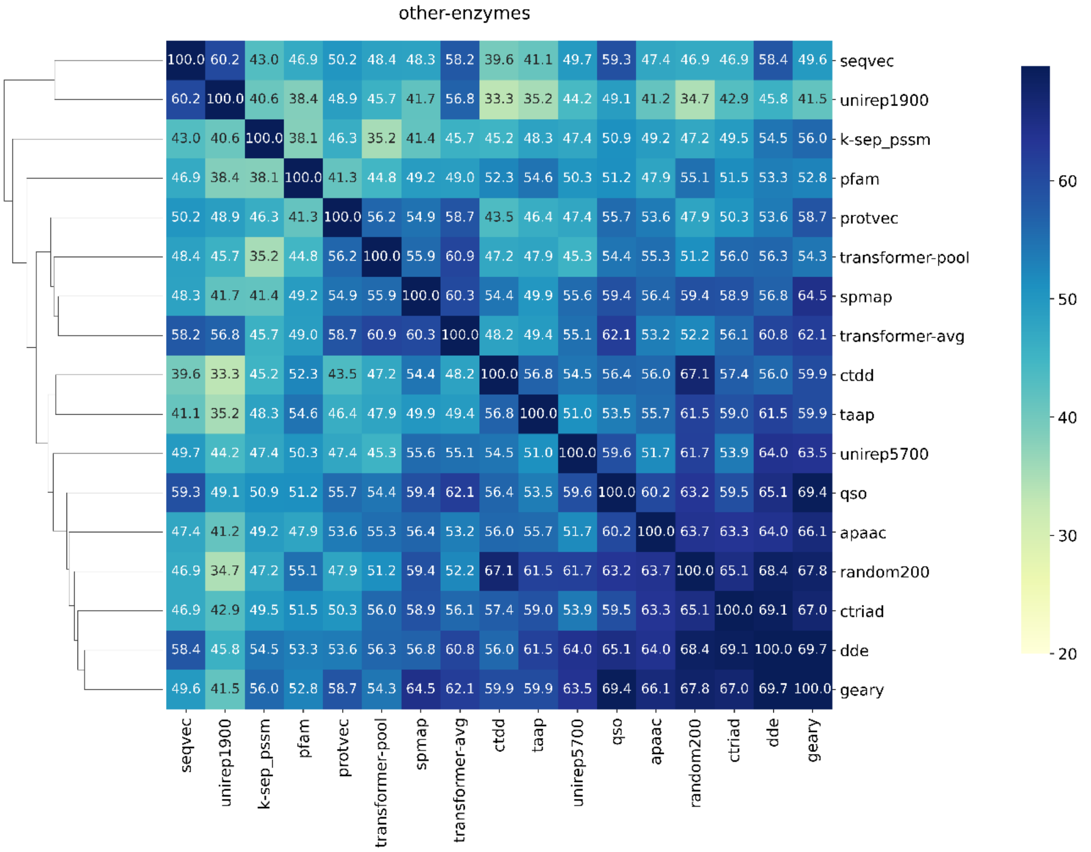
other-enzymes
seqvec 100.0 60.2 43.0 46.9 50.2 48.4 48.3 58.2 39.6 41.1 49.7 59.3 47.4 46.9 46.9 58.4 49.6
unirep1900 60.2 100.0 40.6 38.4 48.9 45.7 41.7 56.8 33.3 35.2 44.2 49.1 41.2 34.7 42.9 45.8 41.5
k-sep_pssm 43.0 40.6 100.0 38.1 46.3 35.2 41.4 45.7 45.2 48.3 47.4 50.9 49.2 47.2 49.5 54.5 56.0
pfam 46.9 38.4 38.1 100.0 41.3 44.8 49.2 49.0 52.3 54.6 50.3 51.2 47.9 55.1 51.5 53.3 52.8
protvec 50.2 48.9 46.3 41.3 100.0 56.2 54.9 58.7 43.5 46.4 47.4 55.7 53.6 47.9 50.3 53.6 58.7
transformer-pool 48.4 45.7 35.2 44.8 56.2 100.0 55.9 60.9 47.2 47.9 45.3 54.4 55.3 51.2 56.0 56.3 54.3
spmap 48.3 41.7 41.4 49.2 54.9 55.9 100.0 60.3 54.4 49.9 55.6 59.4 56.4 59.4 58.9 56.8 64.5
transformer-avg 58.2 56.8 45.7 49.0 58.7 60.9 60.3 100.0 48.2 49.4 55.1 62.1 53.2 52.2 56.1 60.8 62.1
ctdd 39.6 33.3 45.2 52.3 43.5 47.2 54.4 48.2 100.0 56.8 54.5 56.4 56.0 67.1 57.4 56.0 59.9
taap 41.1 35.2 48.3 54.6 46.4 47.9 49.9 49.4 56.8 100.0 51.0 53.5 55.7 61.5 59.0 61.5 59.9
unirep5700 49.7 44.2 47.4 50.3 47.4 45.3 55.6 55.1 54.5 51.0 100.0 59.6 51.7 61.7 53.9 64.0 63.5
qso 59.3 49.1 50.9 51.2 55.7 54.4 59.4 62.1 56.4 53.5 59.6 100.0 60.2 63.2 59.5 65.1 69.4
apaac 47.4 41.2 49.2 47.9 53.6 55.3 56.4 53.2 56.0 55.7 51.7 60.2 100.0 63.7 63.3 64.0 66.1
random200 46.9 34.7 47.2 55.1 47.9 51.2 59.4 52.2 67.1 61.5 61.7 63.2 63.7 100.0 65.1 68.4 67.8
ctriad 46.9 42.9 49.5 51.5 50.3 56.0 58.9 56.1 57.4 59.0 53.9 59.5 63.3 65.1 100.0 69.1 67.0
dde 58.4 45.8 54.5 53.3 53.6 56.3 56.8 60.8 56.0 61.5 64.0 65.1 64.0 68.4 69.1 100.0 69.7
geary 49.6 41.5 56.0 52.8 58.7 54.3 64.5 62.1 59.9 59.9 63.5 69.4 66.1 67.8 67.0 69.7 100.0
seqvec unirep1900 k-sep_pssm pfam protvec transformer-pool spmap transformer-avg ctdd taap unirep5700 qso apaac random200 ctriad dde geary
60
50
40
30
20

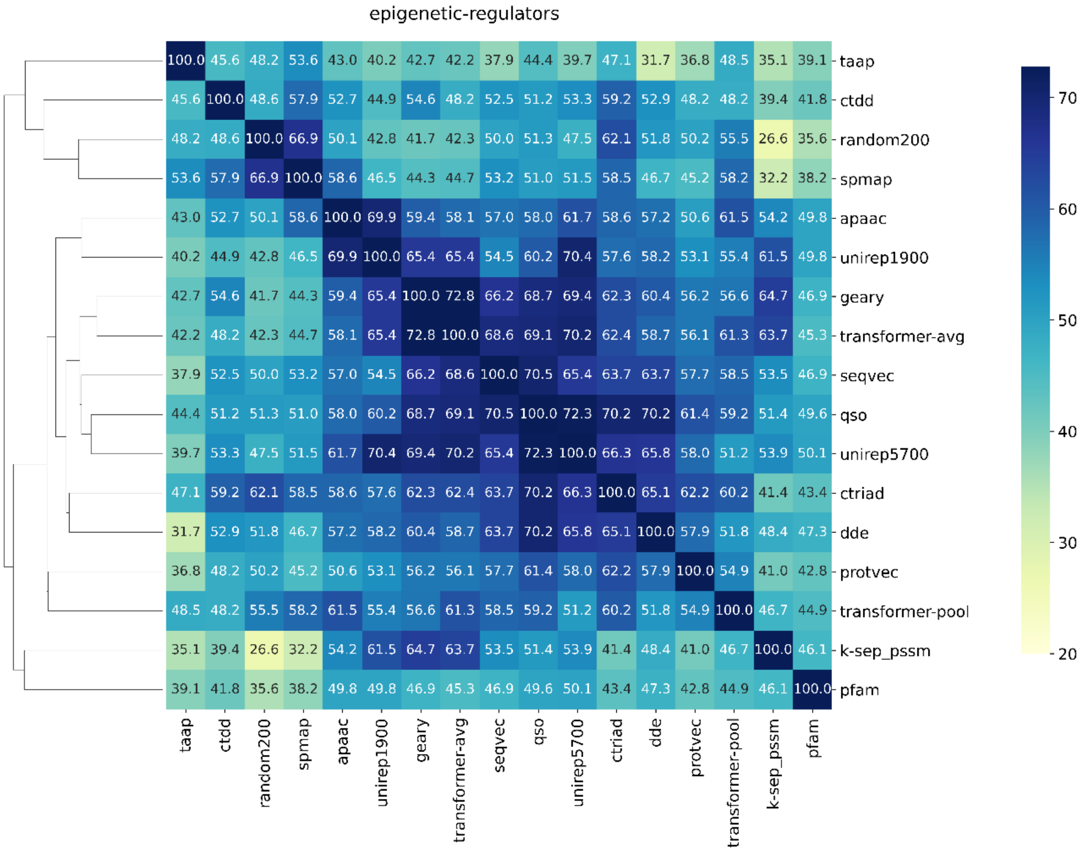
epigenetic-regulators
taap
ctdd
random200
spmap
apaac
unirep1900
geary
transformer-avg
seqvec
qso
unirep5700
ctriad
dde
protvec
transformer-pool
k-sep_pssm
pfam
70
60
50
40
30
20

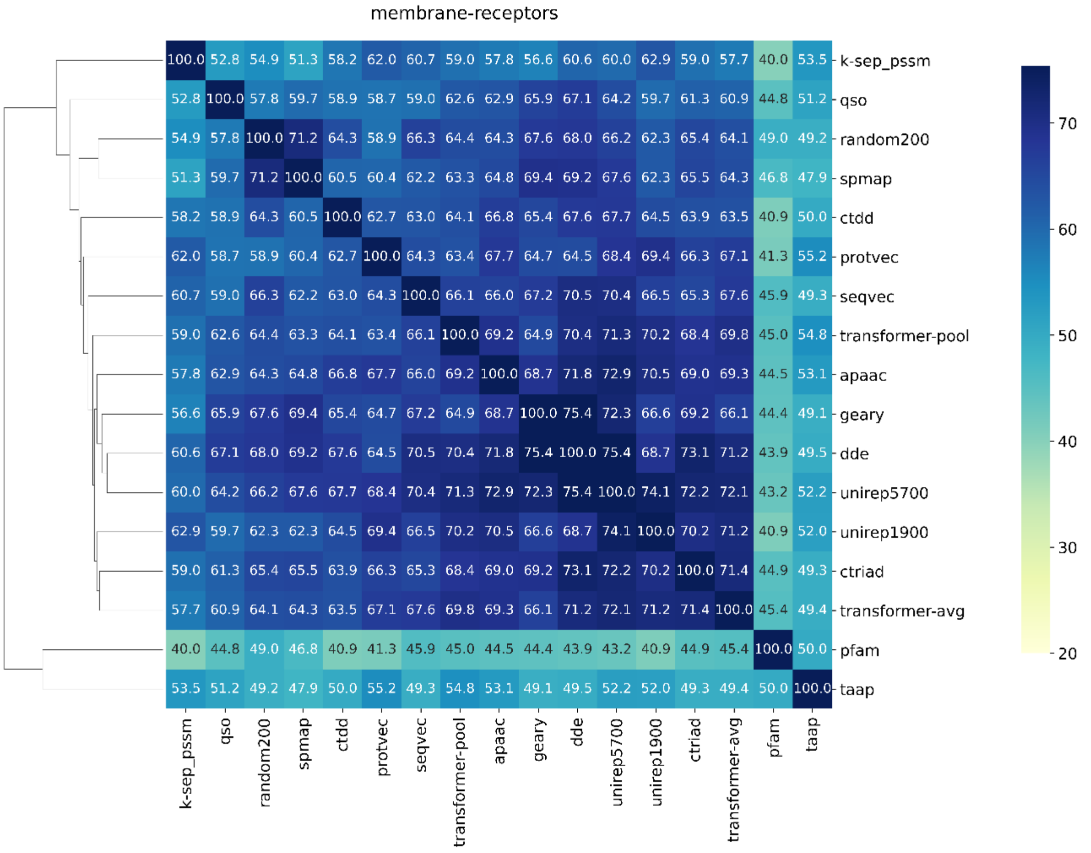
membrane-receptors
k-sep_pssm
qso
random200
spmap
ctdd
protvec
seqvec
transformer-pool
apaac
geary
dde
unirep5700
unirep1900
ctriad
transformer-avg
pfam
taap
70
60
50
40
30
20

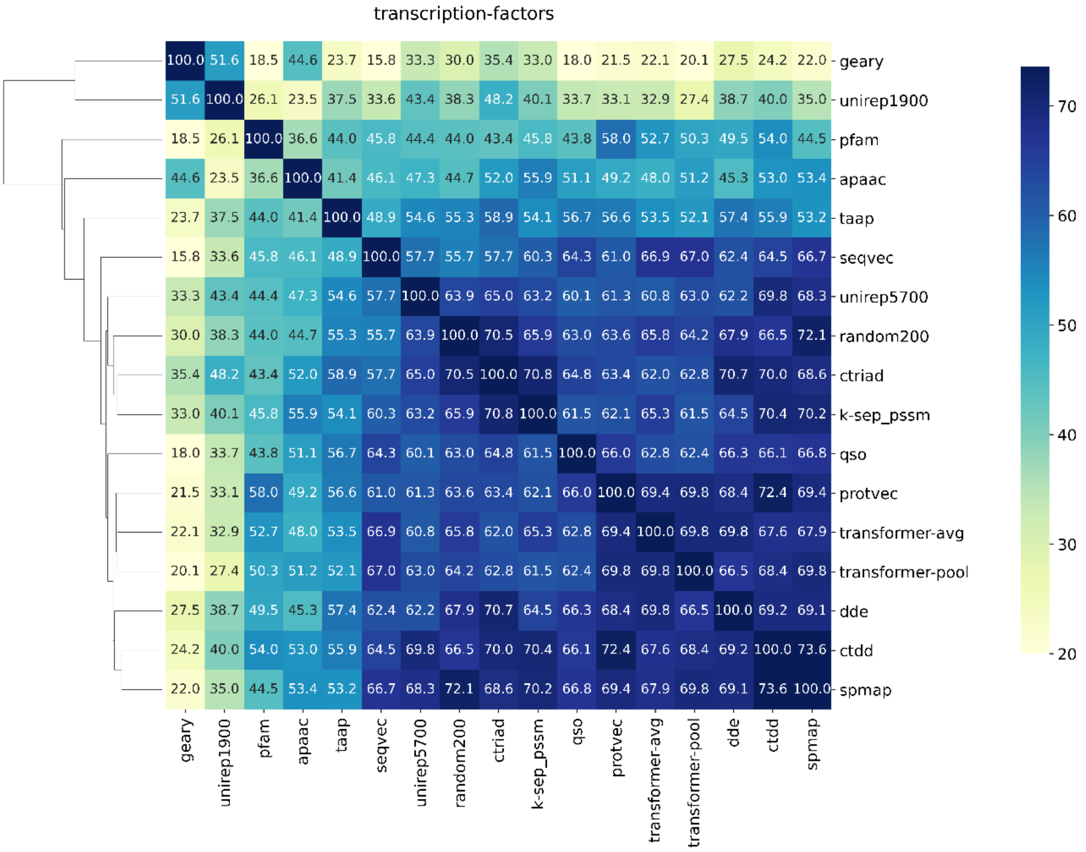


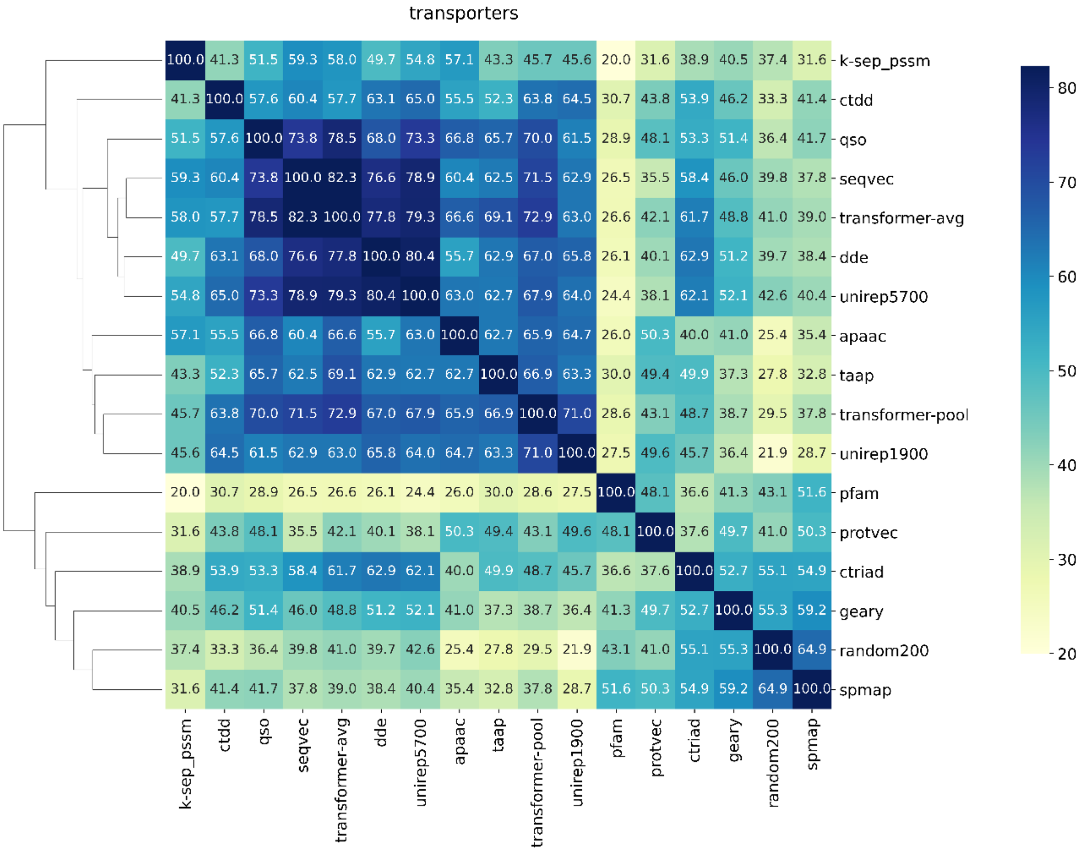


**Figure S3.** Clustered heatmaps of different protein representation approaches for protein families on **(a)** the random-split, **(b)** dissimilar-compound-split, and **(c)** fully-dissimilar-split datasets.
